# Integrase-mediated differentiation circuits improve evolutionary stability of burdensome and toxic functions in *E. coli*

**DOI:** 10.1101/614529

**Authors:** Rory. L Williams, Richard M. Murray

**Affiliations:** Department of Bioengineering, California Institute of Technology, Pasadena, CA; Department of Biomedical Engineering, UC Irvine; Department of Control and Dynamical Systems, California Institute of Technology

## Abstract

Advances in synthetic biology, bioengineering, and computation allow us to rapidly and reliably program cells with increasingly complex and useful functions. However, because the functions we engineer cells to perform are typically unnecessary for cellular survival and burdensome to cell growth, they can be rapidly lost due to the processes of mutation and natural selection. To improve the evolutionary stability of engineered functions in a general manner, we developed an integrase-recombination-based differentiation gene circuit in *Escherichia coli*. In this system, differentiated cells uniquely carry out burdensome or toxic engineered functions but have limited capacity to grow (terminal differentiation), preventing the propagation of selectively advantageous loss of function mutations that inevitably arise. To experimentally implement terminal differentiation, we co-opted the R6K plasmid system, using differentiation to simultaneously activate T7 RNAP-driven expression of arbitrary engineered functions, and inactivate expression of π protein (an essential factor for R6K plasmid replication), thereby allowing limitation of differentiated cell growth through antibiotic selection. We experimentally demonstrate terminal differentiation increases both duration and magnitude of high-burden T7 RNAP-driven expression, and that its evolutionary stability can be further improved with strategic redundancy. Using burdensome overexpression of a fluorescent protein as a model engineered function, our terminal differentiation circuit results in a ~2.8-fold (single-cassette) and ~4.2-fold (two-cassette) increase of total fluorescent protein produced compared to high-burden naive expression in which all cells inducibly express T7 RNAP. Finally, we demonstrate that differentiation can enable the expression of even toxic functions, a feat not achieved to our knowledge by any other strategy for addressing long-term evolutionary stability. Overall, this study provides an effective generalizable strategy for protecting engineered functions from evolutionary degradation.

## Introduction

As synthetic biology aims to engineer cells with the capacity to regulate and execute increasingly complex and burdensome functions, strategies which address the evolutionary instability of synthetic functions will only become more essential. It has long been observed that cell fitness negatively correlates with heterologous gene expression level,^1^ and increased burden results in a shorter evolutionary half-life of engineered functions.^2,3^ Efforts to improve evolutionary stability of engineered functions have taken a variety of forms,^4^ including the most straightforward goals of reducing mutation rate through sequence design^2,5^ or host-strain engineering^5–7^ to delay the appearance of mutations, or reducing burden constitutively^1,2,8^ or dynamically^9^ to mitigate their selective advantage. Additional strategies have improved evolutionary stability by various means of delaying or preventing the selection for mutations, including genomic integration of numerous copies,^10,11^ linking expression of a gene of interest (GOI) to an essential gene,^2,12^ addicting cells to the product of a metabolic pathway,^13^ or iteratively displacing populations of cells before mutational escape occurs.^14^ While these strategies have indeed made headway, they vary in their ability to be generalized to diverse functions, and in the effort required to do so. Furthermore, each of these strategies requires the GOI to be expressed by every cell in the population, fundamentally restricting their application to GOIs that are non-toxic and of low burden.

A homogenous population of cells all performing the same function is largely unique to the laboratory environment, and recent years have seen the merit of breaking with this paradigm by engineering consortia instead of individual strains.^15^ With inspiration from microbial communities, there have been numerous successful implementations of metabolic division of labor for production of biomolecules of interest.^16–18^ This strategy has numerous advantages, including reducing the number of genes and associated metabolic load in each specialized cell type, allowing independent optimization of separate pathways, and spatially separating potentially incompatible functions. While these benefits may be realized by combining in co-culture independently engineered strains or species, additional attractive properties become apparent with dynamically regulated division of labor encompassing both metabolic and reproductive functions within a single strain. The use of differentiation to coordinate such division of labor is a recurring strategy used by microorganisms,^19,20^ but it has not yet been fully explored for addressing evolutionary constraints of engineered functions. While natural systems using differentiation to facilitate metabolic and reproductive division of labor do so with beneficial or essential functions, here we apply this strategy to functions which are instead both unnecessary for host survival and burdensome to cell growth.

To adopt this strategy into a synthetic context, we develop a circuit architecture consisting of two cell types, with the first being specialized for the faithful replication of an encoded function in the absence of circuit burden, and the second—generated upon differentiation of the former—for the execution of the encoded function. We utilize integrase-mediated recombination to simultaneously activate T7 RNAP-driven expression of a burdensome engineered function, and inactivate the expression of π protein (an essential factor for R6K plasmid replication), thereby allowing limitation of differentiated cell growth through antibiotic selection. Maintaining T7 RNAP-driven expression of the function perfectly off prior to differentiation removes the selective pressure for cheaters in the progenitor population, and limitation of differentiated cell growth (terminal differentiation) prevents cheaters from exponentially expanding in the differentiated cell population. Here, we demonstrate computationally and experimentally that this terminal differentiation architecture is robust to burden and cheaters, can be improved with strategic redundancy, increases the evolutionary stability of burdensome engineered functions, and can even be applied to the production of toxic proteins.

## Results

### Terminal differentiation is a general strategy for addressing evolutionary stability

An ideal strategy for improving evolutionary stability is agnostic to the engineered function being expressed, allowing implementation without requiring extensive specialization for each use case. As we will demonstrate computationally, terminal differentiation fulfills this criteria by being robust to both burden level and to mutations which disrupt the function of interest. We develop this intuition with cartoons and deterministic modeling, and compare the performance of differentiation architectures to the benchmark case of engineered expression in which every cell in a homogenous population both encodes and expresses the function. In this strategy, which we designate naive expression (Figure 1A), the initial population of producer cells has a reduced growth rate due to the burden associated with the engineered function. Mutations which inactivate the expression of the burdensome engineered function, designated as burden mutations, give rise to *cheaters* which do not express the function and have a wild-type growth rate. Loss of expression of the function at the population level results from the combination of (1) selective pressure provided by the burdensome engineered function, and (2) the opportunity for cheaters to emerge by mutations during DNA replication and expand in the population through indefinite cell growth. Recognizing this, we considered strategies which would segregate these two features in order to prevent evolution from destroying engineered functions.

**Figure 1:**
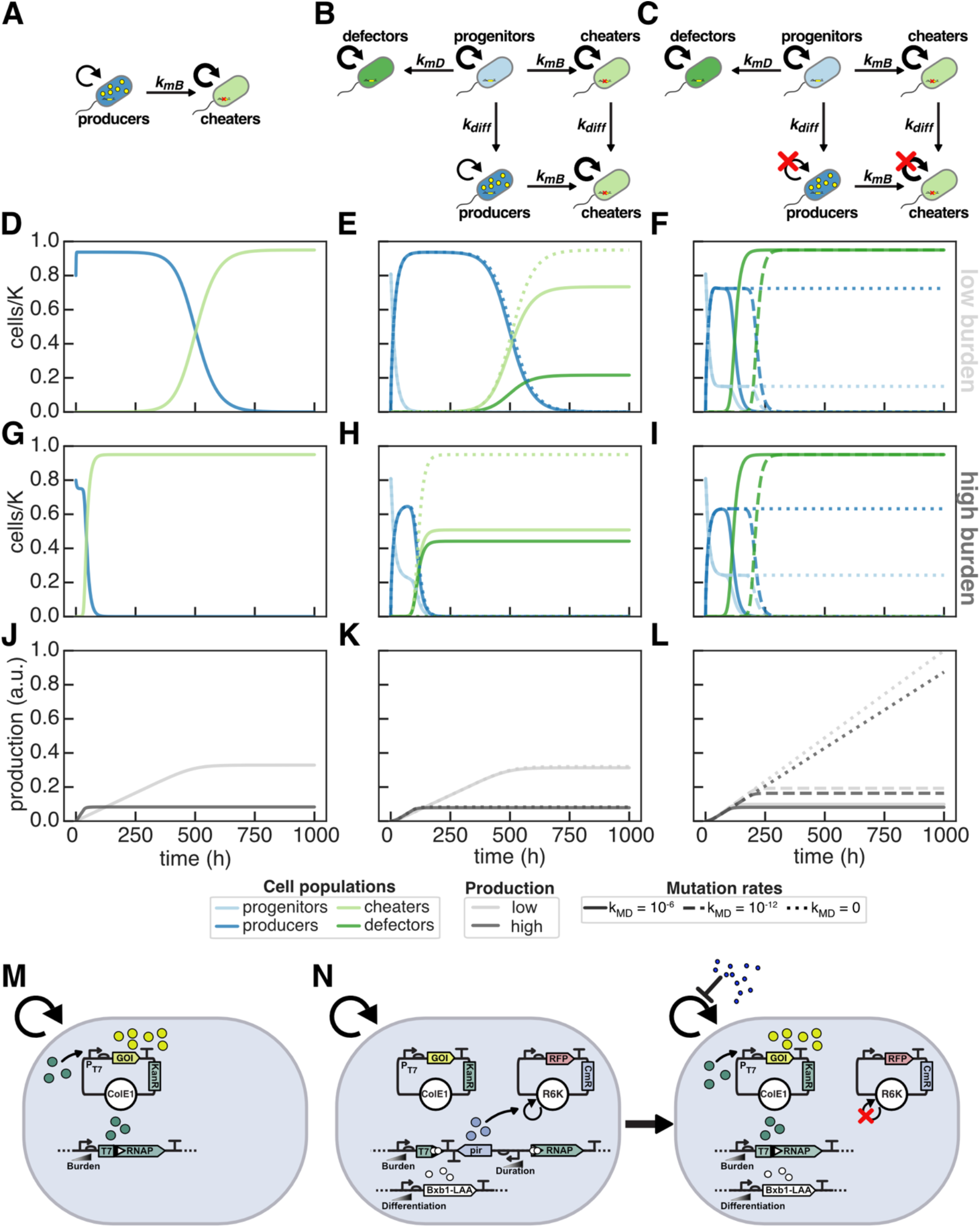
Strategies for burdensome expression. **(A-C)** Schematics for a naive expression **(A)**, differentiation-activated expression (differentiation) **(B)**, and differentiation-activated expression in which the number of cell divisions following differentiation is limited (terminal differentiation) **(C)**. **(D-L)** Deterministic ODE modeling of burdensome expression with circuits **(A-C)** growing with constant dilution and carrying capacity limited growth. *D* = 0.1 h^−1^, μ_N_ = 2 h^−1^, *k*_*MB*_ = 10^−6^ h^−1^, *n*_*div*_ = 4, *K* = 10^9^, initial cell population 8×10^8^ cells, and varying *k*_*MD*_ for differentiation and terminal differentiation (solid line: *k*_*MD*_ = 10^−6^; long-dashed: *k*_*MD*_ = 10^−12^ h^−1^; short-dashed: *k*_*MD*_ = 0). Burden due to production was 20% (μ_P_ = 1.6 h^−1^, D-F) and 80% (μ_P_ = 1.6 h^−1^, G-I). Abundance of subpopulations **(D-F, G-I)** and cumulative production **(J-L)** plotted over time. Producers (dark blue) express the function and have a reduced growth rate, while all other cells (cheaters: light-green; progenitors: light-blue; defectors: dark-green) have a normal growth rate. Experimental circuit design for **(M)** naive T7 RNAP-driven expression and **(N)** integrase-differentiation mediated activation of T7 RNAP which allows antibiotic selection against differentiated cells for terminal differentiation.

Terminal differentiation segregates the functions of DNA replication and expression of the burdensome engineered function by (1) activating expression through unidirectional differentiation, and (2) limiting growth of differentiated cells. With differentiation, a population of non-producing progenitor cells have a wild-type growth rate, and differentiate into producer cells which have a reduced growth rate (Figure 1B, differentiation). We refer to all cells that have incurred a burden mutation as cheaters, both progenitor cells that will differentiate into non-producers, and non-producer differentiated cells. While both exist, selective pressure for cheaters only exists in differentiated cells where the function is activated. To fully prevent the expansion of cheaters, we limit the growth of differentiated cells (Figure 1C, terminal differentiation).

The differentiation architecture, however, is susceptible to a new category of mutations occurring in progenitor cells which would destroy their capacity to differentiate. We refer to these cells that can no longer differentiate as *defectors.* While in the differentiation circuit without limited division, both cheaters and defectors have a selective advantage, only defectors have a selective advantage with terminal differentiation (Figure 1C). We model differentiation, terminal differentiation, and naive expression deterministically with systems of ordinary differential equations describing carrying capacity limited growth in a chemostat with constant dilution, production of an arbitrary protein by producer cells, varying production burden through adjusting the specific growth rate of producer cells, and varying burden mutation rates (generating cheaters). For differentiation, progenitor cells have a wild-type growth rate, the rates of differentiation and differentiation mutations (generating defectors) are varied, and the number of post-differentiation cell divisions is modeled explicitly for terminal differentiation.

Our intuition behind these architectures bears out in modeling, revealing terminal differentiation to be uniquely robust to cheaters. Following what has long been observed experimentally, with naive expression cheaters overtake the population more quickly with higher burden expression, resulting in a faster loss of function (Figure 1D, G, J; Figure S1D). While this is also true for differentiation without limited growth (Figure 1E, H, K; Figure S1D), the duration of function for terminal differentiation, strikingly, is unaffected by burden, as cheaters are never able to expand in the population (Figure 1 F, I, L; Figure S1D). While the performance of naive expression and the differentiation circuit without limited cell division are susceptible to cheaters and therefor sensitive to the rate of burden mutations, terminal differentiation is robust to this rate (Figure 1 D-E, G-H; Figure S1G). Further, decreasing the rate of the differentiation mutation (delaying emergence of defectors) uniquely improves longevity for terminal differentiation, and in the limit where the rate of this mutation is 0 we achieve indefinite function. While eliminating this mutation may not be possible, we are motivated to address this because it is the singular Achille’s heel of the terminal differentiation architecture. Because this circuit is agnostic to the specific function being expressed and robust to the rate of mutations which disrupt that function, improving the evolutionary stability of this architecture by reducing the probability of breaking the differentiation mechanism will improve the durability of any function regardless of burden.

### Development of an integrase-mediated terminal differentiation circuit

In order to experimentally implement the differentiation strategy, we required (1) burdensome expression to be fully off in the progenitor cell population, (2) irreversible and inducible activation of an arbitrary function through differentiation, and, in the case of terminal differentiation, (3) means of limiting the growth of differentiated cells. To accomplish this, we turned to bacteriophage serine integrases, a class of proteins capable of unidirectional DNA recombination between specific sequences of DNA.^21^ With strategic placement of integrase attachment sites on the genome, a single integrase-mediated recombination event can simultaneously activate and inactivate the expression of desired genes.^22–24^ In order to limit the capacity of differentiated cells to proliferate in the case of terminal differentiation, we take advantage of the reliance of R6K plasmid replication on the π-protein encoded by *pir*.^25^ We used the 3OC12-HSL (Las-AHL) inducible promoter P_LasAM_ and its cognate transcription factor LasR^AM^ to control the expression of π-protein, and placed its expression cassette such that the recombination event results in its excision (Figure 1N). The π-protein abundance and R6K plasmid copy number at the time of differentiation can therefore be tuned with Las-AHL, and as the R6K plasmid encodes the sole source of chloramphenicol resistance (CmR), the induction level of π-protein sets the limit on number of divisions possible upon differentiation when chloramphenicol selection is applied. In an initial evaluation of integrase differentiation, we demonstrated that the differentiation rate and R6K copy number could be controlled, and fraction of cells in the progenitor and differentiated state tuned with a combination of chloramphenicol selection and Las-AHL/salicylate induction (Figure S2).

Both to allow any arbitrary function to be expressed and to prevent leaky expression of the function in progenitor cells, we selected T7 RNAP, an orthogonal RNA polymerase broadly used in synthetic biology and bioproduction, ^26^ to be activated by this recombination. Importantly, T7 RNAP can then drive the expression of any protein or burdensome engineered function desired by the user. To allow the expression level and burden to be tuned, the evolved IPTG inducible promoter P_Tac_ and associated transcription factor LacI^AM^ were used to control the expression of T7 RNAP.^27^ Recombination-activatable T7 RNAP was integrated in a single copy on the *E. coli* genome, and a high copy Cole1-AmpR plasmid with T7 RNAP-driven sfGFP was used to report T7 RNAP expression. As initial designs which contained a ribosomal binding site (RBS) adjacent to an intact T7 RNAP coding sequence prior to recombination (Figure S3A-B) displayed leaky sfGFP expression in progenitor cells above negative control lacking T7 RNAP (Figure S3F), we relied on previous studies splitting T7 RNAP into functional domains to rationally choose a split site.^28^ With this strategy, there is no potential for leaky expression of functional T7 RNAP prior to differentiation, and the full-length coding sequence that is generated upon recombination contains a 17 amino acid insertion from the attL site and additional bases inserted to conserve the reading frame (Figure 1N, Figure S3C). In the absence of Bxb1 integrase, sfGFP production was equivalent to the control without T7 RNAP presence, with induction of Bxb1 allowing high level T7 RNAP-driven expression (Figure S3F).

In addition to the first three necessary criteria that have been fulfilled, the emergence of defectors ideally should be delayed in the terminal differentiation architecture by addressing the rate of differentiation mutations. From our initial deterministic modeling, we observed that decreasing the rate or probability of the differentiation mutation improves the duration of expression of the burdensome engineered function (regardless of burden level) of the terminal differentiation circuit by delaying the emergence of defectors. We reasoned that instead of aiming to decrease the rate of mutations by sequence design, increasing the number of independent mutations required to break the differentiation mechanism would yield more significant improvements. To this end, we envisioned an identical circuit design (Figure 1N) that instead has two T7 RNAP differentiation cassettes. Importantly the recombination of a single cassette would both activate the function and enable limiting the growth of differentiated cells. If a second identical cassette was integrated, recombination of both cassettes would be required to cease replication of the R6K plasmid and allow antibiotic selection mediated limitation of growth. Therefore a mutation preventing the recombination of one cassette would be sufficient to enable indefinite growth, giving opportunity for cheaters to emerge. However, if each differentiation cassette encoded a unique half of the π-protein, a single recombination event would ablate the expression of functional π-protein and with it the replication of the R6K plasmid. In this case, two independent mutations would be required to generate defectors, and cheaters would have no opportunity to gain a selective advantage. We therefore split the *pir* coding sequence and tagged the N- and C-terminal fragments with the N- and C terminal fragments of the Cfa intein,^29^ respectively, functionally screened for R6K plasmid replication, and identified a functional split site (Figure S4). Expression of the intein-tagged fragments allows R6K plasmid replication, and inactivation of either the 5’ fragment or 3’ fragment through integrase-mediated recombination results in loss of the R6K plasmid.

### Integrase-mediated differentiation circuits improve the evolutionary stability of burdensome T7 RNAP-driven functions

To assess the capacity of differentiation strategies to improve the evolutionary stability of burdensome functions, we performed long-term experiments with T7 RNAP-driven expression of a fluorescent protein, which here serves as a model burdensome engineered function. We compared the duration and total amount of production achieved in cells with one or two copies of inducible T7 RNAP (1x and 2x naive) to our single-cassette and two-cassette differentiation circuits (1x and 2x differentiation). Both the single-cassette differentiation and two-cassette differentiation strains have two copies of inducible integrase, and critically all components in the naive and differentiation circuits were genomically integrated, ensuring precise copy number control and preventing effects due to plasmid partitioning (Figure 2A–C). Experimental comparison of differentiation with terminal differentiation required only including chloramphenicol in the medium in the case of terminal differentiation, as without antibiotic present differentiated cells would grow unhindered after losing the R6K plasmid. Inducer and antibiotic conditions were uniform throughout the duration of the experiment, with the degree of burden tuned with IPTG (P_Tac_ T7 RNAP) and differentiation rate tuned with salicylate (P_SalTTC_ Bxb1-LAA). Experiments were ran for a total of 16 consecutive batch growths with 50x dilutions into a total volume of 300 μL every 8 hours, for a total of 128 hours and ~88 doublings.

**Figure 2:**
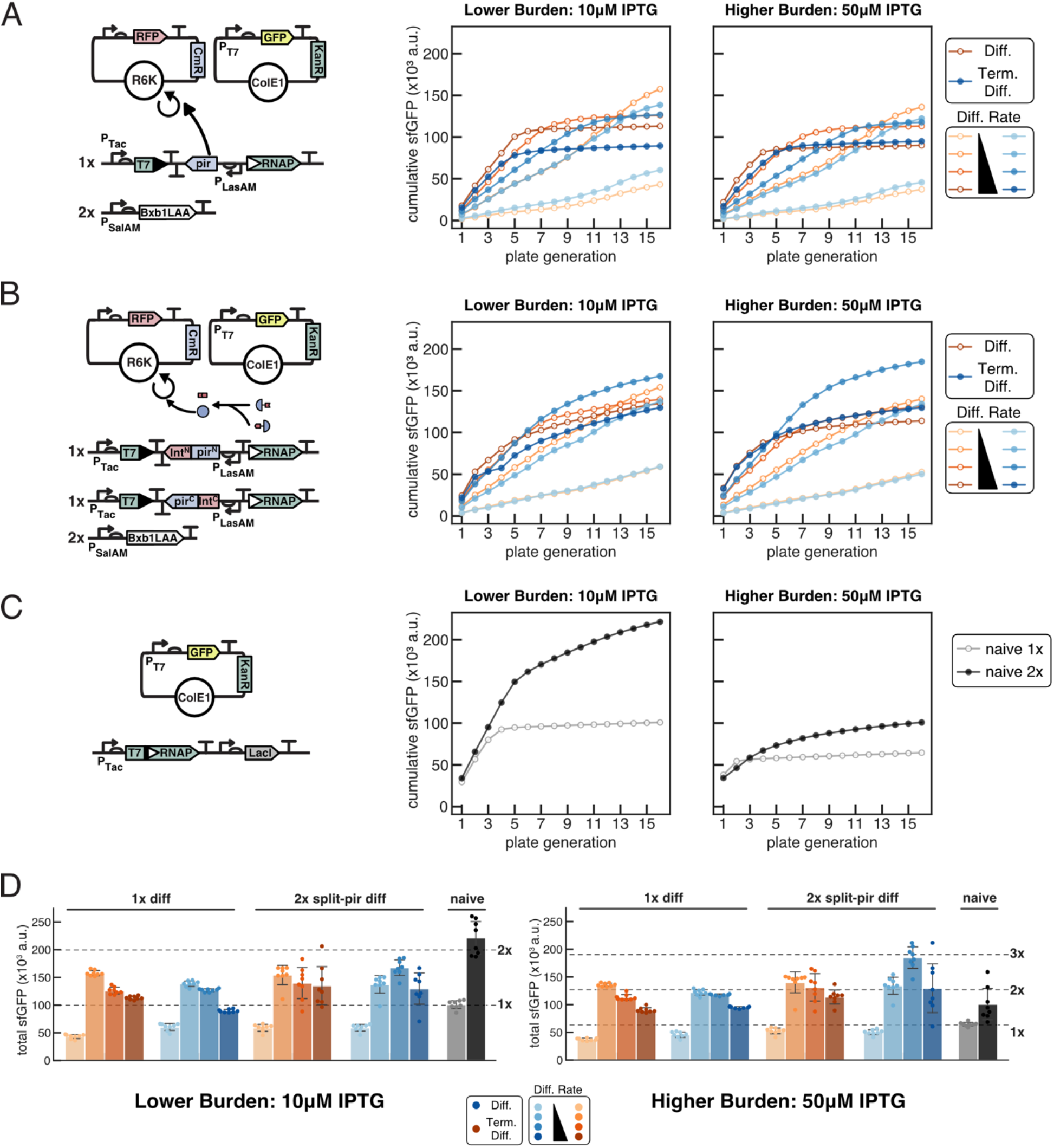
Assessing the evolutionary stability of burdensome T7 RNAP driven expression from a high copy ColE1 KanR plasmid. **(A-C)** 8 independent transformants were outgrown for 8 hours in LB media with appropriate antibiotics and inducers before 50x dilution into experimental conditions. Cells were grown in 96 well plates in 0.3 mL, diluted 50x every 8 h for 16 total growths, and 50 μL endpoint samples taken to measure OD700, sfGFP (485/515 nm), and mScarlet (565/595 nM) fluorescence in 384 well matriplates. Average cumulative sfGFP production plotted for each condition. **(A-B)** 1x differentiation and 2x differentiation. Each differentiation cassette additionally encodes NahR^AM^, LasR^AM^, and LacI^AM^ (Figure S17). Cells were co-transformed with ColE1-KanR-P_T7_ GFP and R6K-CmR-mScarlet, and plated on LB +kan/chlor/30 nM Las-AHL. Colonies were outgrown in LB +kan/chlor/10 nM Las-AHL before 50x dilution into experimental conditions in LB kan with chlor (blue, filled circles or without chlor (orange, open circles) with varying concentrations of salicylate (10, 15, 20, 30 μM) and IPTG (10, 50 μM). **(C)** Cells with one (naive 1x) or two (naive 2x) copies of genomically integrated T7 RNAP were transformed with ColE1-KanR-P_T7_ GFP, plated on LB + kan, and outgrown in LB + kan before dilution into experimental conditions. **(D)** Total cumulative production +/− SD after 16 growths for all strains in all conditions.

In these long-term experiments, we observe both the benefit of redundancy, and the superiority of terminal differentiation particularly with higher burden T7 RNAP-driven expression. Comparing 1x naive to 2x naive expression, we clearly see the benefit of redundancy in both the duration of production and total production achieved (Figure 2C–D). As each copy of inducible T7 RNAP is genomically integrated, each copy must independently mutate in order to fully disrupt its expression. The magnitude of this redundancy benefit is reduced in the higher burden case, however, with 2x naive yielding ~2.2x that of 1x naive in the lower burden case (~221000 +/− 31000 vs. ~101000 +/− 8000), and ~1.6x that of 1x naive in the higher burden case (~101000 +/− 30000 vs. ~65000 +/− 4000). For both 1x and 2x differentiation and terminal differentiation, the initial rate of GFP production increases with the differentiation rate (Figure 2A–B). However, higher differentiation rates also lead to more rapid cessation of production. These two counteracting features result in an intermediate differentiation rate yielding the most total sfGFP production (Figure 2D). With 1x differentiation, the terminal differentiation condition had a moderate negative effect on total production for both the lower burden (~1.56x 1x naive expression for diff. vs. ~1.37x for term. diff. with 15 μM sal/10 μM IPTG) and higher burden (~2.1x 1x naive expression for diff. vs. ~1.89x for term. diff. with 15 μM sal/10 μM IPTG) conditions. With 2x differentiation, terminal differentiation had minimal benefit in the lower burden condition (~1.53x 1x naive expression diff. w/ 15 μM sal/10 μM IPTG vs. ~1.66x term. diff. w/ 20 μM sal/10 μM IPTG), and a more significant benefit in the higher burden condition (~2.17x 1x naive expression for diff. w/ 15 μM sal/50 μM IPTG vs. ~2.86x for term. diff. w/ 20 μM sal/50 μM IPTG).

As expected, naive expression is more sensitive to burden level, and performs worse at higher burden in comparison to both differentiation and terminal differentiation (Figure 2D). In interpreting the effect of terminal differentiation for both 1x and 2x differentiation, we note that this has two opposing effects on the amount of production that will be achieved over the lifetime of circuit function. Terminal differentiation decreases output by limiting the growth of producer cells, but increases output by preventing the expansion of cheaters. Because these effects are opposing, one may dominate the other depending on the characteristic parameters. In the case of 1x differentiation, the latter postive effect cannot overcome the former negative effect at either low or high burden, and terminal differentiation performs worse. With 2x differentiation, the positive effect dominates, particularly so at higher burden. The critical difference between 1x and 2x differentiation which explains this is that while 1x requires only one differentiation mutation to yield defectors, 2x requires two mutations to yield defectors. This delay in the emergence of defectors provides additional time for the positive effect from suppression of cheaters to overcome the negative effect from limitation of producer growth.

### Terminal differentiation is uniquely robust to plasmid-based effects

As terminal differentiation is uniquely robust to cheaters, we speculated this robustness could extend to other sources, mutational or otherwise, which could impact expression. Because the GOI being expressed by T7 RNAP in these experiments is encoded on a high copy plasmid, plasmid mutations and copy number fluctuations (or even plasmid loss) can impact its expression without requiring mutation of the genomically encoded T7 RNAP. To investigate how such plasmid effects could influence each of our circuit architectures, we expanded the modeling framework previously discussed to reflect the experimental circuit design more accurately, model mutations stochastically, and incorporate plasmid-based effects. Specifically, we explicitly modeled the genotype of each cassette, incorporated integrase expression cassettes, modeled integrase mutations and differentiation cassette mutations separately, and modeled the differentiation rate of each cassette as being linearly dependent on the number of non-mutated integrase expression cassettes. Though plasmid copy number can certainly fluctuate widely, and fraction of plasmids with mutations vary due to random plasmid partitioning effects, for simplicity and tractability we considered plasmid mutation or plasmid loss to be binary, with a single stochastic event either mutating all plasmids, or resulting in complete plasmid loss. Furthermore, though experimentally antibiotic selection is used to ensure plasmid maintenance, communal resistance for certain antibiotics (e.g., β-lactams) can allow antibiotic sensitive cells to persist in the presence of antibiotic.^30^ To capture this effect, we modeled stochastic plasmid loss, varied the rate of Michaelis-Menten degradation of antibiotic by plasmid-containing cells, and assigned the growth rate of cells having lost the plasmid with a Heaviside function, where the growthrate is 0 if the antibiotic concentration is at or above the minimum inhibitory concentration (MIC), and is that of a non-producer if below the MIC. In simulating cells with one or two copies of inducible T7 RNAP in both the naive and differentiation cases, assumptions were required about the relative burden levels and production rates. In the case of differentiation, the producer growth rate (μ_P_) was the growth rate of a cell with one activated cassette of T7 RNAP, and the burden of the second copy produced a proportionate decrease in growth rate. For example, if a non-producer grows at rate 1 h^−1^ (μ_N_) and a cell with one cassette active grows at rate 0.5 h^−1^ (μ_P_), a cell with two cassettes active would grow at rate 0.25 h^−1^ (μ_N_(μ_P_/μ_N_)^2^). Production then was assumed to increase linearly with the decrease in growth rate. For full description of model implementation, see supplementary information.

From these simulations, we recapitulate several observations from the initial deterministic modeling of the general strategies, and from our experiments. For both differentiation and terminal differentiation with one and two copies, we see that lower differentiation rates result in slower production that lasts longer, high differentiation rates yields faster production that breaks more quickly, and intermediate rates strike a balance and achieve the most total production (Figure 3D–E). At low burden (higher μ_P_), terminal differentiation is counterproductive, but becomes beneficial as burden increases (Figure 3D–E). We also see that naive expression performs comparatively well at low burden relative to high burden. For 2x naive expression we model both the case where one cassette alone yields the growth rate μ_P_ and its corresponding production rate (2x*), and where the two cassettes together yield the growth rate μ_P_ (2x). As expected, at low burden and high differentiation rate, 1x differentiation without growth limitation approximates the performance of both 1x and 2x naive, and 2x differentiation without growth limitation approximates the performance of 2x* (Figure 3C–E). Further, the redundancy and mutational robustness provided with 2x differentiation improves performance relative to the one cassette case for both differentiation and terminal differentiation. Though we do not experimentally interrogate the impact of number of post-differentiation divisions in the case of terminal differentiation, we do so computationally, and while there is a benefit of increasing *n*_*div*_ at low burden, this effect disappears at higher burden (Figure S18).

**Figure 3:**
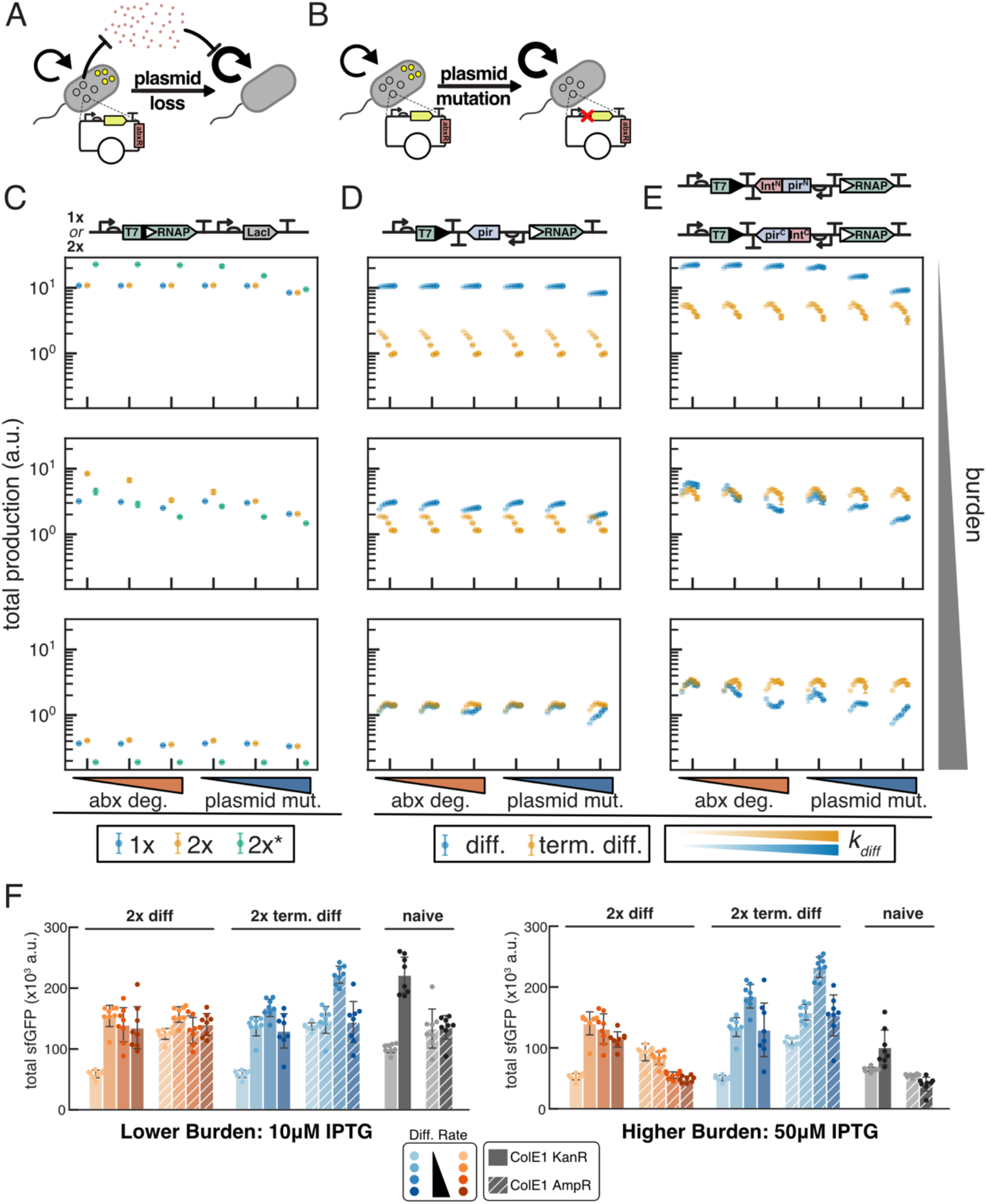
Terminal differentiation is robust to plasmid effects which hinder naive and differentiation architectures. **(A)** Schematic describing stochastic loss of a burdensome plasmid which encodes an antibiotic resistance gene that allows for communal degradation. **(B)** Schematic describing stochastic plasmid mutation which relieves burden but does not impact antibiotic resistance. **(C-E)** Stochastic simulations of burdensome production in 1x and 2x naive **(C)**, differentiation **(D)**, and terminal differentiation **(E)** architectures. Mean total production +/− SD of 8 stochastic simulations of 20 consecutive batch growths with 50x dilutions. μ_P_ = 2 h^−1^; 10, 50, 90 percent burden (increasing top to bottom); *K* = 10^9^ cells; *k*_*MB*_ = *k*_*MD*_ 10^−6^ h^−1^; *k*_*diff*_ = 0.2, 0.4, 0.6, 0.8, 1, 1.2 h^−1^; *n*_*div*_ = 4. Production rate and burdens/growth rates as described in supplementary model information. Naive 2x indicates two functional cassettes yields the indicated burden level 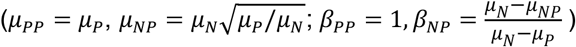 indicates one functional cassette yields that burden 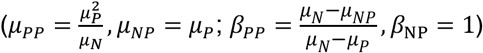. Simulations with antibiotic degradation are with plasmid loss rate *k*_*PL*_ = 10^−4^ h^−1^; 100 μg/mL antibiotic; minimum inhibitory concentration MIC = 1.1 μg/mL; and antibiotic degradation *V*_*max*_ = 0, 2.52 × 10^−6^, and 1.26 × 10^−5^ μg/cell/h (left to right increasing abx deg). Simulations with plasmid mutation were modeled with *k*_*PL*_ = 10^−8^, 10^−6^, 10^−4^ h^−1^ (increasing left to right), 0 μg/mL antibiotic, and *V*_*max*_ = 0. (F) Comparison of production achieved (mean +/− SD) with ColE1-KanR (solid) and ColE1 AmpR (diagonal stripes) variants of T7 RNAP-driven GFP for experiments described in Figure 2 and Figure S11).

Critically, this stochastic modeling reveals that the robustness of terminal differentiation to cheaters generalizes to plasmid-based mechanisms of cheating: both plasmid loss compensation by communal antibiotic degradation, and plasmid mutation. Incorporating stochastic plasmid loss in conjunction with antibiotic degradation and growth inhibition can negatively affect performance of the naive and differentiation architectures in a burden dependent manner. This effect is also much greater in the two cassette case (Figure 3C–E). Intuitively this makes sense, as even though the rate of plasmid loss is faster than of mutation, the difference in amount of time required to generate (1) a cell with no plasmid and (2) a cell with all copies of T7 RNAP mutated is much greater in the two cassette case. The performance of the terminal differentiation architecture, however, is robust to this plasmid instability. We see this both in the total production achieved (Figure 3D–E), and in tracking the population of cells which have lost the plasmid (Figure S22-S27). When there is no antibiotic degradation, we see no accumulation of cells lacking the plasmid for any circuit, but with a sufficiently high rate of antibiotic degradation, we see a transitory rise in the fraction of cells that have lost the plasmid for naive (Figure S19-S21) and differentiation architectures, but not for terminal differentiation (Figure S22-S27). This effect increases with higher rates of antibiotic degradation, is more pronounced at intermediate (50%) burden than very low (10%) or very high (90%) burden for naive expression, and, in the case of differentiation, is influenced by both burden level and differentiation rate. As these plasmid-deficient cells are dependent on communal antibiotic degradation, they do not completely take over the population, but instead are eventually displaced by mutated cells which retain the plasmid.

Similarly, when we neglect antibiotic degradation and instead consider plasmid mutation, we see that increasing the rate of plasmid mutation negatively impacts naive and differentiation architectures (Figure 3). Though this effect is similar to plasmid loss in that it affects the redundant architectures more significantly, it is different in that it reveals its effect at lower burden (10%), and because there is no dependence on communal antibiotic degradation, the accumulation of cells with plasmid mutations is not transitory (Figure S19-S27). Critically, the robustness of terminal differentiation circuits holds true when considering plasmid mutations, and we observe no accumulation of cells with mutated plasmids or any effect on production (Figure 3, Figure S22-S27).

While deliberately varying the rate of plasmid mutations experimentally is difficult, we may instead address plasmid instability through the choice of antibiotic resistance marker. Both to test this mechanism and to select any alternative selectable marker to use for the ColE1 plasmid, we used a plasmid identical to ColE1-KanR-P_T7_-GFP which instead had ampicillin resistance (ColE1-AmpR-P_T7_-GFP), and performed co-culture experiments of naive 1x and the parental strain JS006 transformed with these plasmids (Figure S6). As only naive 1x cells produce appreciable levels of GFP from these plasmids, we could observe that cells with only AmpR could allow cells with only KanR to grow in LB with both kanamycin and carbenicillin, while cells with only KanR did not allow cells with only AmpR to grow in the same condition. We therefore concluded that the choice of AmpR on the ColE1 plasmid would likely allow loss of expression through plasmid loss and shared antibiotic resistance, while the same mechanism would not hold with KanR.

Performing the same long-term evolution experiments described in Figure 2 with the single modification of using AmpR marker instead of KanR corroborates the intuition we gained from modeling. With both the lower and higher burden conditions, changing the selectable marker largely removes the benefit from redundancy with naive expression (Figure 3F, Figure S11). As well, while production is negatively affected by this change at higher burden for 2x differentiation without limited growth, production is actually higher with AmpR than KanR for 2x terminal differentiation at both burden levels (Figure 3F, Figure S11). While we cannot explain this performance benefit from our model, we speculate that the concentration of kanamycin used (50 μg/mL) can negatively affect expression even when KanR is expressed.^31^ This observation—that redundant terminal differentiation is positively affected, while redundant naive and non-terminal differentiation are negatively affected by plasmid instability—highlights the critical feature of terminal differentiation that the robustness to burden mutations extends generally to plasmid-based mechanisms of cheating.

### Differentiation enables the expression of toxic functions

As the terminal differentiation architecture is robust to the level of burden associated with the function of interest, this suggests that even toxic functions could be expressed and made evolutionarily stable. To test this, we aimed to demonstrate that the differentiation circuit we developed could allow the production of a toxic protein that will result in cell death: dnaseI. As progenitor cells do not produce any T7 RNAP, we reasoned that a T7 RNAP-driven dnaseI would not be expressed in the progenitor cells, allowing the encoded function to be replicated without toxicity or selective pressure for mutations. However, construction of a dnaseI expression plasmid identical to that of sfGFP yielded only mutated plasmids. Characterization of leaky expression from a P_T7_-GFP plasmid in the absence of T7 RNAP revealed fluorescence above background, explaining this inability to isolate functional plasmids (Figure S15). Incorporating two insulating terminators upstream of the T7 promoter mitigated leaky expression in the absence of T7 RNAP (Figure S15), and this insulation in conjunction with reducing the RBS strength allowed construction and isolation of a correctly sequenced dnaseI expression construct. While leak could have also been reduced by using the T7/lacO promoter and an additional source of LacI on the expression plasmid, this would not eliminate leaky expression in progenitor cells upon induction of differentiation and T7 RNAP. Highlighting the importance of preventing leaky expression of toxic functions, transformation of 1x differentiation and 2x differentiation cells with insulated dnaseI plasmid yielded ~600 cfu and ~1000 cfu, respectively, while naive 1x and naive 2x strains yielded 1 and 0 colonies, respectively, compared to >10^4^ cfu for both when transformed with ColE1-AmpR-P_T7_-GFP control (Figure 4, Table S1).

**Figure 4:**
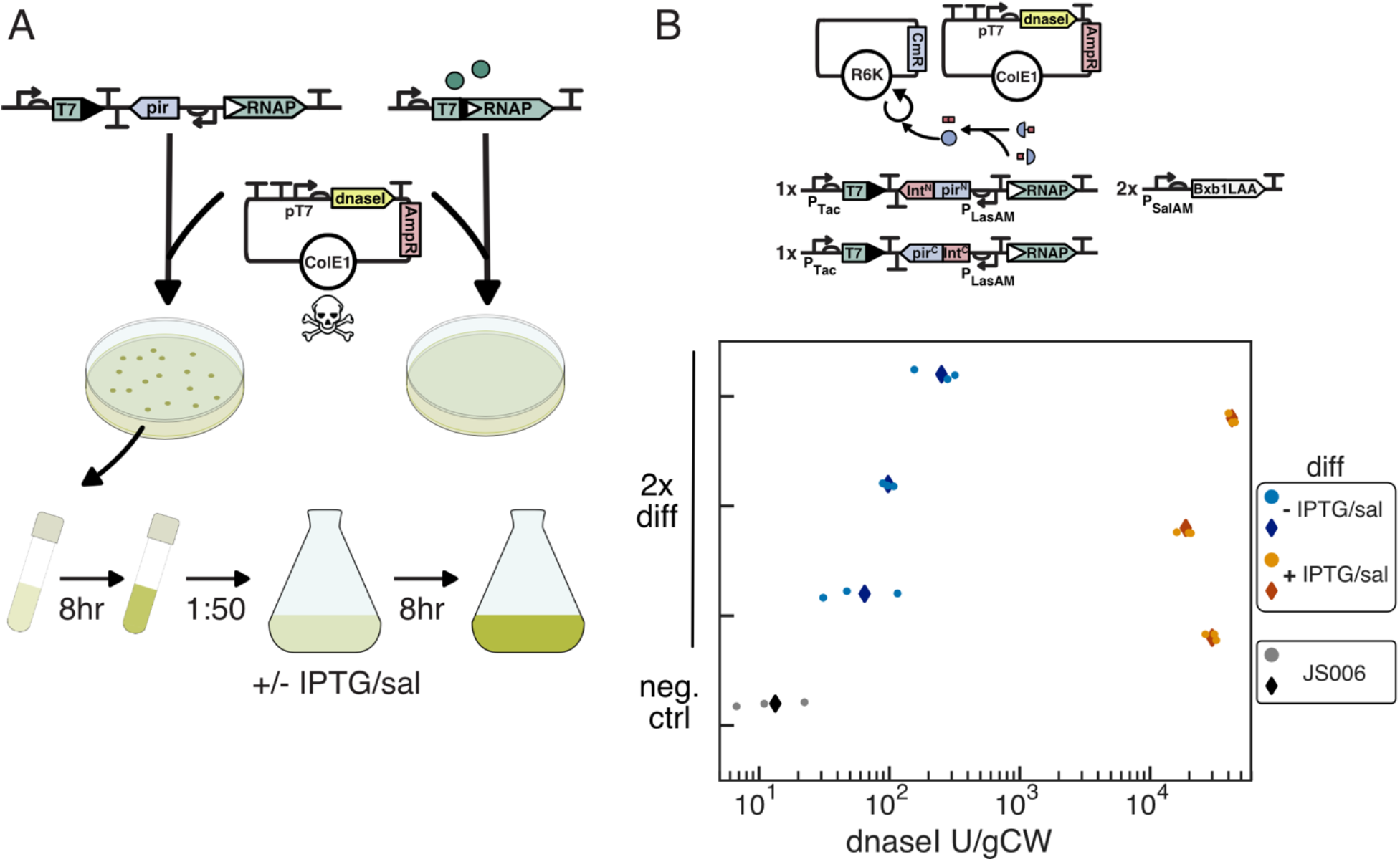
Production of functional dnaseI using integrase mediated differentiation. **(A)** Non-leaky T7 RNAP expression enables differentiation strains to replicate a plasmid encoding an insulated P_T7_ dnaseI cassette, while cells with leaky T7 RNAP cannot. **(B)** For assaying dnaseI production, 2x differentiation cells were co-transformed with R6K-CmR-empty and Cole1-AmpR-T13m-T12m-P_T7_-B0032-dnaseI-T7T, and plated on LB + carb/chlor/30 nM Las-AHL. After 8 h outgrowth in LB + carb/chlor/10 nM Las-AHL, cultures were diluted 1:50 into 25 mL media +/− 10 μM IPTG/20 μM sal. Pellets for experimental cultures and JS006 negative control were harvested after 8 h, and lysate assayed for dnaseI activity (Figure S16). Lysates from three independent cultures for each condition, and one JS006 culture were assayed in triplicate. Three measurements (circles) plotted with average (diamonds) for induced (orange/red), uninduced (light blue/blue), and JS006 control (grey/black).

Using the insulated T7 RNAP-driven dnaseI construct, we demonstrated that differentiation could enable expression of this toxic product. To assess the capacity of differentiation to enable functional dnaseI expression, we co-transformed the 2x differentiation strain with the insulated dnaseI expression plasmid, and an R6K-CmR-empty plasmid. After outgrowth without induction, cultures were diluted into 25 mL cultures with or without induction with 20 μM salicylate and 10 μM IPTG. After 8 hours of growth, un-induced cultures grew to cell densities equivalent to JS006 control (7.2-7.6 g wet weight per liter vs. 7.6 ctrl), while induced cultures reached lower cell densities (2-4.4 gCW/L). Diluting 1:50 into fresh media yielded similar densities for the uninduced cultures after 8 hours growth (7.2-7.6 gCW/L), while induced cultures had grown minimally after 8 h (OD600 < 0.05) and to a range of densities after 16 h total growth (2-6.8 gCW/L). The growth of the induced and uninduced cultures indicated that the dnaseI plasmid minimally affected growth when T7 RNAP is not expressed, and that 20 μM salicylate induction likely resulted in more complete differentiation in a large shaking culture in comparison to small volumes in 96 well microplates in previous experiments. Expression of functional dnaseI was quantified with an activity-based assay on lysate extracted from cell pellets (Figure S16), with activity measured equivalent to ~1.9-4.2×10^4^ U/gCW (~3.7×10^4^-1.9×10^5^ U/L) in the three independent induced cultures and ~65-250 U/gCW (~500-1800 U/L) in the uninduced cultures, compared to ~13 U/gCW (~100 U/L) for the JS006 negative control (Figure 4B). This yield of dnaseI is on the same order of magnitude as yields reported using T7 RNAP to drive the expression of recombinant dnaseI using the LacI repressible T7 promoter in Bl21(DE3)[pLysS] (1.5×10^4^ U/L) and Bl21(DE3)[pLysE] (7.5×10^4^ U/L).^32^

## Discussion

Here we have developed architectures for implementing differentiation and terminal differentiation in *E. coli* for the expression of burdensome T7 RNAP-driven functions. Importantly, in our circuit design progenitor cells do not have an intact coding sequence for T7 RNAP, completely eliminating leaky T7 RNAP expression prior to differentiation. We computationally demonstrated that limiting the growth of differentiated cells with terminal differentiation provides robustness to burden level and cheaters, and that reducing the rate of differentiation mutations delays the emergence of defectors, thereby improving the evolutionary stability of the terminal differentiation architecture and with it any arbitrary function.

Experimentally, we developed a differentiation-activated T7 RNAP architecture in which all circuit components that could mutate to disrupt the process of differentiation or expression of T7 RNAP were integrated on the genome, both ensuring exact copy number control and preventing plasmid partitioning effects from accelerating circuit failure.^33^ With the goal of delaying the emergence of defectors and thereby increasing the longevity of the terminal differentiation architecture, we developed a novel split-π protein system using split-inteins.^29^ In long-term experiments with repeated dilutions of independent cultures, we compared the performance of 1x and 2x naive T7 RNAP-driven expression to 1x and 2x differentiation and terminal differentiation. We demonstrated that the rate, duration, and total amount of production could be tuned by varying the differentiation rate, with lower differentiation rates enabling longer duration expression but at a slower rate. Differentiation was particularly beneficial in comparison to naive expression with higher burden as expected from modeling, and the redundancy and robustness to differentiation mutations provided by incorporating the split-π protein design into the terminal differentiation architecture proved effective. Though here we demonstrate the effectiveness of terminal differentiation and the stability benefit gained from requiring two instead of one mutations to generate defectors, this can be viewed as demonstration of the power that redundancy can provide in synthetic biology. In considering the scaling of this strategy to longer times and larger population sizes, we expect that higher degree of redundancy will be necessary. Scaling this specific architecture through further splitting of the π protein may be infeasible, but the tool-kit of synthetic biology certainly has means of allowing this strategy to scale further, through the inactivation of essential genes, activation of toxins, or otherwise.

We demonstrated both computationally and experimentally that effects due to instability of T7 RNAP-driven expression plasmid and communal antibiotic resistance can negatively affect the performance of naive expression and differentiation, but that terminal differentiation circuits are robust to this effect. We further showed computationally that the robustness of terminal differentiation circuits to cheaters extends to the general case of plasmid mutations which disrupt the function of interest. Though genomic integration of functions is more time consuming and cumbersome than plasmid transformation, plasmid instability and considerations of cost for use of antibiotics and inducers in large cultures have often made genomic integration of constitutively expressed functions the preferred method for bioproduction in industry.^10,34^ However, because with terminal differentiation effects of plasmid instability can be substantially or entirely mitigated, we can potentially get the stability benefits of genomic integration with the ease of plasmid transformation.

Finally, because there should be no limit on the degree of burden or toxicity of a function expressed with our differentiation system (so long as the toxicity is limited to the cells expressing the function), as a proof of concept we demonstrated that differentiation could enable the expression of functional dnaseI. In the course of this demonstration, we discovered that in the absence of leaky expression of T7 RNAP, non-T7 RNAP sources of leak could prevent isolation of correctly assembled dnaseI expression plasmids. While we mitigated this problem through the incorporation of insulating terminators to prevent transcriptional read-through from upstream of the T7 promoter, reducing the strength of the RBS was still required to isolate correctly sequenced plasmid. Improving this insulation and/or reducing any leaky expression that may be coming directly from the T7 promoter through directed evolution efforts may prove beneficial. Addressing this would allow the use of higher strength RBS sequences without concern for leak, thereby enabling improved yields. While the expression of toxic or highly burdensome products has long been of interest in bioproduction and synthetic biology, and effective strategies have been implemented to accomplish this,^35^ to our knowledge all existing strategies only work for single use batch culture inductions. The critical difference with our strategy of terminal differentiation is that progenitor cells continuously differentiate to replenish the population of cells expressing the toxic function, thereby allowing a toxic product to be produced continuously.

We envision this strategy to be readily applied to the expression of burdensome and toxic proteins or metabolic pathways with little to know modification of the system, simply by transformation of plasmid encoding the desired T7 RNAP-driven function. With the performance of the current redundant terminal differentiation architecture, expression of the GOI can continue for 10-16+ plate generations (~55-88 doublings) depending on differentiation rate. This equates to the number of doublings occurring in ~100-150+ hours of continuous culture with a dilution rate of 0.4 h^−1^, suggesting this could enable continuous bioproduction. Furthermore, in contrast to naive expression where cell growth must be considered in optimizing expression, with terminal differentiation this optimization can be done with production yield as the sole factor, potentially enabling higher per cell production rates. This feature naturally motivates the application of metabolic engineering strategies. While tools like flux balance analysis can inform genetic modification of strains to improve the yield of valuable chemicals,^36^ these strategies naturally must be concerned with the growth of the organism. However, with terminal differentiation, genomic and metabolic knobs could be tuned to maximize yield without regard for the long-term viability of the cells. CRISPR/Cas systems have been demonstrated to allow activation and repression in *E. coli*, have been applied in metabolic engineering efforts, and could be co-opted in this context.^37,38^ We are excited both by these engineering opportunities that are enabled by the ability to neglect cell growth in the producer cell population, and for future development of terminal differentiation to further extend the evolutionary stability of arbitrary functions.

## Materials and Methods

### Strains and constructs

The wild-type *E. coli* strain JS006 was the base strain for the construction of all differentiation and naive circuit strains. Constructs were assembled with a combination of Golden Gate and Gibson assembly using 3G,^39^ and were integrated into the *E. coli* genome using clonetegration.^40^ Because the R6K origin used for propagation of pOSIP plasmids from the clonetegration method of genomic integration is the same origin in our differentiation architecture R6K plasmid, we PCR amplified pOSIP backbones in two pieces, removing the R6K origin, for use in Gibson assemblies with desired inserts. For Gibson assembly with linear pOSIP pieces, POS1 and POSX were used as terminal adapters instead of UNS1 and UNSX. The 1x naive strain and 1x differentiation strain was constructed by integration at the P21 (T) landing site, and the 2x naive and 2x split-π protein differentiation strains by additional integration at the HK022 (H) landing site. 1x differentiation and 2x differentiation strains were integrated two additional times with the inducible Bxb1-LAA expression construct at the primary and secondary phage 186 (O) landing sites (Figure S17). Following transformation, integrations were checked via colony PCR with the pOSIP p4 primary corresponding to the landing site and a reverse primer common to all pOSIP plasmids (5’ ATTACTCAACAGGTAAGGCG 3’). Fidelity of integrations was checked with a combination of sequencing and functional screening prior to transformation with pE-FLP to excise the antibiotic resistance cassette and integration module, and integration of subsequent constructs. Final strains for 1x and 2x naive, 1x differentiation 2x Bxb1, and 2x split-π differentiaton 2x Bxb1 were whole-genome sequenced with minION using the Rapid Barcoding Kit (Nanopore SQK-RBK004) for verification. Reads were assembled with Flye (https://github.com/fenderglass/Flye/) and mapped to reference genomes containing intended genomic insertions in Geneious.

Modified MoClo^41^ compatible parts for T7 RNAP, integrase attachment sites, and terminators were generated with standard molecular biology techniques (PCR, Gibson, oligo annealing and phosphorylation), and modified UNS adapters used for construction of polycistronic or inverted transcriptional units. The R6K-CmR backbone was constructed with Golden Gate using an R6K origin amplified from the pOSIP plasmids. Sequences for Bxb1 integrase attachment sites attB and attP were obtained from Ghosh.^42^ NahR^AM,^ LasR ^AM,^ and LacI ^AM,^, and their corresponding evolved promoters P_SalTTC_, P_LasAM_, and P_Tac_ were provided by Adam Meyer.^27^ The CIDAR MoClo Parts Kit which includes various promoter, RBS, CDS, and terminator parts used in the constructs described were provided by Douglas Densmore (Addgene kit 1000000059).

### Differentiation experiments

Chemically competent cells were prepared from the naive and differentiation strains grown in LB without selection, with differentiation strains induced with 30 nM Las-AHL to allow π protein expression for R6K plasmid replication. 1x and 2x naive strains were transformed with ColE1-KanR-P_T7_-GFP or ColE1-AmpR-P_T7_-GFP and plated on LB with 50 μg/mL kanamycin or 100 μg/mL carbenicillin, respectively. Differentiation strains were co-transformed with R6K-CmR-mScarletI and ColE1-KanR-P_T7_-GFP or ColE1-AmpR-P_T7_-GFP, recovered in SOC with 30 nM Las-AHL, and plated on LB agar with 34 μg/mL chloramphenicol, 30 nM Las-AHL, and 50 μg/mL kanamycin or 100 μg/mL carbenicillin, respectively. Eight independent colonies were picked from each transformation, and grown at 37C in 300 μL LB in 96 square deep well plates (Southern Labware SKU# 502062) sealed with breathable film (Diversified Biotech BERM-2000) for 8 hours. Naive strains were grown in LB with appropriate antibiotic, and differentiation strains grown in LB with chlor and carb or kan with 10 nM Las-AHL. Following outgrowth, cells were diluted 1:50 into experimental conditions. Cells were diluted every 8 hours for sixteen total growths in constant antibiotic and induction conditions, and sfGFP (485/515 nm), mScarlet (565/595 nm), and OD700 measured by taking 50 μL aliquots of endpoint culture and measuring in 384 well black wall clear bottom microplates (Thermo Scientific 142761). Average of two reads for each measurement in each well were used.

### dnaseI expression and quantification

Chemically competent 1x and 2x naive strains, and 1x and 2x differentiation strains were transformed with 10 ng of Cole1-AmpR-P_T7_-GFP or 10 ng Cole1-AmpR with insulated P_T7_ dnaseI, and all or 10 percent plated on LB carb. Plates with more than 1000 colonies on the 10 percent plate were reported as >10^4^ cfu. For dnaseI expression experiments, 2x split-π differentiation cells were co-transformed with an empty R6K-CmR plasmid and the insulated ColE1-AmpR P_T7_ dnaseI expression plasmid, recovered in SOC with 30 nM Las-AHL, and plated on LB agar with carb/chlor/30 nM Las-AHL. Three independent colonies were inoculated into 3 mL LB cultures with carb/chlor/10 nM Las-AHL, outgrown for 8 hours at 37C, and diluted 1:50 into 25 mL media with or without 20 μM salicylate and 10 μM IPTG to induce differentiation and T7 RNAP expression. After 8 hours of growth, cultures were diluted 1:50 into the same conditions, and the remaining culture harvested. Wet weight of pellets after washing with PBS was recorded before storing at −20C. JS006 parental strain without the dnaseI expression plasmid was grown similarly in LB without antibiotics and inducers for negative control. The second growth of uninduced cultures was harvested after 8 hours as before, and induced cultures allowed to grow for an additional 8 hours as minimal growth was observed after the initial growth.

Pellets were lysed via sonication of a 33 percent (w/v) cell suspension in 10 mM Tris pH 7.5/2 mM CaCl2 with protease inhibitor (Roche, 11836170001), cleared with centrifugation at 4C, and supernatant collected and kept on ice before assaying dnaseI activity. Buffers used for assay were as described in Kunitz,^43^ though to allow simultaneous measurement of many samples and to avoid problems we observed with background absorbance in crude cell lysate when performing the Kunitz assay, we developed a fluorescence-based assay similar to Vogel and Frantz.^44^ Briefly, dnaseI assay buffer was prepared by diluting SYBR Safe (Invitrogen, S33102) 1:1000 into a solution of 100 mM sodium acetate/5 mM magnesium sulfate with 26.3 μg/mL calf thymus DNA (Sigma D1501). Assay plate was prepared by aliquoting 190 μL dnaseI assay buffer into 96 round-well clear bottom plates and equilibrating in the dark at 25C. Standards were prepared by adding various amounts of dnaseI (Invitrogen AM2222) to JS006 lysate diluted 1:10 in 0.85 percent NaCl. Samples to assay were diluted 1:10 or 1:50 in 0.85 percent NaCl, and 10 μL of sample or standard pipetted with a multi-channel pipette into triplicate wells immediately before assay. The final amount of DNA per well was 5 μg. After shaking briefly fluorescence (487/528 nM) was measured every minute for 2 hours at 25C. Fluorescence fold-change over the course of the two hour assay was used in fitting a standard curve (Fig S16), and dnaseI activity calculated from appropriate dilutions.

### Model simulations

Simulations were run in Python using systems of ODEs using Euler’s method with a time step of 0.01 hours. This custom ODE solver was used to allow choice of modeling mutations deterministically using first-order rate constants for mutations, or stochastically by drawing from a binomial distribution. Production was modeled as being proportional to the ratio of specific growth rate (actual growth rate after accounting for effect due to carrying capacity) to maximum growth rate for the specific cell type, and production rate was 1 a.u. per cell per hour for all simulations regardless of burden. For terminal differentiation, the number of divisions allowed (*n*) is explicitly modeled. Immediately after differentiation, a cell has divided *i* = 0 times. Division of a cell for which *i* = 0 generate two cells with *i* = 1. Cells for which *i* = *n*, instead of dividing, then die at the same rate. Simulations in Figure 1 and Figure S1 were modeled deterministically with constant dilution and carrying capacity limited growth for 1000 hours of simulated time. For Figure 3 and Figure S18-S27, simulations were of batch growths with 1:50 dilutions every 8 hours for 20 total growths. Growth, differentiation, and production were modeled deterministically, while all mutations were modeled stochastically by drawing from a binomial distribution. 8 replicate simulations were run for each condition. For full description of model implementation, see supplementary information.

## Acknowledgements

The authors would like to thank the Sim Lab at the University of California Irvine for use of lab space and equipment; Andrey Shur for his help with minION sequencing; Andy Halleran, Anandh Swaminathan, and Andrey Shur for productive conversations; and Prof. Chang Liu, John Marken, and Gordon Rix for providing comments on the manuscript. NahR^AM,^, LasR^AM,^ and LacI^AM,^, and their corresponding evolved promoters P_SalTTC_, P_LasAM_, and P_Tac_ were provided by Adam Meyer.^27^ The CIDAR MoClo Parts Kit which includes various promoter, RBS, CDS, and terminator parts used in the constructs described were provided by Douglas Densmore (Addgene kit 1000000059). This research was supported by the Army Research Office (ARO) through grant W911NF-19-2-0026, and by the Army Research Lab/Institute for Collaborative Biotechnologies (ARL/ICB) through grant W911NF-09-D-0001. The content of the information on this page does not necessarily reflect the position or the policy of the Government, and no official endorsement should be inferred.

