## Supplementary Information for "Integrase-mediated differentiation circuits improve evolutionary stability of burdensome and toxic functions in *E. coli*"



| | |
|---|---|
| Figure S1 | Deterministic modeling of strategies for expression of burdensome functions |
| Figure S2 | Implementation of a tunable integrase-mediated differentiation circuit |
| Figure S3 | Integrase mediated activation allows leak-free expression of T7 RNAP |
| Figure S4 | Integrase inactivation of π protein expression ablates R6K plasmid replication |
| Figure S5 | Characterization of burden level of naive T7 RNAP driven GFP |
| Figure S6 | Assessing communal antibiotic resistance for carbenicillin and kanamycin |
| Figure S7-S10 | Pilot T7 RNAP differentiation experiments with varying inducer concentrations |
| Figure S11 | Assessing the evolutionary stability of burdensome T7 RNAP driven expression from a high copy ColE1-AmpR plasmid |
| Figure S12 | 1x and 2x naive evolution experiment fluorescence and OD data by plate generation |
| Figure S13 | 1x differentiation evolution fluorescence and OD data by plate generation |
| Figure S14 | 2x differentiation evolution experiment fluorescence and OD data by plate generation |
| Figure S15 | Assessment of leaky expression in the absence of T7 RNAP |
| Figure S16 | Fluorescence-based assay of dnaseI activity in cell lysate |
| Table S1 | Colony counts from dnaseI expression plasmid transformations |
| Figure S17 | Full circuit diagrams for 1x and 2x naive and differentiation circuits as depicted in 3.3 and 3.5 |
| Figure S18-S27 | Stochastic modeling of strategies incorporating plasmid mutation or plasmid loss/antibiotic degradation |
| Table S2 | Summary of strains used in this study |
| Table S3 | Naive 1X strain with IPTG inducible T7 RNAP |
| Table S4 | Naive 2X strain with IPTG inducible T7 RNAP |
| Table S5 | 1x differentiation strain with 2 copies salicylate inducible Bxb1-LAA |
| Table S6 | 2x differentiation strain with split-π protein design with 2 copies salicylate inducible Bxb1-LAA |
| Table S7 | Summary of plasmids used in this study. |
| | Determining inducer concentrations for differentiation experiments |
| | Supplementary methods |
| | Full description of model implementation |

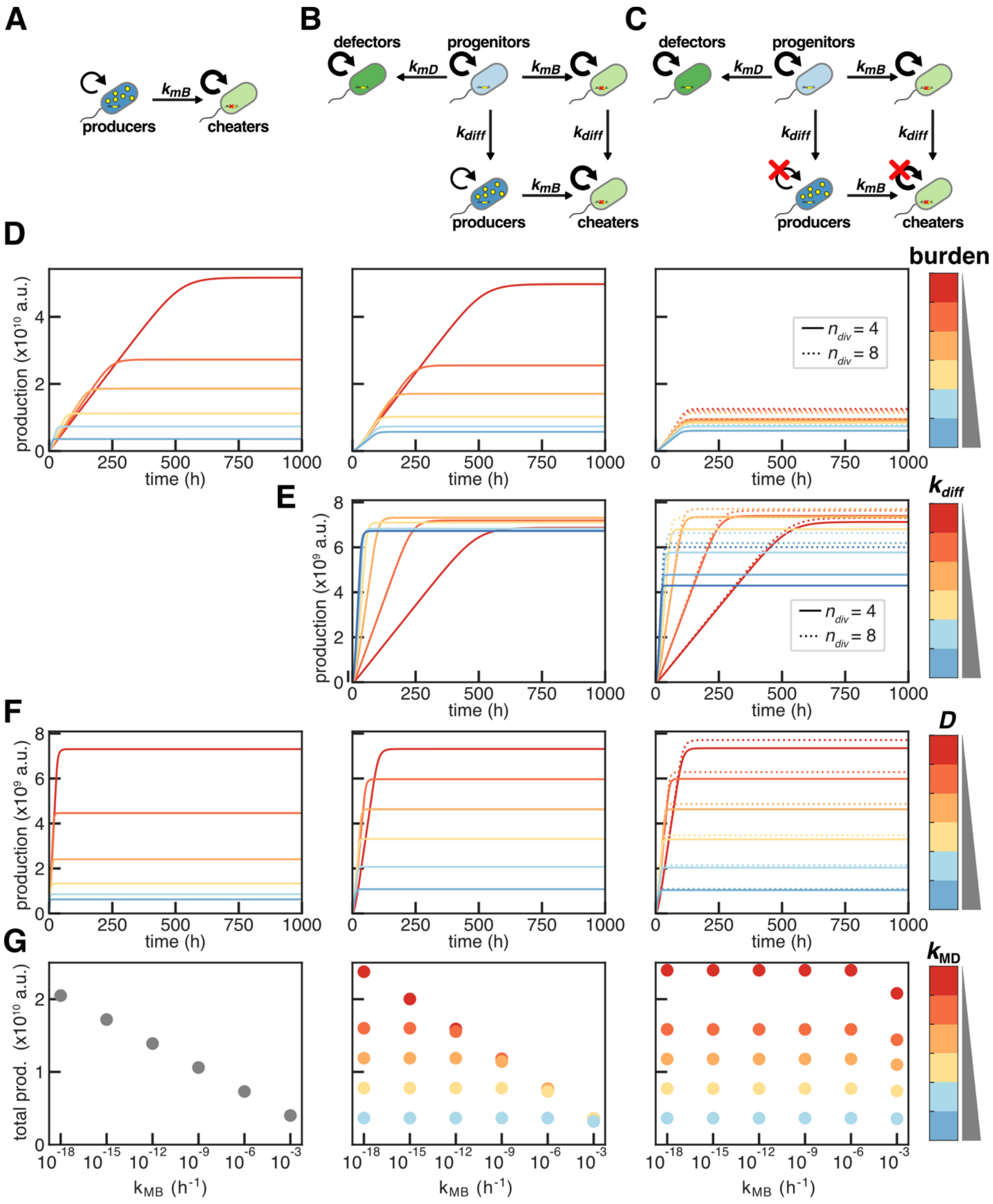


Figure S1: Deterministic modeling of strategies for expression of burdensome functions. **(A-C)** Schematics for a naive expression **(A)**, differentiation-activated expression (differentiation) **(B)**, and differentiation-activated expression in which the number of cell divisions following differentiation is limited (terminal differentiation) **(C)**. **(D-G)** Deterministic simulations circuits in chemostat with constant dilution and carrying capacity limited growth. $K = 10^9$ cells, initial population of $8\text{x}10^8$,$\mu_N = 2\ h^{-1}$, 1000 h duration. **(D)** Simulations with varying burden for naive (left), differentiation (center) and terminal differentiation (right). $k_{MB} = k_{MD} = 10^{-6}$ h$^{-1}$; $D = 0.2$ h$^{-1}$; $k_{diff} = 0.4$ h$^{-1}$; burden = {10, 20, 30, 50, 70, 90%}. **(E)** Simulations with varying differentiation rates ($k_{diff}$) for differentiation (center) and terminal differentiation (right). $k_{MB} = k_{MD} = 10^{-6}$ h$^{-1}$; $D = 0.2$ h$^{-1}$; 70% burden; $k_{diff}$ = {0.2, 0.4, 0.6, 0.8, 1.0, 1.2 h$^{-1}$}. **(F)** Simulations with varying dilution rates ($D$). $k_{MB} = k_{MD} = 10^{-6}$ h$^{-1}$; 70% burden; $k_{diff} = 0.4$ h$^{-1}$; $D$ = 0.2, 0.4, 0.6, 0.8, 1.0, 1.2 h$^{-1}$. **(G)** Endpoint total production of simulations with varying mutation rates. $k_{diff} = 0.4$ h$^{-1}$; $D = 0.2$ h$^{-1}$; 70% burden; $k_{MB}$ = {$10^{-18}$, $10^{-15}$, $10^{-12}$, $10^{-9}$, $10^{-6}$, $10^{-3}$ h$^{-1}$}; $k_{MD}$ = {$10^{-18}$, $10^{-15}$, $10^{-12}$, $10^{-9}$, $10^{-6}$, $10^{-3}$ h$^{-1}$}.

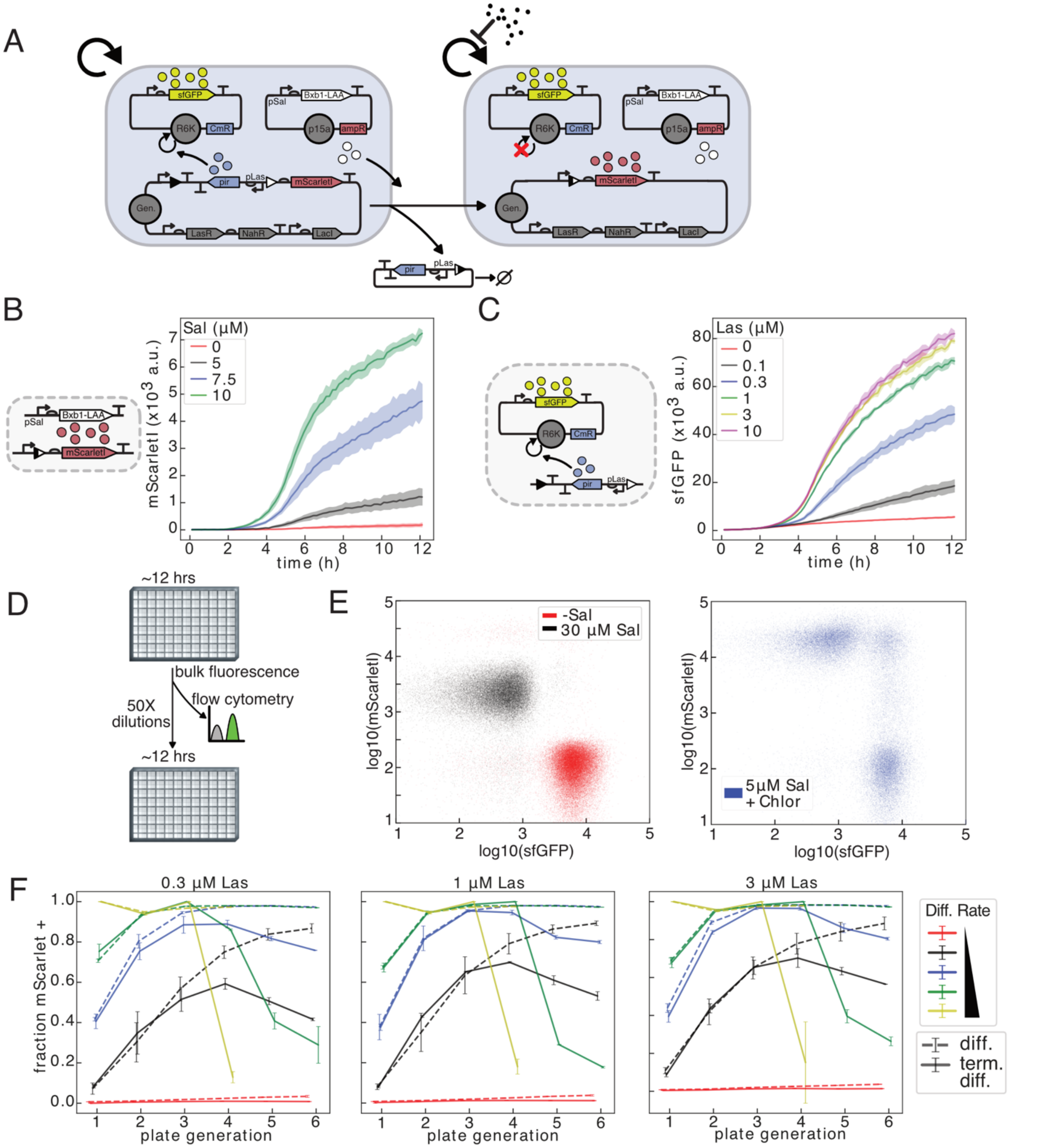


Figure S2: Implementation of a tunable integrase-mediated differentiation circuit. (A) Schematic of a tunable integrase differentiation circuit. Las-AHL and salicylate (sal) induce expression of *pir* encoded π protein and degradation-tagged Bxb1, respectively. (B-C) Batch culture experiments of JS006 with circuit depicted in (A) grown in M9CA media. (B) mScarletI fluorescence with varying induction levels of sal in M9CA + carb. (C) sfGFP fluorescence with varying induction levels of Las-AHL in M9CA + carb/chlor. (B-C) Mean +/- standard deviation of three replicates. (D-F) Cells are grown in 300 µL M9CA media with varying inducer concentrations, and diluted 50X every 12 hours. Samples are taken for flow cytometry after each growth. (E) Flow cytometry results after the third 12-hour growth for cells grown in 0.3 µM Las-AHL with carb +/- 30 µM sal(left), and cells grown media with carb/chlor, 0.3 µM Las-AHL, 5 µM sal(right). (F) Results of flow cytometry analysis of cells grown for six consecutive 12-hour growths in varying inducer concentrations. Average fraction mScarletI positive for two replicate wells +/- standard deviation.

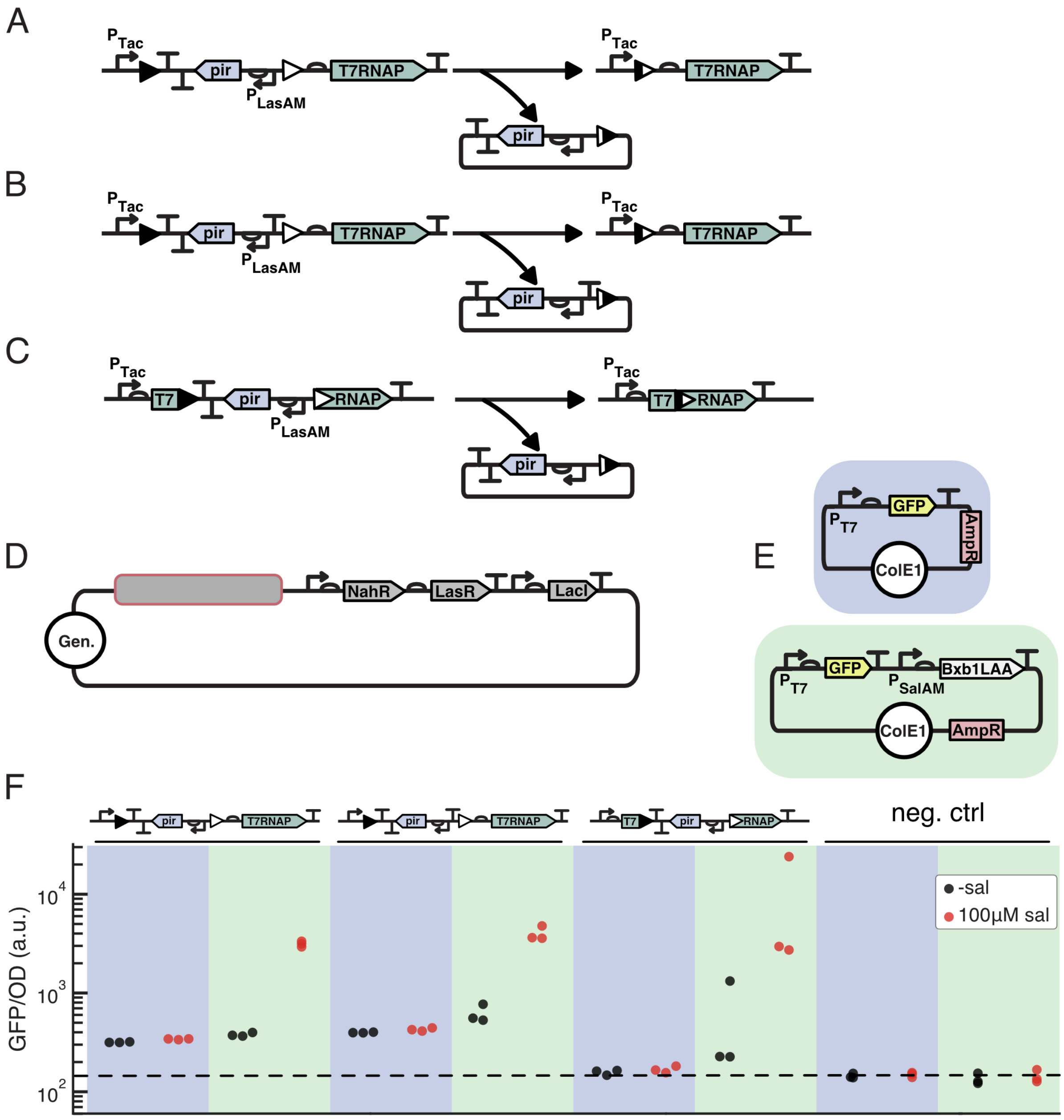


Figure S3: Integrase mediated activation allows leak-free expression of T7 RNAP. (A-D) Circuit designs for Bxb1 integrase activated expression of genomically integrated T7 RNAP. (A-B) Integrase recombination joins pTac promoter with RBS and full length CDS with (B) or without (A) a terminator in front of the RBS. (C) Integrase recombination joins promoter, RBS, and left half of CDS with right half of CDS. (F) JS006 negative control and genomically integrated strains were transformed with a ColE1 plasmid encoding T7 RNAP-driven GFP alone (blue shaded, F: top) or with salicylate inducible Bxb1-LAA (green shaded, F: bottom). OD normalized fluorescence recorded after 12 h of growth in LB + carb with (red) or without (black) 100 µM salicylate.

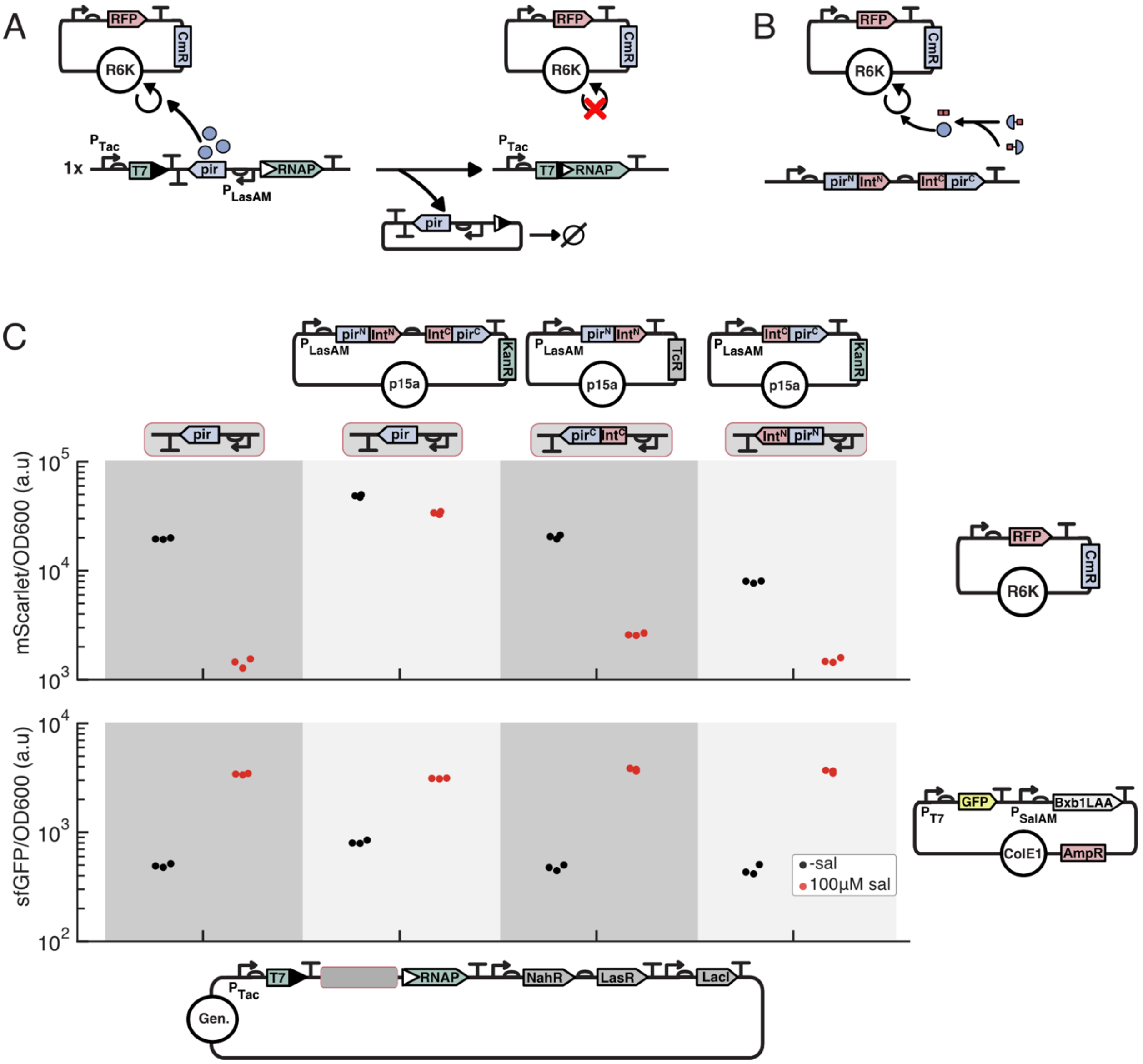


Figure S4: Integrase inactivation of π protein expression ablates R6K plasmid replication. (A) Schematic of integrase excision of *pir* resulting in loss of π protein and cessation of R6K plasmid replication. (B) Design of a split-π protein system in which each half is tagged with an intein fragment, and generate a full-length protein upon protein trans-splicing. (C) Strains harboring genomic integrations of circuits in which Bxb1 recombination simultaneously activates the expression of T7 RNAP and inactivates full-length or split-π protein, and containing an R6K plasmid with chlorR and constitutive mScarletI w/ or w/o a p15a plasmid encoding one or both halves of the split-π protein, were transformed with a ColE1-AmpR plasmid with $P_{T7}$ GFP and $P_{SalTTC}$ Bxb1-LAA. OD normalized RFP (top) and RFP (bottom) were recorded after 18 h of growth in LB w/ 100 µg/mL carb, 34 µg/mL chlor, 30 nM Las-AHL, with (red) or without (black) 100 µM salicylate.

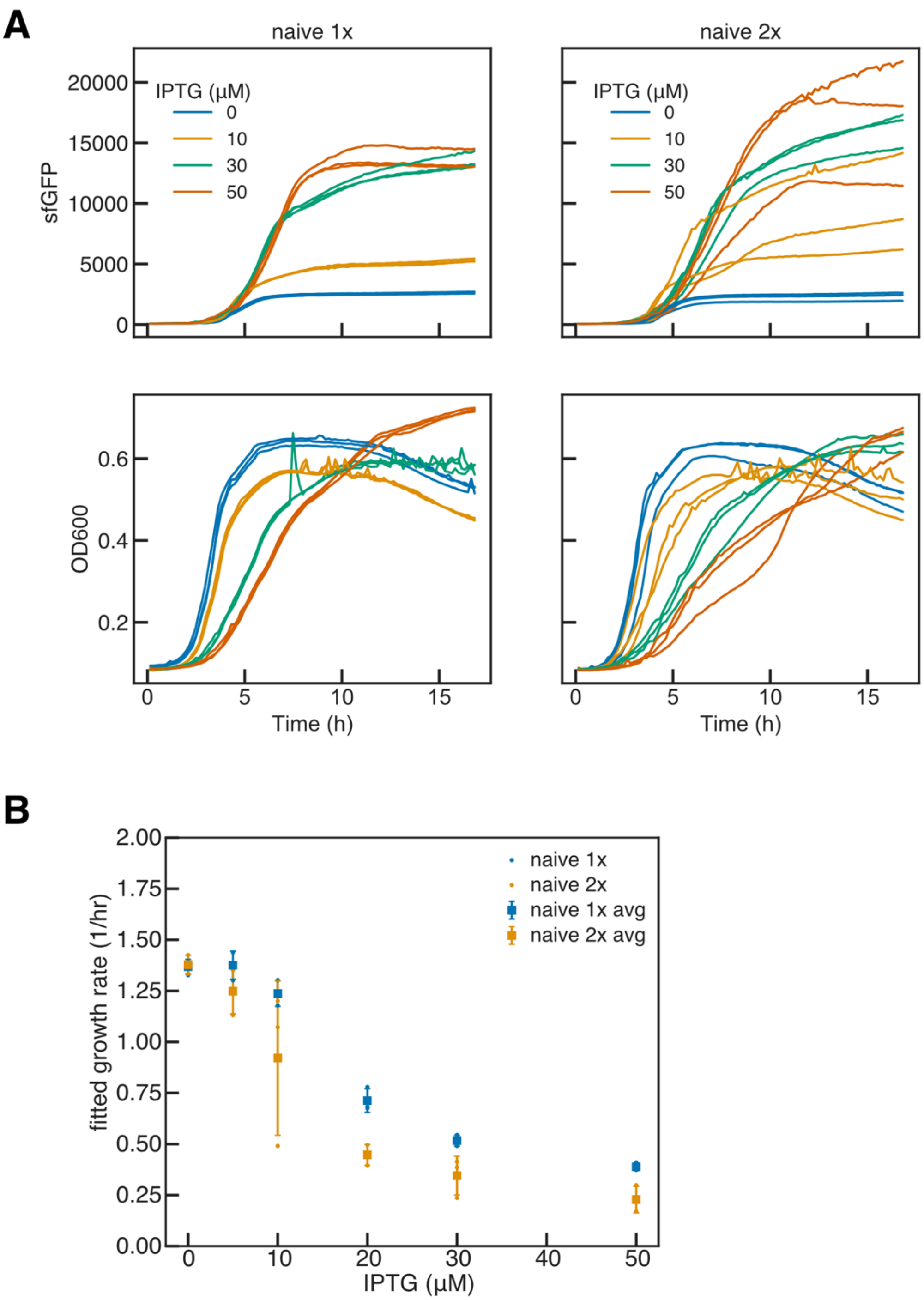


Figure S5: Characterization of burden level of naive T7 RNAP-driven GFP. JS006 cells with 1 and 2 copies of naive IPTG inducible T7 RNAP were transformed with a high copy ColE1 plasmid with kanamycin resistance. Cells were outgrown in LB with kanamycin (50 µg/mL) for ~6 hours then diluted 1:50 into LB kan with varying concentrations of IPTG. (A) GFP production (top) and OD600 (bottom) were measured every 10 minutes for 1x naive (left) and 2x naive (right) grown in triplicate at 37C in 0.3 mL. (B) OD600 curves were trimmed to 60 percent of maximum OD600 achieved and used to fit an exponential growth model with noise floor, initial population, and growth rate parameters. Mean growth rate +/- SD of 3 replicates fitted separately plotted for naive 1x and 2x.

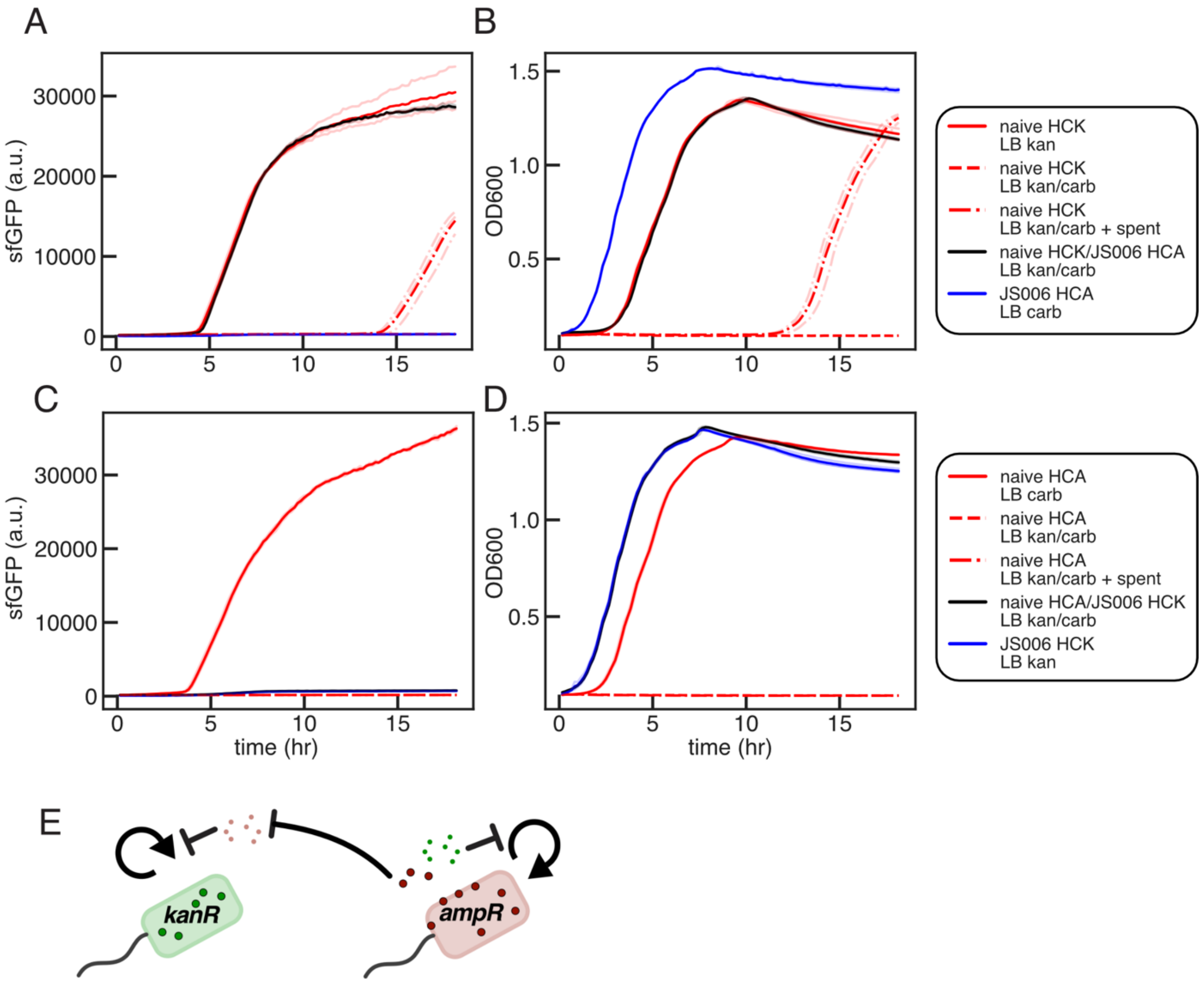


Figure S6: Assessing communal antibiotic resistance for carbenicillin and kanamycin. JS006 and naive 1x cells were transformed with ColE1-AmpR-$P_{T7}$-GFP or ColE1-KanR-$P_{T7}$-GFP, and colonies outgrown for 8 h in LB with kan (50 µg/mL) or carb (100 µg/mL). Spent media was harvested by pelleting JS006/ ColE1-AmpR-$P_{T7}$-GFP or JS006/ ColE1-KanR-$P_{T7}$-GFP and filtering supernatant (0.2 µM). Cultures were diluted 1:50 into 0.2 mL experimental monocultures or co-cultures in triplicate in 96 well plates, and OD600 and sfGFP (485/515nM) monitored over 18 h growth. (A-B) naive 1x + HCK pT7 GFP and JS006 + ColE1-AmpR-$P_{T7}$-GFP grown in mono- and co-culture. JS006 alone (blue) grows (B) but does not produce GFP (A). naive + ColE1-KanR-$P_{T7}$-GFP grows and produces GFP in LB Kan, in co-culture with JS006 HCA in LB carb/kan, and with delay in LB carb/kan/10 percent JS006 HCA spent media, but not in LB kan/carb. (C-D) naive 1x + ColE1-AmpR-$P_{T7}$-GFP and JS006 + ColE1-KanR-$P_{T7}$-GFP grown in mono- and co-culture. (E) Schematic of shared antibiotic resistance. AmpR cells degrade carb inside the cell, in the periplasm, as well as in the medium, allowing amp sensitive cells to grow. Antibiotic resistance from KanR cells however is not shared significantly enough to allow growth of sensitive cells.

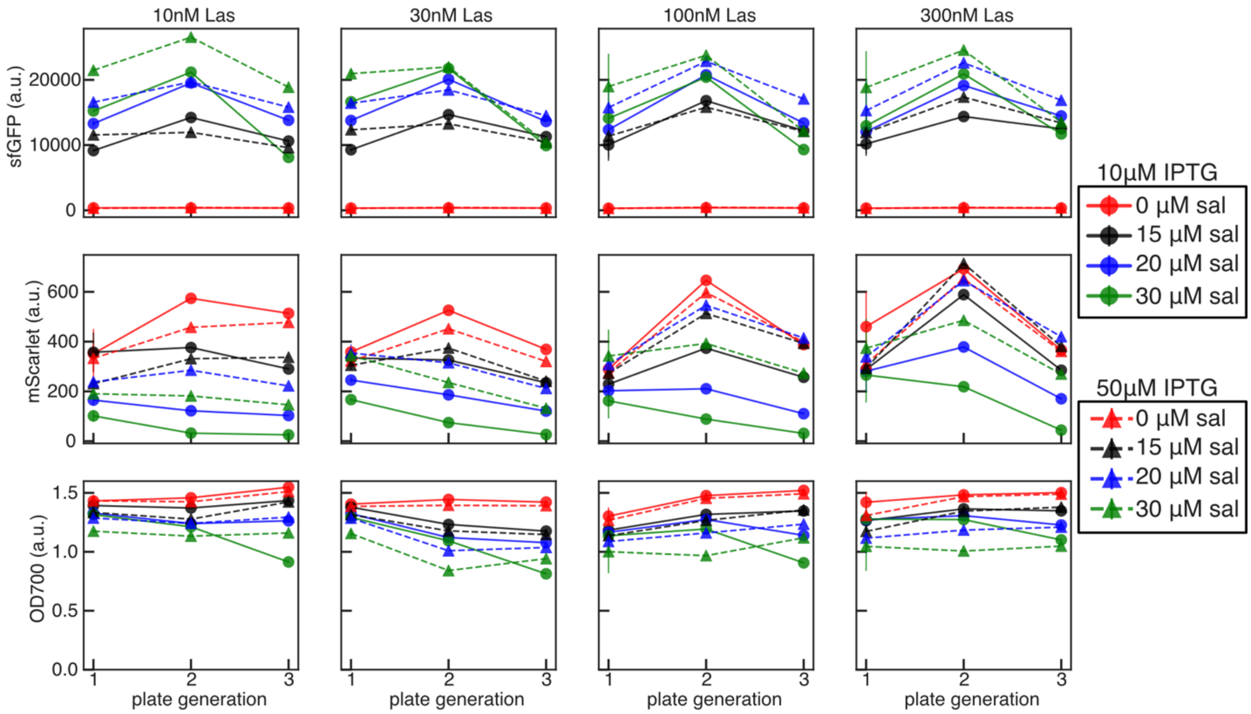


Figure S7: 1x differentiation with ColE1-KanR-$P_{T7}$-GFP and R6K-CmR-mScarlet; Las-AHL, salicylate, and IPTG gradient fluorescence and OD data by plate generation. Three replicate colonies outgrown for 8 hours in LB kan/chlor with varying concentrations of Las-AHL, then diluted 1:50 every 8 hours into 0.3 mL LB kan/chlor/Las-AHL with varying concentrations of sal and IPTG. Mean +/- SD of GFP (top), mScarlet (middle), and OD700 (bottom) plotted for three total growths. Color indicates sal concentration; 10 µM IPTG (circles, solid lines), 50µM IPTG (triangles, dashed lines).

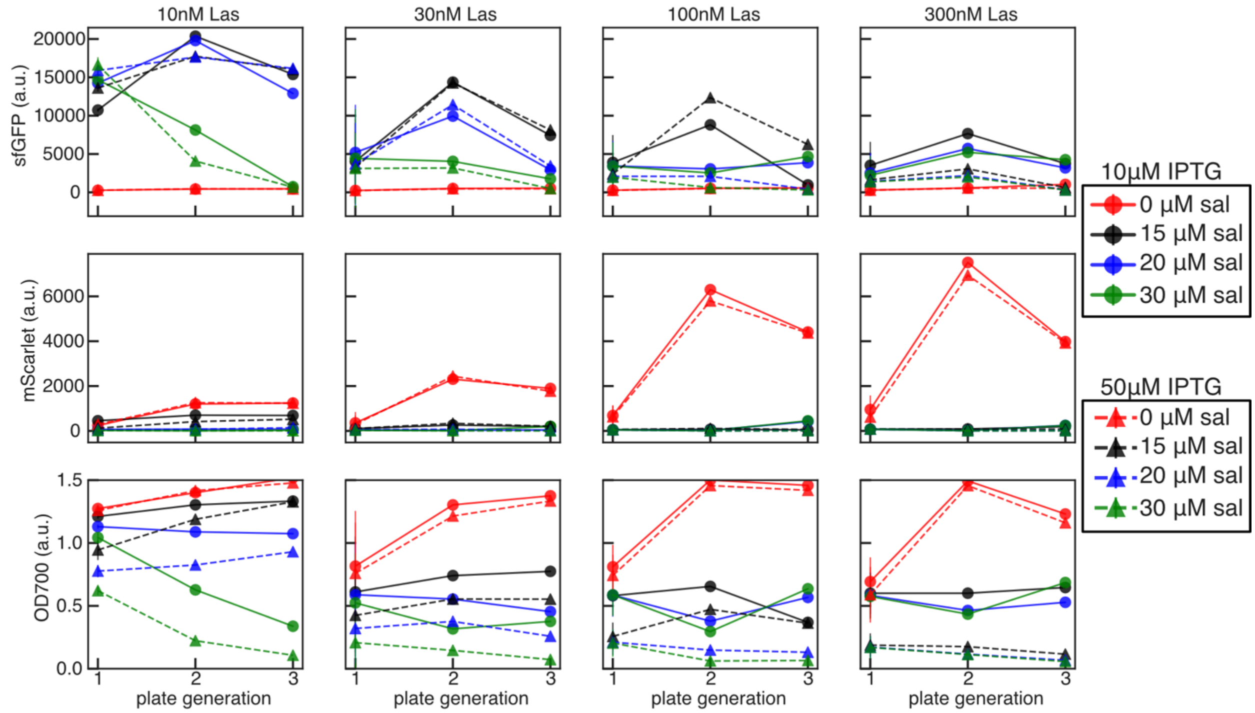


Figure S8: 2x differentiation with ColE1-KanR-$P_{T7}$-GFP and R6K-CmR-mScarlet; Las-AHL, salicylate, and IPTG gradient fluorescence and OD data by plate generation. Three replicate colonies outgrown for 8 hours in LB kan/chlor with varying concentrations of Las-AHL, then diluted 1:50 every 8 hours into 0.3 mL LB kan/chlor/Las with varying concentrations of sal and IPTG. Mean +/- SD of GFP (top), mScarlet (middle), and OD700 (bottom) plotted for three total growths. Color indicates sal concentration; 10 µM IPTG (circles, solid line), 50 µM IPTG (triangles, dashed line).

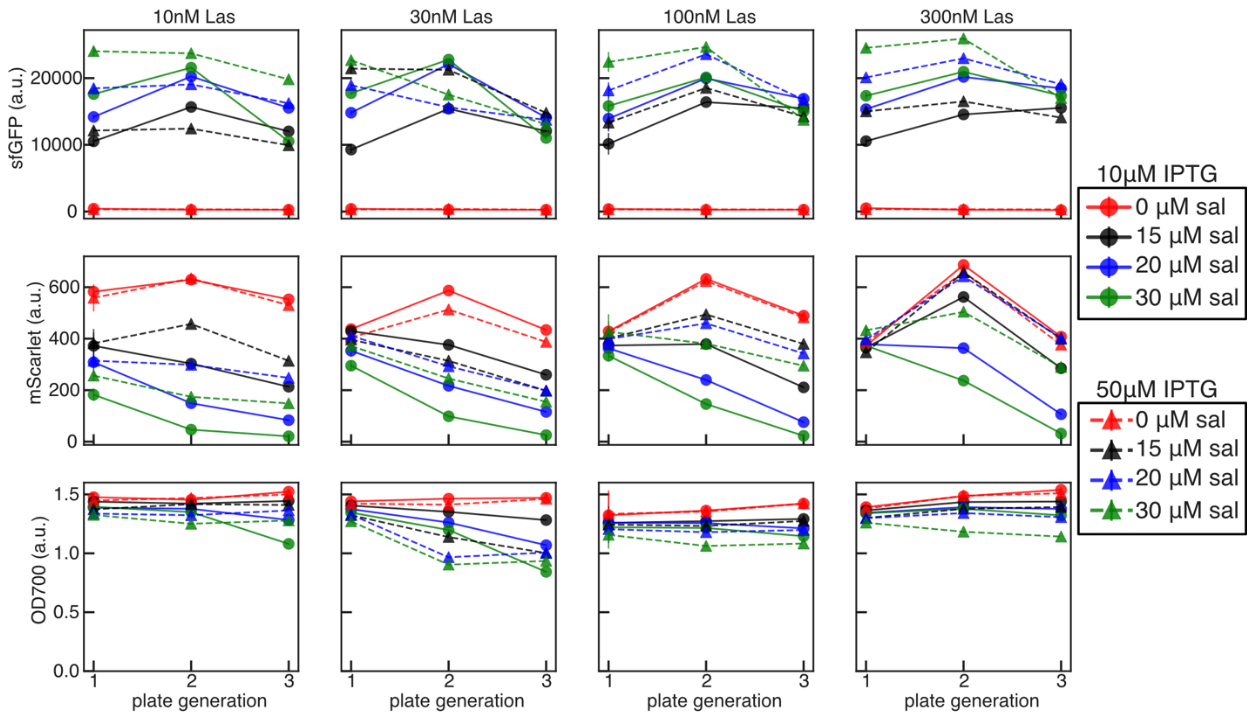


Figure S9: 1x differentiation with ColE1-AmpR-$P_{T7}$-GFP and R6K-CmR-mScarlet; Las-AHL, salicylate, and IPTG gradient fluorescence and OD data by plate generation. Three replicate colonies outgrown for 8 hours in LB carb/chlor with varying concentrations of Las-AHL, then diluted 1:50 every 8 hours into 0.3 mL LB carb/chlor/Las with varying concentrations of sal and IPTG. Mean +/- SD of GFP (top), mScarlet (middle), and OD700 (bottom) plotted for three total growths. Color indicates sal concentration; 10 µM IPTG (circles, solid lines), 50 µM IPTG (triangles, dashed lines).

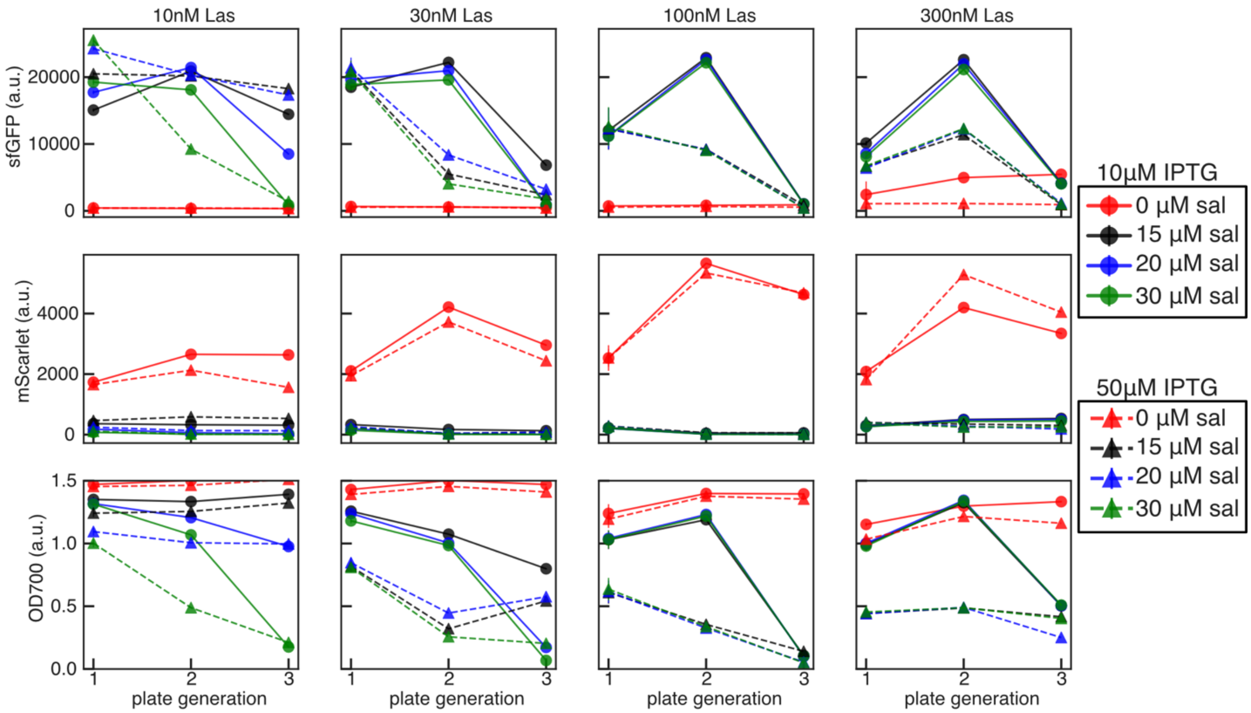


Figure S10: 2x differentiation with ColE1-AmpR-$P_{T7}$-GFP and R6K-CmR-mScarlet; Las-AHL, salicylate, and IPTG gradient fluorescence and OD data by plate generation. Three replicate colonies outgrown for 8 hours in LB carb/chlor with varying concentrations of Las-AHL, then diluted 1:50 every 8 hours into 0.3 mL LB carb/chlor/Las-AHL with varying concentrations of sal and IPTG. Mean +/- SD of GFP (top), mScarlet (middle), and OD700 (bottom) plotted for three total growths. Color indicates sal concentration; 10 µM IPTG (circles, solid lines), 50 µM IPTG (triangles, dashed lines).

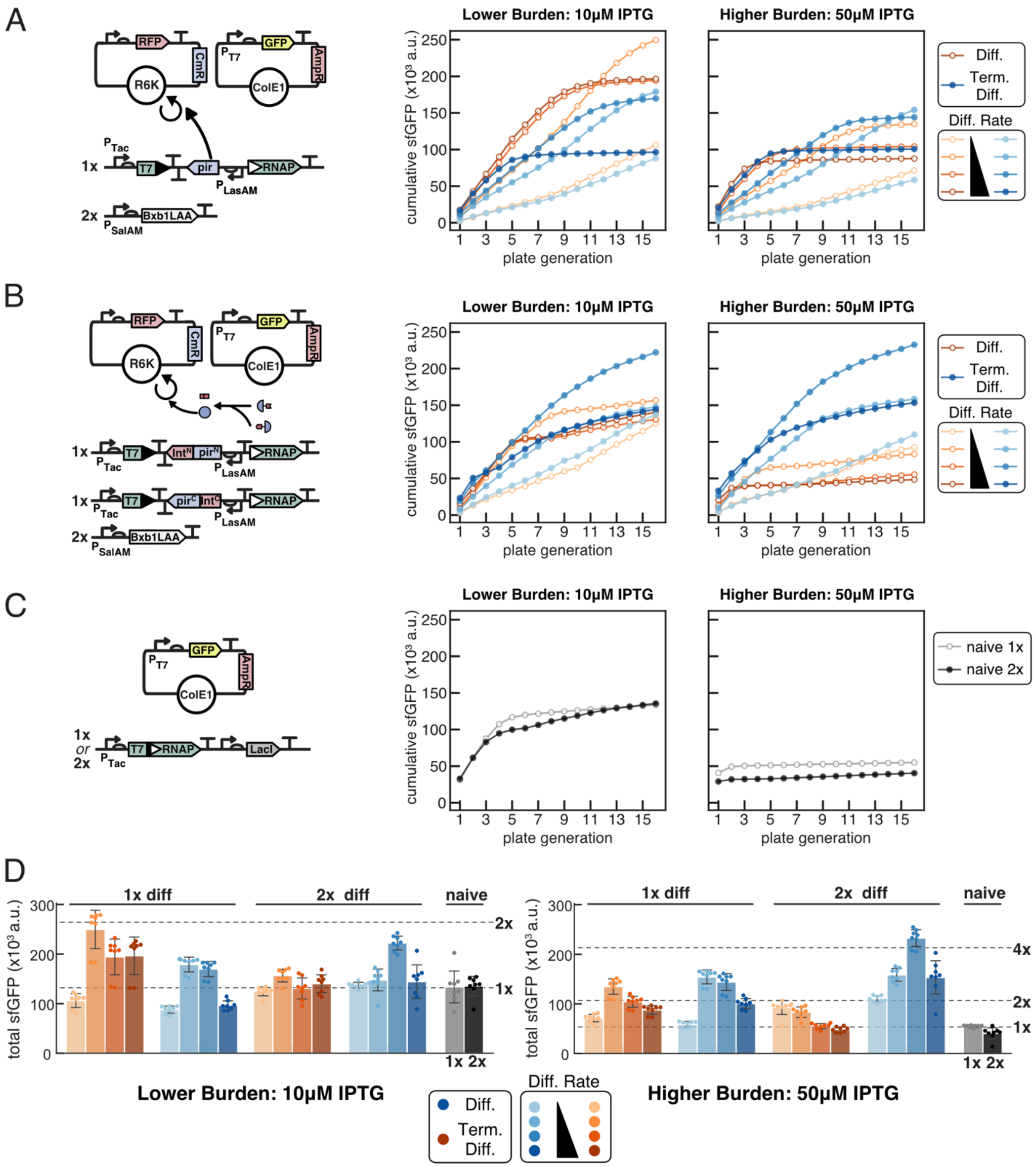


Figure S11: Assessing the evolutionary stability of burdensome T7 RNAP driven expression from a high copy ColE1-AmpR plasmid. (A-C) 8 independent transformants were outgrown for 8 hours in LB media with appropriate antibiotics and inducers before 50x dilution into experimental conditions. Cells were grown in 96 well plates in 0.3 mL, diluted 50x every 8 h for 16 total growths, and 50 μL endpoint samples taken to measure OD700, sfGFP (485/515 nm), and mScarlet (565/595 nM) fluorescence in 384 well matriplates. Average cumulative sfGFP production plotted for each condition. (A-B) 1x differentiation and

2x differentiation. Each differentiation cassette additionally encodes NahR$_{AM}$, LasR$_{AM}$ and LacI$_{AM}$ (Figure 3.27 for full circuit diagram). Cells were co-transformed with ColE1-AmpR-P$_{T7}$-GFP and R6K-CmR-mScarlet, and plated on LB +carb/chlor/30 nM Las-AHL. Colonies were outgrown in LB +carb/chlor/10 nM Las-AHL before 50x dilution into experimental conditions in LB carb with chlor (blue, filled circles or without chlor (orange, open circles) with varying concentrations of salicylate (10, 15, 20, 30 µM) and IPTG (10, 50 µM). (C) Cells with one (naive 1x) or two (naive 2x) copies of genomically integrated T7 RNAP were transformed with ColE1-AmpR-P$_{T7}$-GFP, plated on LB + carb, and outgrown in LB + carb before dilution into experimental conditions. (D) Total cumulative production +/- SD after 16 growths for all strains in all conditions.

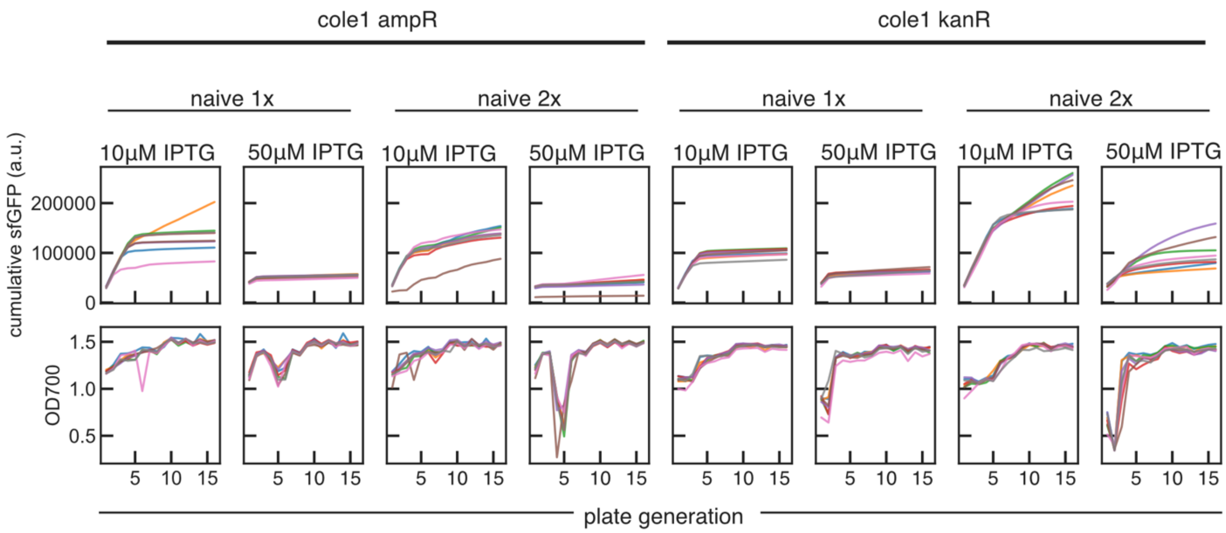


Figure S12: 1x and 2x naive evolution experiment fluorescence and OD data by plate generation. (A-B) Cumulative endpoint sfGFP fluorescence (top row) and OD700 (bottom row) for individual experimental replicates measured in 50 µL in 384 well matriplates. (Left) Data from experiment with ColE1-AmpR-P$_{T7}$-GFP of cells grown in LB carb with varying concentrations of IPTG. (B) (Right) Data from experiment with ColE1-KanR-P$_{T7}$-GFP of cells grown in LB kan with varying concentrations of IPTG.

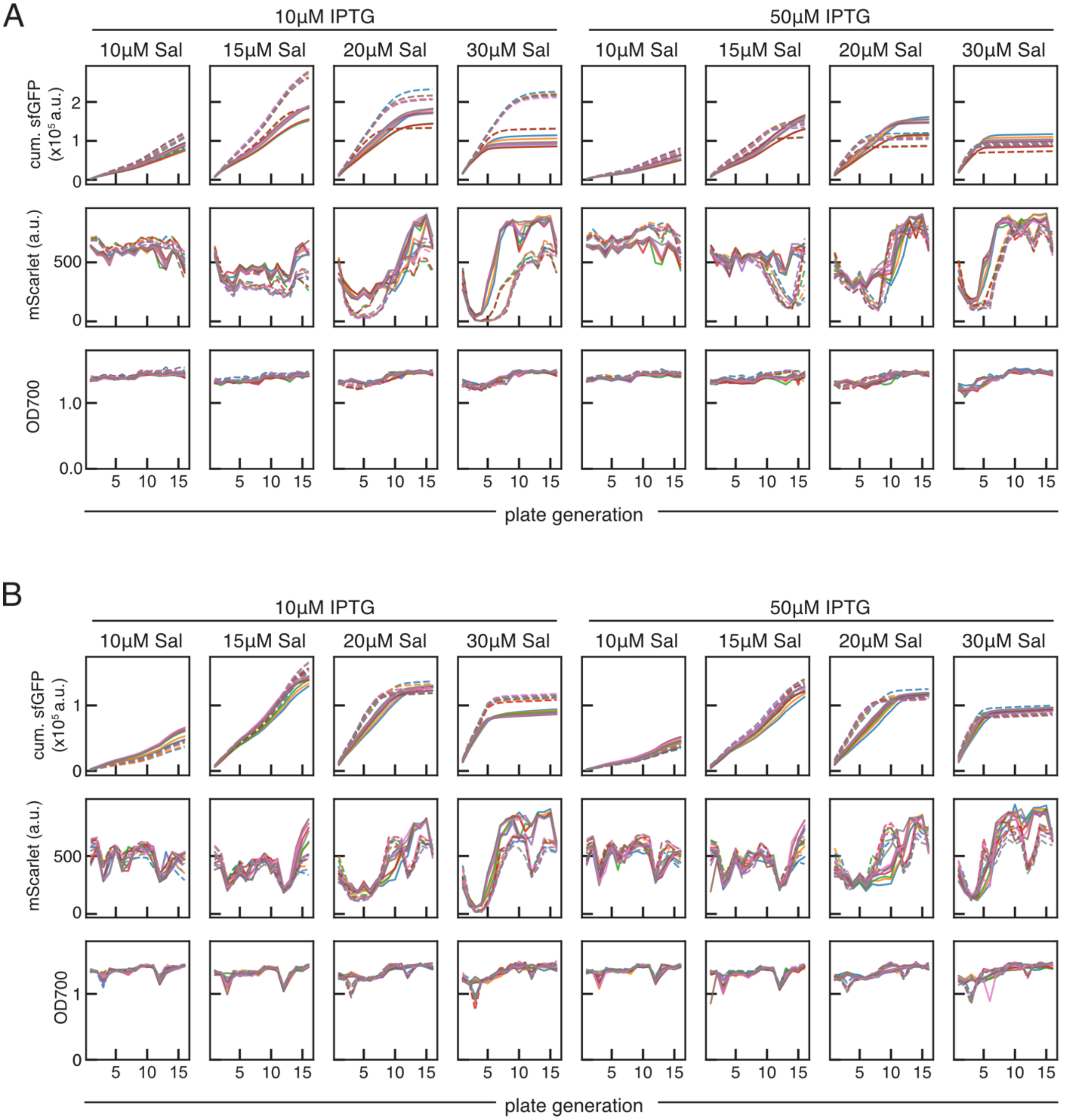


Figure S13: 1x differentiation evolution fluorescence and OD data by plate generation. (A-B) Cumulative endpoint sfGFP fluorescence (top row), endpoint mScarletI fluorescence (middle row) and OD700 (bottom row) for individual experimental replicates measured in 50 µL in 384 well matriplates. (A) Data from experiment with ColE1-AmpR-$P_{T7}$-GFP of cells grown in LB carb/chlor/10 nM Las (solid lines) and carb/10 nM Las (dashed lines) with varying concentrations of IPTG. (B) Data from experiment with ColE1-KanR-$P_{T7}$-GFP of cells grown in LB kan/chlor/10 nM Las (solid lines) and kan/10 nM Las (dashed lines) with varying concentrations of IPTG.

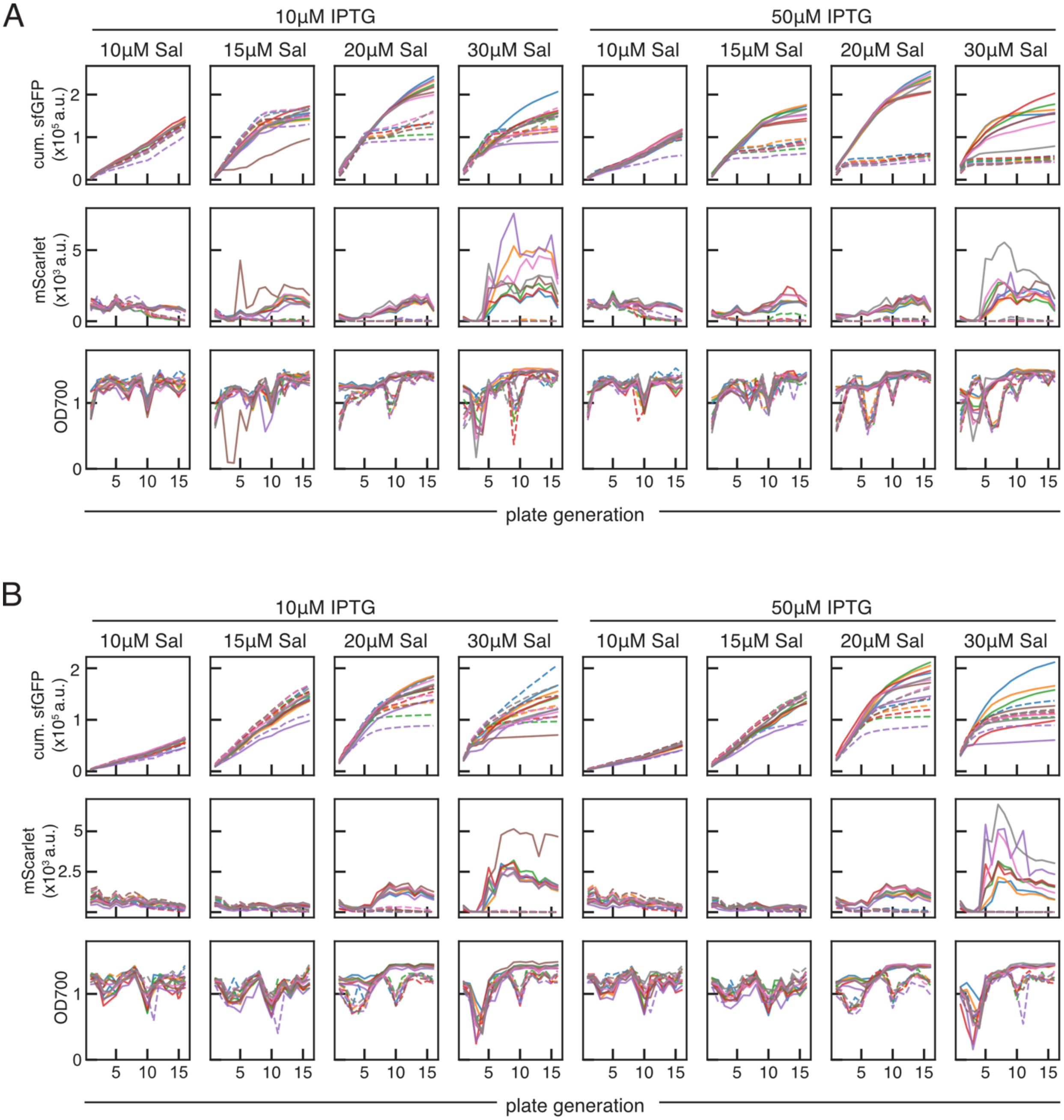


Figure S14: 2x differentiation evolution experiment fluorescence and OD data by plate generation. (A-B) Cumulative endpoint sfGFP fluorescence (top row), endpoint mScarletI fluorescence (middle row) and OD700 (bottom row) for individual experimental replicates measured in 50 µL in 384 well matriplates. (A) Data from experiment with ColE1-AmpR-$P_{T7}$-GFP of cells grown in LB carb/chlor/10 nM Las (solid lines) and carb/10 nM Las (dashed lines) with varying concentrations of IPTG. (B) Data from experiment with ColE1-KanR-$P_{T7}$-GFP of cells grown in LB kan/chlor/10 nM Las (solid lines) and kan/10 nM Las (dashed lines) with varying concentrations of IPTG.

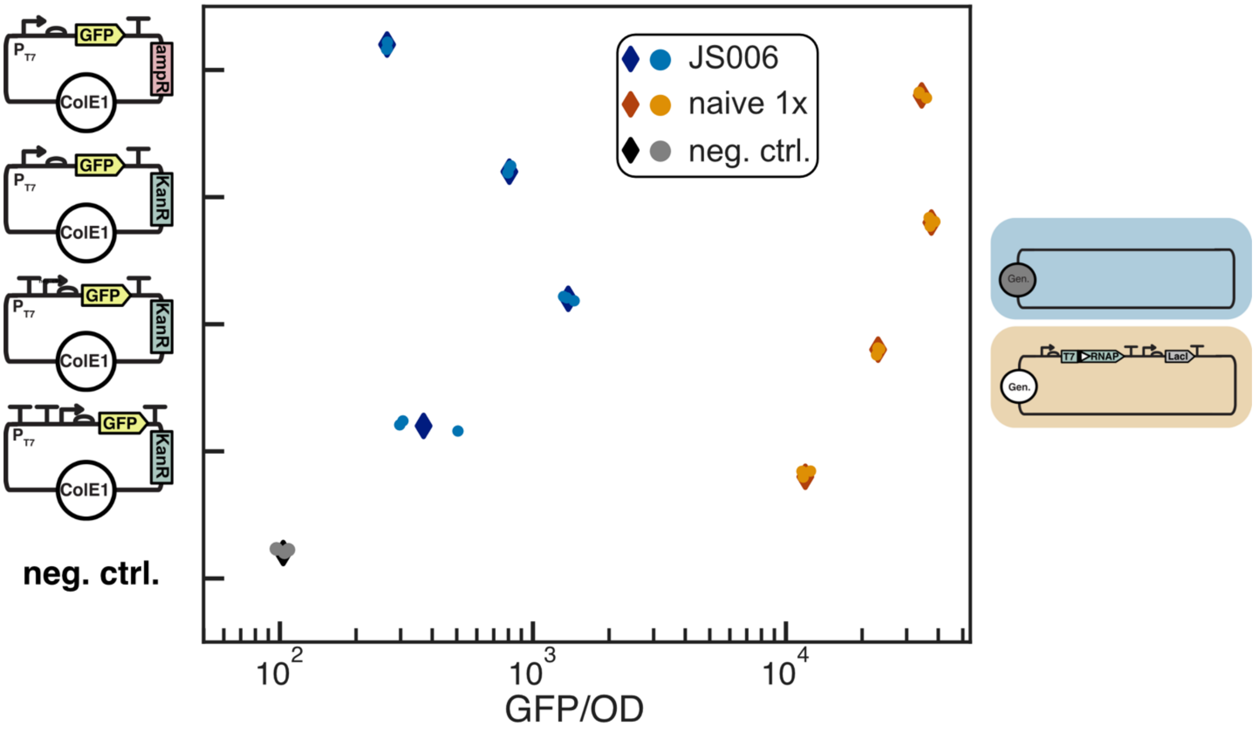


Figure S15: Assessment of leaky expression in the absence of T7 RNAP. JS006 and naive 1x cells were transformed with plasmids containing $P_{T7}$-B0034-sfGFP-T7T, with 0, 1, or 2 insulating terminators. OD600 normalized sfGFP fluorescence reported, mean (diamonds) +/- SD of three replicates (circles) after 12 h growth compared to negative control JS006 lacking any GFP expression plasmid.

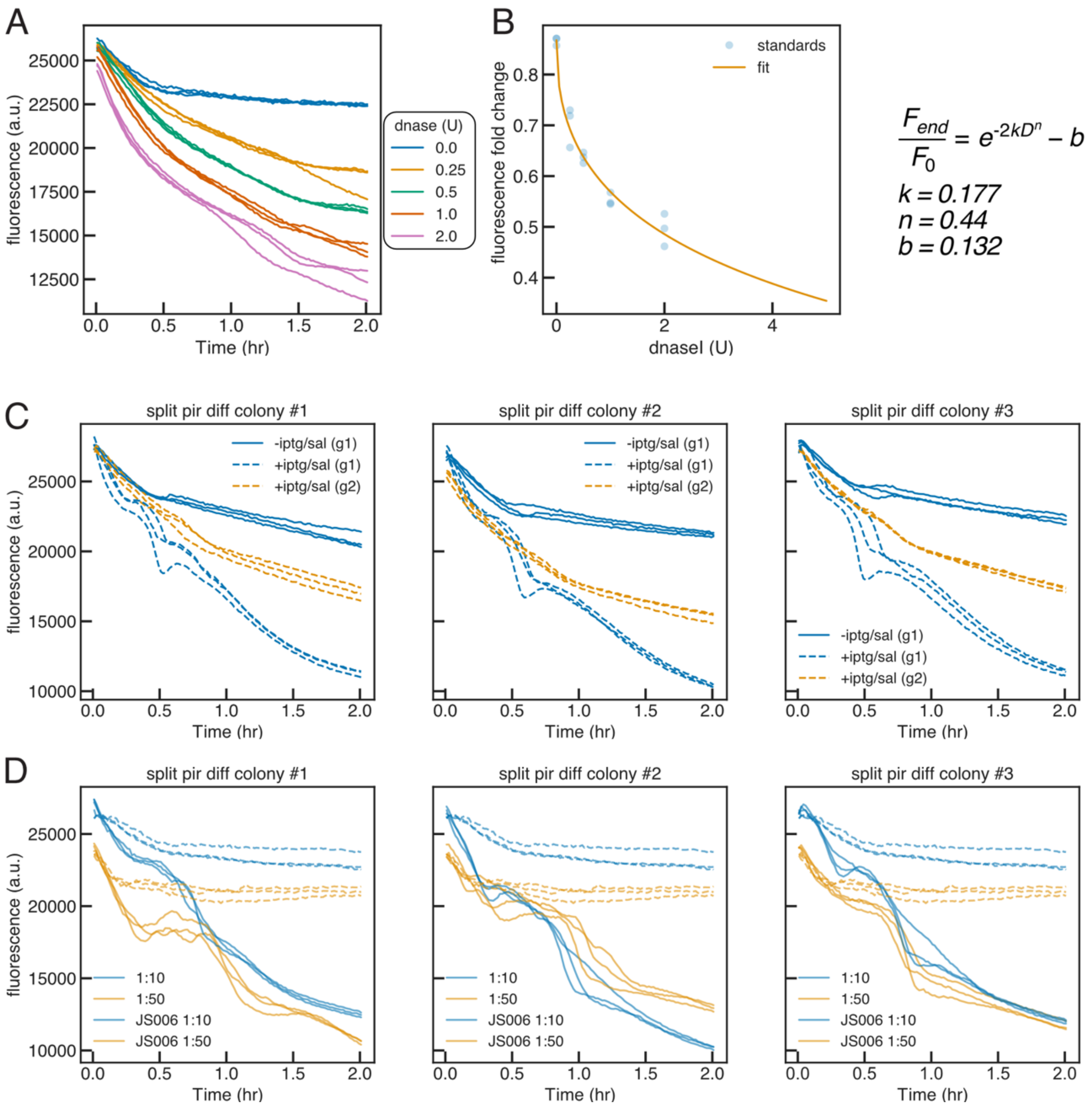


Figure S16: Fluorescence-based assay of dnaseI activity in cell lysate. (A) JS006 lysate diluted 1:10 in 0.85 percent NaCl, with 0, 0.25, 0.5, 1, or 2 U of dnaseI per 10 µL volume diluted sample. 10 µL sample added to 190 µL dnaseI assay buffer (100 mM sodium acetate/5 mM magnesium sulfate, 5 µg calf thymus DNA,1:1000 SYBR Safe). Fluorescence (487/528 nm) time-course of samples in triplicate measured every minute for 2 hours. (B) Fluorescence fold-change (endpoint/initial) used to a fit a model, where *b* describes background loss of fluorescence through photo-bleaching or other non-dnaseI related means of loss of fluorescence, *k* is the first-order rate constant describing the degradation of DNA by dnaseI, *D* is the concentration of dnaseI (U/rxn), *n* is a phenomenological constant which captures the non-linear relationship between dnaseI concentration and observed loss of fluorescence, and 2 is the time in hours for which the assay was ran. (C-D) Time-course traces of dnaseI assay of lysate from three independent experiments. (C) Assay performed on 1:10 dilutions of lysate from uninduced first growth (blue solid), and induced first (dashed blue) and second (dashed orange) growths. (D) Re-assayed lysate samples of induced first growth (solid) diluted 1:10 (blue) and 1:50 (orange), and JS006 lysate (dashed) diluted 1:10 (blue) and 1:50 (orange).

| Strain | GFP plasmid | dnaseI plasmid |
|---|---|---|
| **1x naive** | >$10^4$ cfu | 1 cfu |
| **2x naive** | >$10^4$ cfu | 0 cfu |
| **1x diff** | >$6.6x10^3$ cfu | 590 cfu |
| **2x diff** | >$10^4$ cfu | 1090 cfu |

Table S1: Colony counts from dnaseI expression plasmid transformations. 50 µL chemically competent cells were transformed with 10 ng of ColE1-AmpR-$P_{T7}$-sfGFP or 10 ng ColE1 AmpR T13m-T12m- $P_{T7}$-B0032-dnsaeI-T7T, and 5 µL or 50 µL plated on LB carb. >$10^4$ indicates more than 1000 colonies from plating 5 µL of transformation.

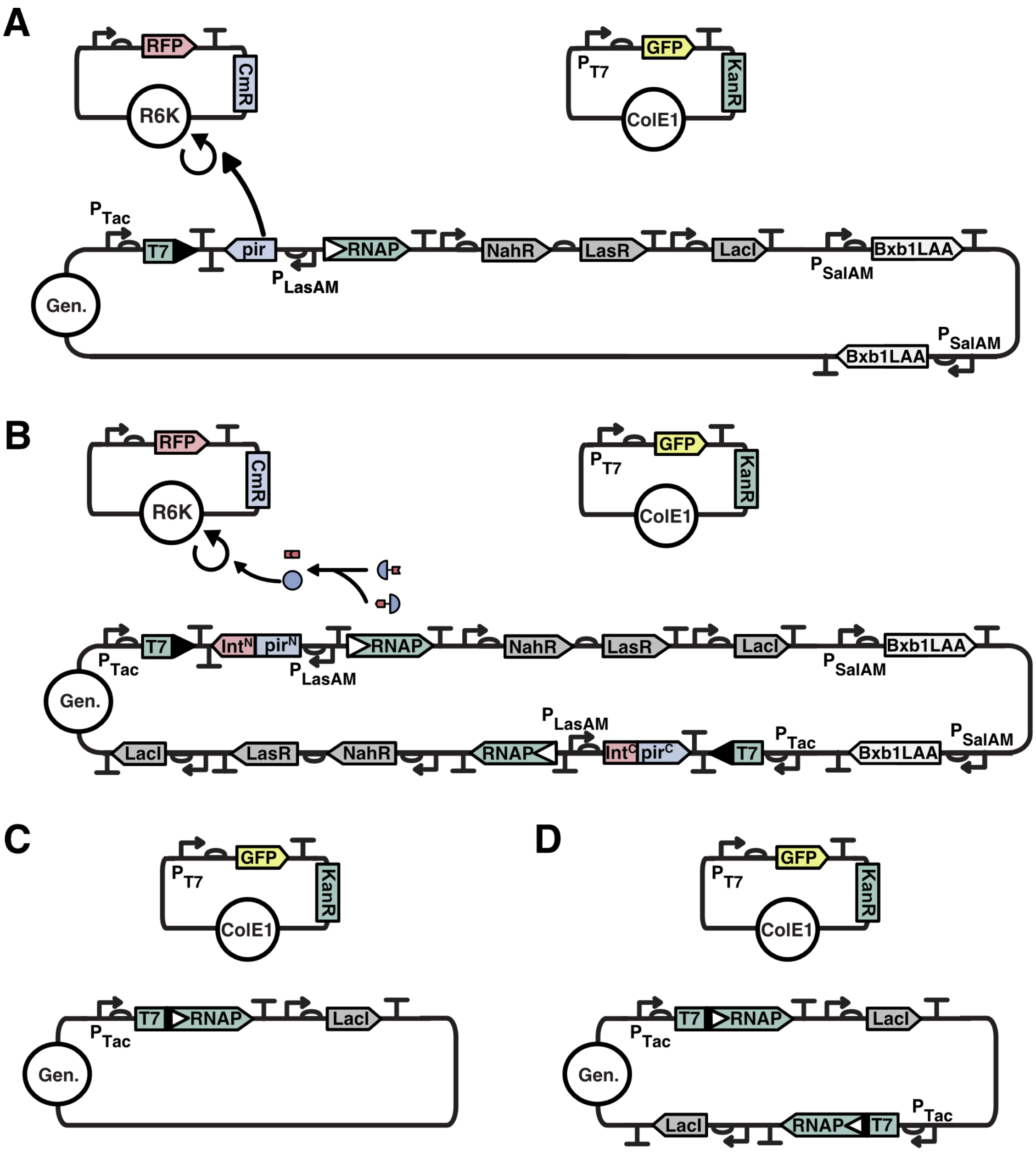


Figure S17: Full circuit diagrams for 1x and 2x naive and differentiation circuits as depicted in 3.3 and 3.5. (A) 1x differentiation circuit diagram. Differentiation cassette encoding excisable *pir*, recombination activated T7 RNAP, and NahR$_{AM}$, LasR$_{AM}$, and LacI$_{AM}$, is integrated with clonetegration[38] at the P21 (T) landing site. 2 copies of pSal$_{TTC}$-B0034-Bxb1LAA-T2m were integrated at the primary and secondary 186 (O) landing sites. (B) 2x differentiation circuit diagram. As in (A), with the the T site integration encoding the N-terminal portion of the split-$\pi$ protein, and the second differentiation cassette integrated at the HK022 (H) site encoding the C-terminal portion of the split-$\pi$ protein. (C) 1x naive expression cassette encoding inducible T7 RNAP and LacI$_{AM}$ integrated at the T site. (D) As in (C) with second identical integration at the H site.

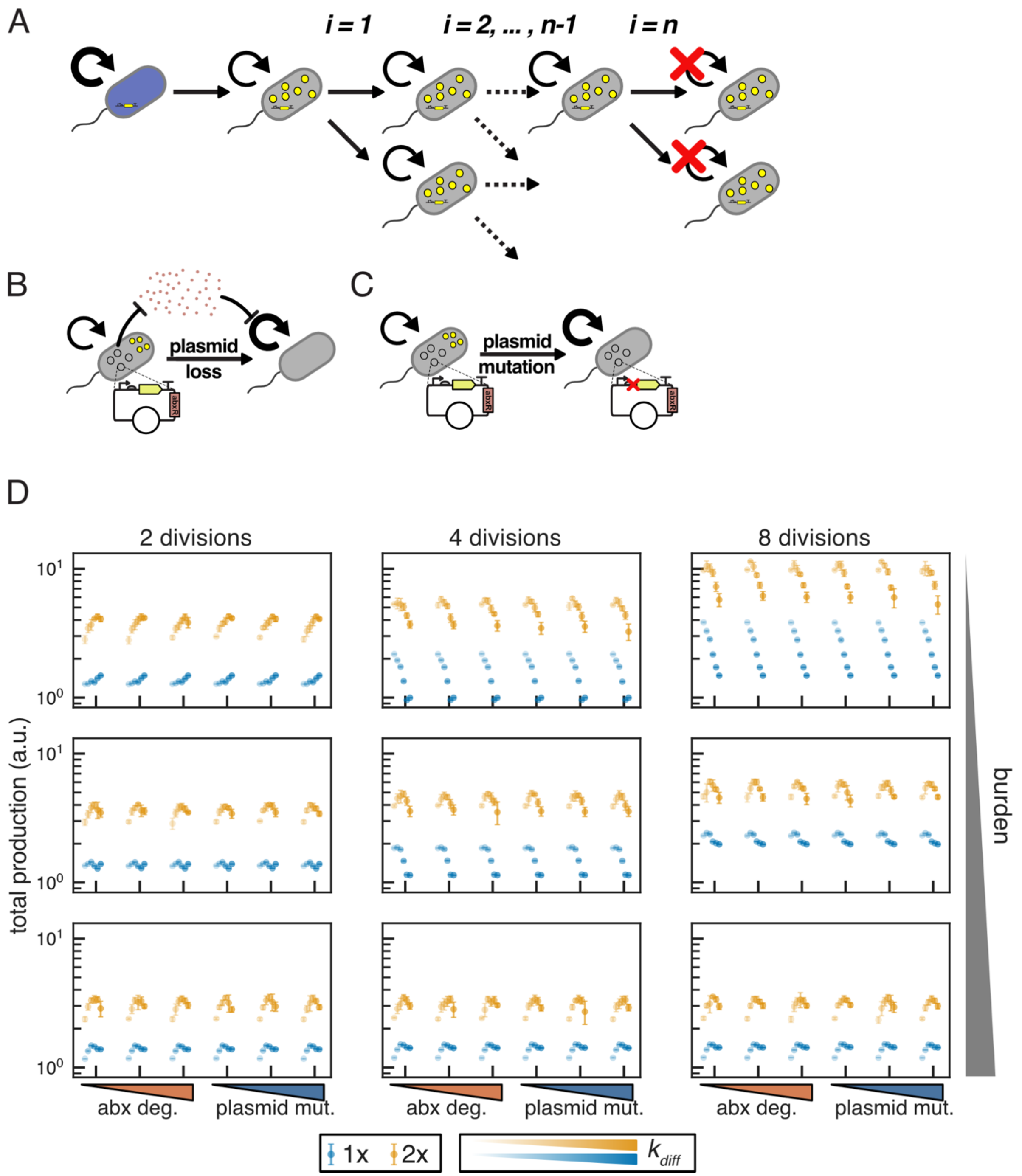


Figure S18: Effect of post-differentiation number of divisions on terminal differentiation architectures. (A) Following differentiation, cells grow and divide exponentially for *n* divisions. (B) Schematic describing stochastic loss of a burdensome plasmid which encodes an antibiotic resistance gene that allows for communal degradation. (C) Schematic describing stochastic plasmid mutation which relieves burden but does not impact antibiotic resistance. (D) Stochastic simulations of burdensome production in 1x and 2x terminal differentiation architectures with varying $n_{div}$. Mean total production +/- SD of 8 stochastic simulations of 20 consecutive batch growths with 50x dilutions. $\mu_P$=2 h$^{-1}$; 10, 50, 90 percent burden (increasing top to bottom); $K$ = 10$^9$ cells; $k_{MB}$ = $k_{MD}$ 10$^{-6}$ h$^{-1}$; $k_{diff}$ = 0.2, 0.4, 0.6, 0.8, 1, 1.2 h$^{-1}$. Simulations with antibiotic degradation are with plasmid loss rate $k_{PL}$ = 10$^{-4}$ h$^{-1}$; 100 µg/mL antibiotic; MIC = 1.1 µg/mL; and $V_{max}$ = 0, 2.52 x 10$^{-6}$ , and 1.26 x 10$^{-5}$ µg/cell/h (left to right increasing abx deg). Simulations with plasmid mutation were modeled with $k_{PL}$ = $10^{-8}$, $10^{-6}$, $10^{-4}$ $h^{-1}$ (increasing left to right), 0 µg/mL antibiotic, and $V_{max}$ = 0.

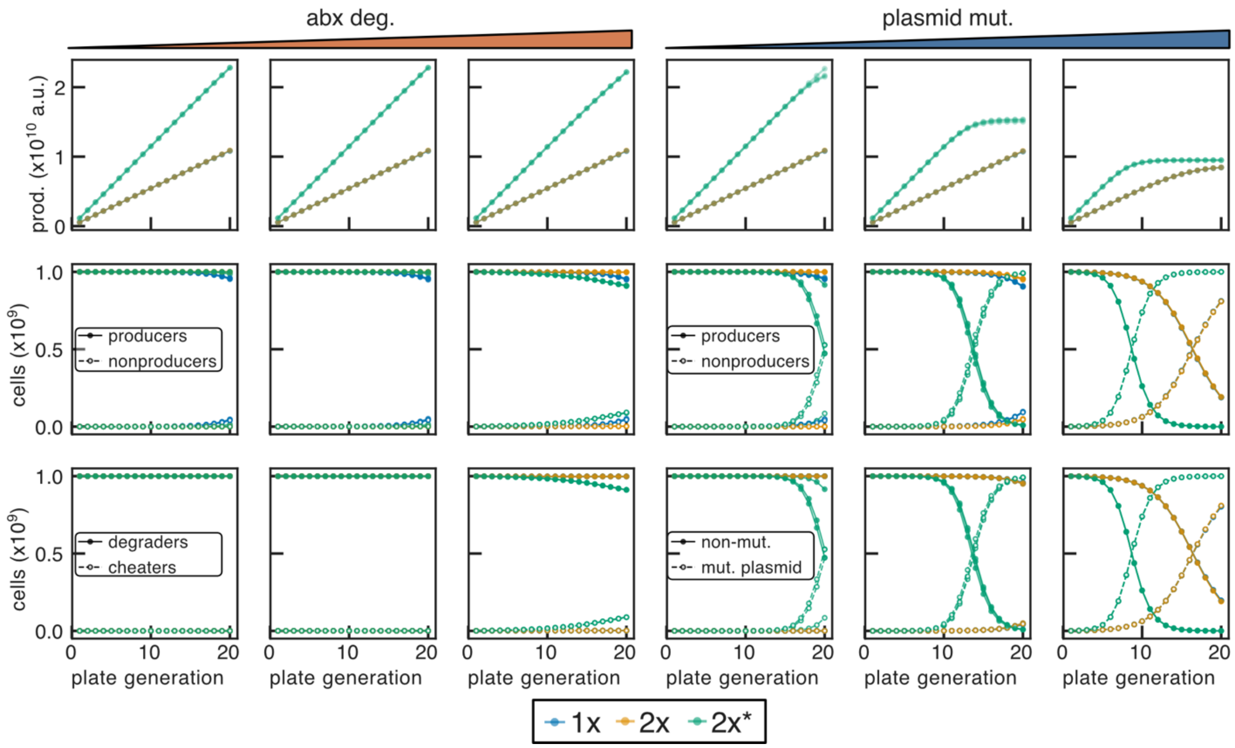


Figure S19: Simulations of 1x and 2x naive circuit architectures with low (10%) burden. Endpoint cumulative production (top); producer/non-producer fractions (middle); fraction retaining (degraders) or having lost (cheaters) the antibiotic resistance/expression plasmid (bottom) for simulations modeling plasmid loss ($k_{PL}$ = $10^{-4}$ $h^{-1}$) with varying rates of antibiotic degradation ($V_{max}$ = 0, 2.52 x $10^{-6}$ , and 1.26 x $10^{-5}$ µg/cell/h left to right); and fraction of cells with non-mutated functional plasmid and mutated non-functional plasmid (bottom) for simulations modeling plasmid mutation with varying rates ($k_{PL}$ = $10^{-8}$, $10^{-6}$, $10^{-4}$ $h^{-1}$ increasing left to right; 0 µg/mL antibiotic, and $V_{max}$ = 0) plotted for 20 consecutive batch growths. 3 of 8 total stochastic simulations plotted for each model. Endpoint production data from all 8 simulations shown in Figure 3.

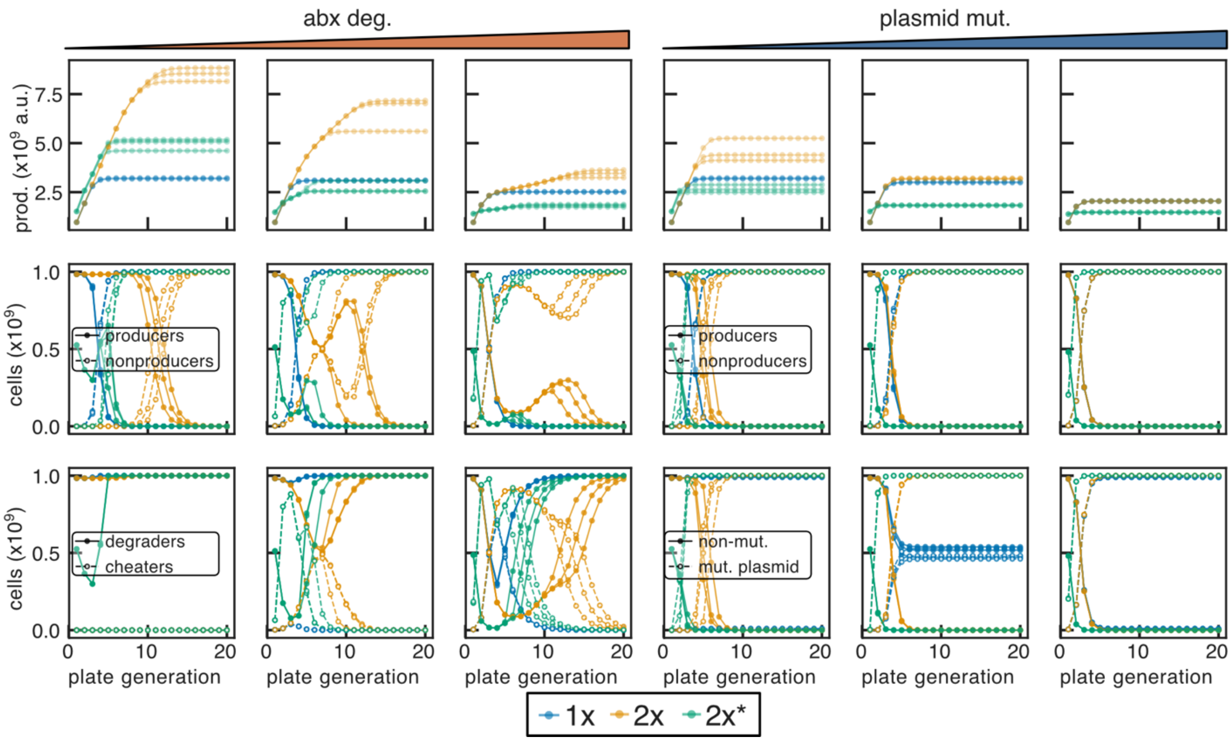


Figure S20: Simulations of 1x and 2x naive circuit architectures with intermediate (50%) burden. Endpoint cumulative production (top); producer/non-producer fractions (middle); fraction retaining (degraders) or having lost (cheaters) the antibiotic resistance/expression plasmid (bottom) for simulations modeling plasmid loss ($k_{PL}$ = $10^{-4}$ $h^{-1}$) with varying rates of antibiotic degradation ($V_{max}$ = 0, 2.52 x $10^{-6}$ , and 1.26 x $10^{-5}$ µg/cell/h left to right); and fraction of cells with non-mutated functional plasmid and mutated non-functional plasmid (bottom) for simulations modeling plasmid mutation with varying rates ($k_{PL}$ = $10^{-8}$, $10^{-6}$, $10^{-4}$ $h^{-1}$ increasing left to right; 0 µg/mL antibiotic, and $V_{max}$ = 0) plotted for 20 consecutive batch growths. 3 of 8 total stochastic simulations plotted for each model. Endpoint production data from all 8 simulations shown in Figure 3.

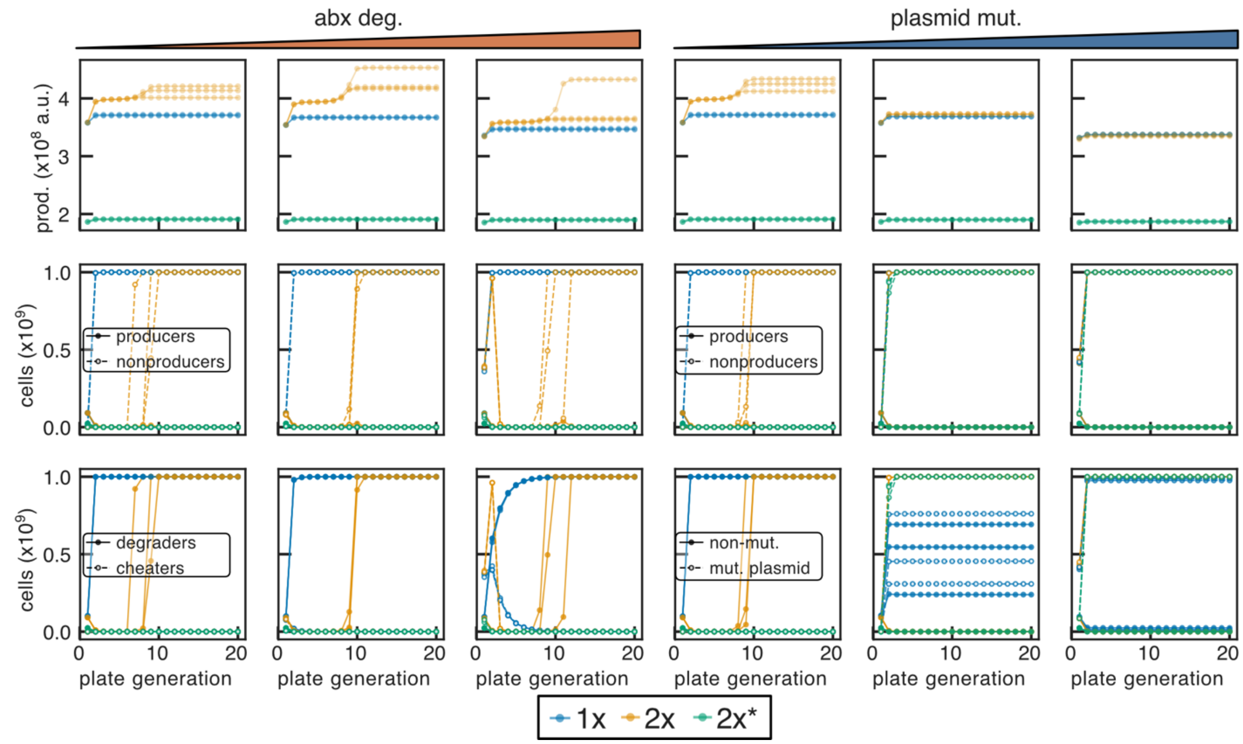


Figure S21: Simulations of 1x and 2x naive circuit architectures with high (90%) burden. Endpoint cumulative production (top); producer/non-producer fractions (middle); fraction retaining (degraders) or having lost (cheaters) the antibiotic resistance/expression plasmid (bottom) for simulations modeling plasmid loss ($k_{PL}$ = $10^{-4}$ $h^{-1}$) with varying rates of antibiotic degradation ($V_{max}$ = 0, 2.52 x $10^{-6}$ , and 1.26 x $10^{-5}$ µg/cell/h left to right); and fraction of cells with non-mutated functional plasmid and mutated non-functional plasmid (bottom) for simulations modeling plasmid mutation with varying rates ($k_{PL}$ = $10^{-8}$, $10^{-6}$, $10^{-4}$ $h^{-1}$ increasing left to right; 0 µg/mL antibiotic, and $V_{max}$ = 0) plotted for 20 consecutive batch growths. 3 of 8 total stochastic simulations plotted for each model. Endpoint production data from all 8 simulations shown in Figure 3.

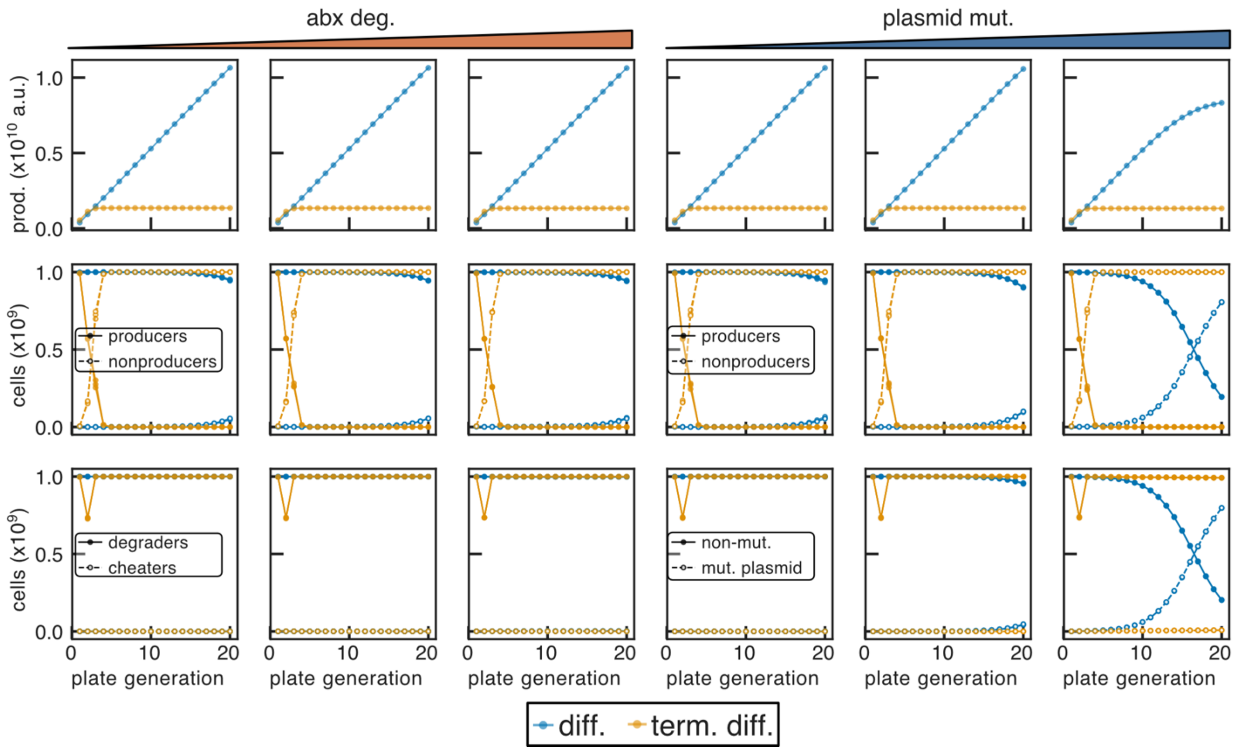


Figure S22: Simulations of 1x differentiation and terminal differentiation architectures with low (10%) burden. Endpoint cumulative production (top); producer/non-producer fractions (middle); fraction retaining (degraders) or having lost (cheaters) the antibiotic resistance/expression plasmid (bottom) for simulations modeling plasmid loss ($k_{PL}$ = $10^{-4}$ $h^{-1}$) with varying rates of antibiotic degradation ($V_{max}$ = 0, 2.52 x $10^{-6}$ , and 1.26 x $10^{-5}$ µg/cell/h left to right); and fraction of cells with non-mutated functional plasmid and mutated non-functional plasmid (bottom) for simulations modeling plasmid mutation with varying rates ($k_{PL}$ = $10^{-8}$, $10^{-6}$, $10^{-4}$ $h^{-1}$ increasing left to right; 0 µg/mL antibiotic, and $V_{max}$ = 0) plotted for 20 consecutive batch growths. Differentiation rate ($k_{diff}$) for all simulations was 0.8 $h^{-1}$; $n_{div}$ = 4 for terminal differentiation. 3 of 8 total stochastic simulations plotted for each model. Endpoint production data from all 8 simulations shown in Figure 3.

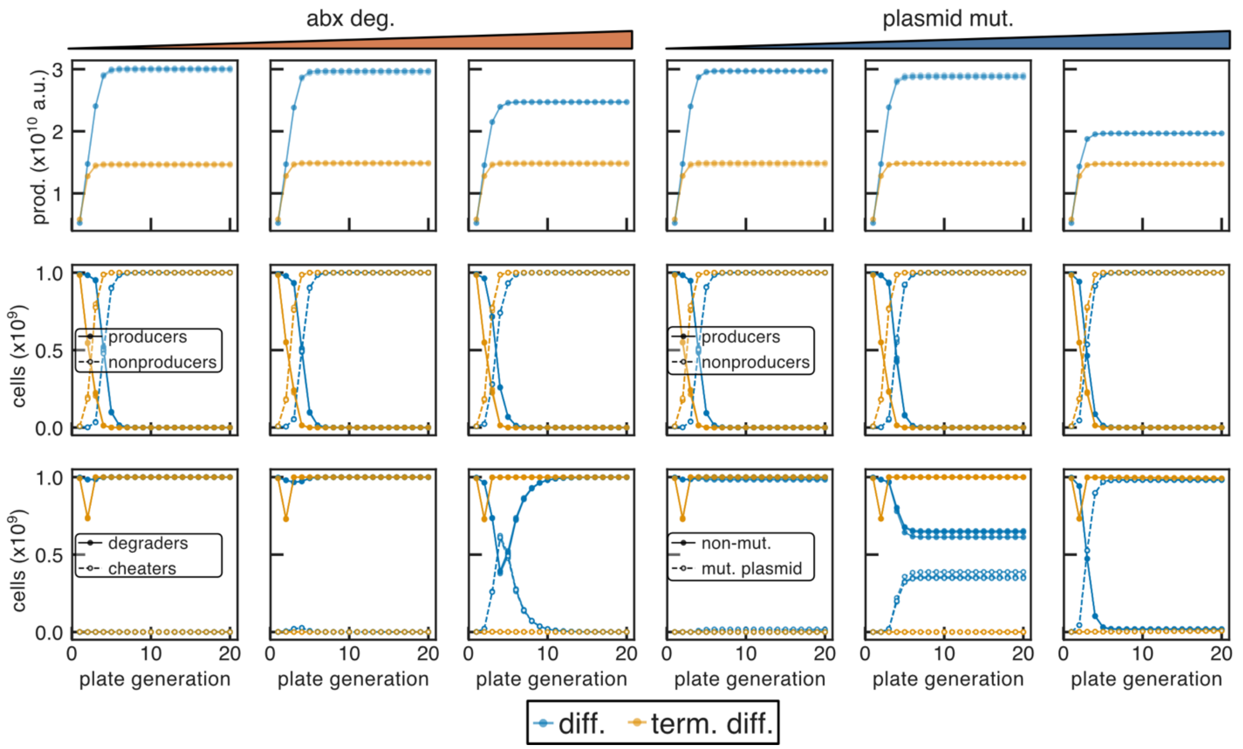


Figure S23: Simulations of 1x differentiation and terminal differentiation architectures with intermediate (50%) burden. Endpoint cumulative production (top); producer/non-producer fractions (middle); fraction retaining (degraders) or having lost (cheaters) the antibiotic resistance/expression plasmid (bottom) for simulations modeling plasmid loss ($k_{PL}$ = $10^{-4}$ $h^{-1}$) with varying rates of antibiotic degradation ($V_{max}$ = 0, 2.52 x $10^{-6}$ , and 1.26 x $10^{-5}$ µg/cell/h left to right); and fraction of cells with non-mutated functional plasmid and mutated non-functional plasmid (bottom) for simulations modeling plasmid mutation with varying rates ($k_{PL}$ = $10^{-8}$, $10^{-6}$, $10^{-4}$ $h^{-1}$ increasing left to right; 0 µg/mL antibiotic, and $V_{max}$ = 0) plotted for 20 consecutive batch growths. Differentiation rate ($k_{diff}$) for all simulations was 0.8 $h^{-1}$; $n_{div}$ = 4 for terminal differentiation. 3 of 8 total stochastic simulations plotted for each model. Endpoint production data from all 8 simulations shown in Figure 3.

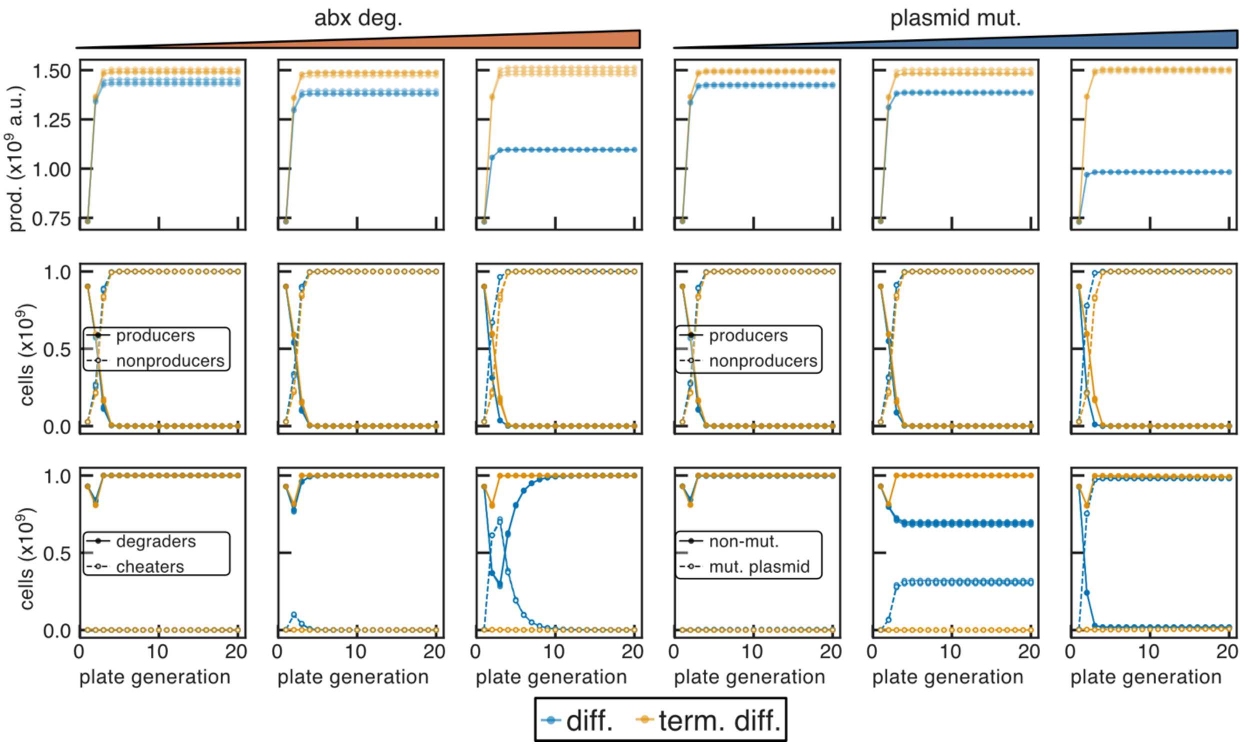


Figure S24: Simulations of 1x differentiation and terminal differentiation architectures with high (90%) burden. Endpoint cumulative production (top); producer/non-producer fractions (middle); fraction retaining (degraders) or having lost (cheaters) the antibiotic resistance/expression plasmid (bottom) for simulations modeling plasmid loss ($k_{PL}$ = $10^{-4}$ $h^{-1}$) with varying rates of antibiotic degradation ($V_{max}$ = 0, 2.52 x $10^{-6}$ , and 1.26 x $10^{-5}$ µg/cell/h left to right); and fraction of cells with non-mutated functional plasmid and mutated non-functional plasmid (bottom) for simulations modeling plasmid mutation with varying rates ($k_{PL}$ = $10^{-8}$, $10^{-6}$, $10^{-4}$ $h^{-1}$ increasing left to right; 0 µg/mL antibiotic, and $V_{max}$ = 0) plotted for 20 consecutive batch growths. Differentiation rate ($k_{diff}$) for all simulations was 0.8 $h^{-1}$; $n_{div}$ = 4 for terminal differentiation. 3 of 8 total stochastic simulations plotted for each model. Endpoint production data from all 8 simulations shown in Figure 3.

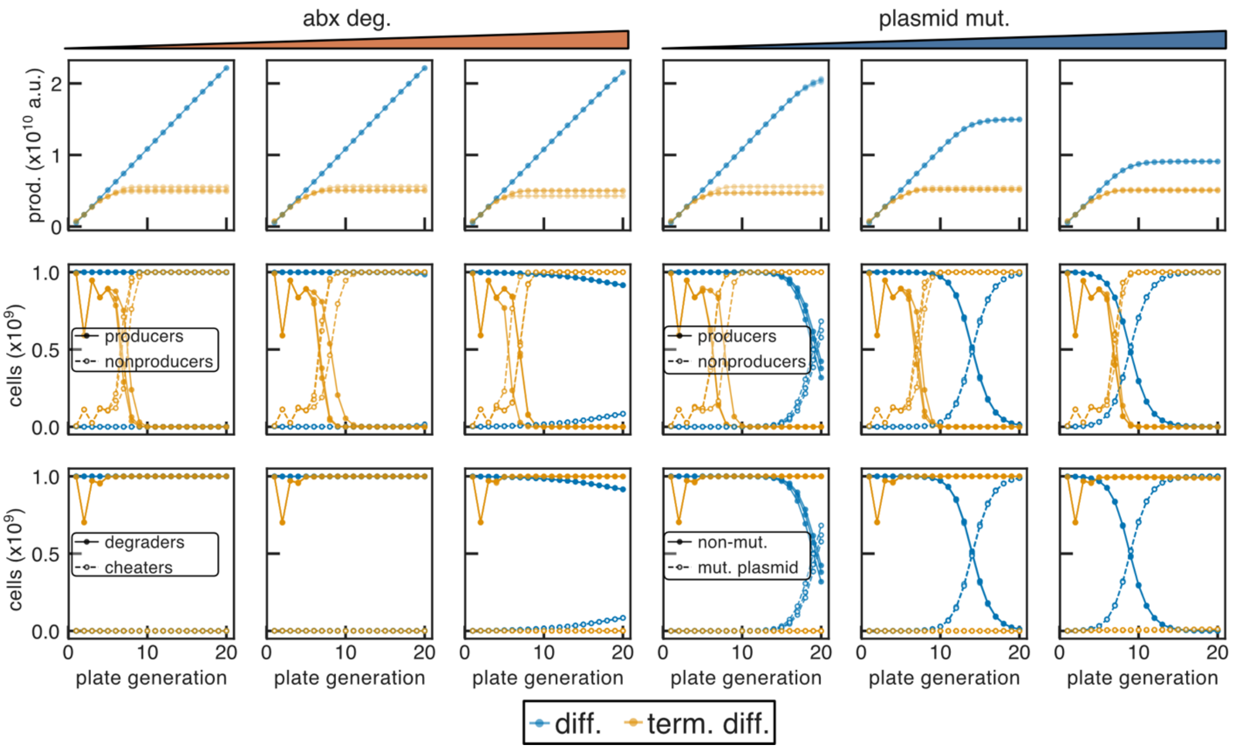


Figure S25: Simulations of 2x differentiation and terminal differentiation architectures with low (10%) burden. Endpoint cumulative production (top); producer/non-producer fractions (middle); fraction retaining (degraders) or having lost (cheaters) the antibiotic resistance/expression plasmid (bottom) for simulations modeling plasmid loss ($k_{PL}$ = $10^{-4}$ $h^{-1}$) with varying rates of antibiotic degradation ($V_{max}$ = 0, 2.52 x $10^{-6}$, and 1.26 x $10^{-5}$ µg/cell/h left to right); and fraction of cells with non-mutated functional plasmid and mutated non-functional plasmid (bottom) for simulations modeling plasmid mutation with varying rates ($k_{PL}$ = $10^{-8}$, $10^{-6}$, $10^{-4}$ $h^{-1}$ increasing left to right; 0 µg/mL antibiotic, and $V_{max}$ = 0) plotted for 20 consecutive batch growths. Differentiation rate ($k_{diff}$) for all simulations was 0.8 $h^{-1}$; $n_{div}$ = 4 for terminal differentiation. 3 of 8 total stochastic simulations plotted for each model. Endpoint production data from all 8 simulations shown in Figure 3.

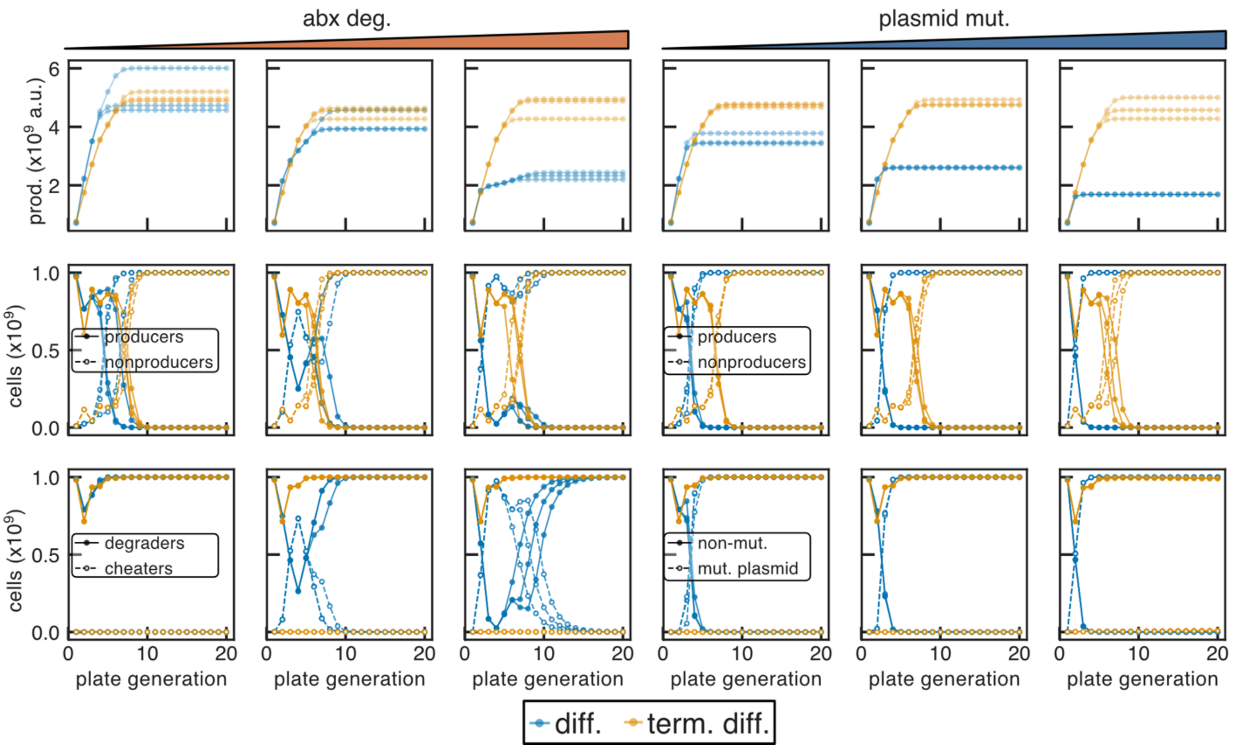


Figure S26: Simulations of 2x differentiation and terminal differentiation architectures with intermediate (50%) burden. Endpoint cumulative production (top); producer/non-producer fractions (middle); fraction retaining (degraders) or having lost (cheaters) the antibiotic resistance/expression plasmid (bottom) for simulations modeling plasmid loss ($k_{PL}$ = 10$^{-4}$ h$^{-1}$) with varying rates of antibiotic degradation ($V_{max}$ = 0, 2.52 x 10$^{-6}$ , and 1.26 x 10$^{-5}$ µg/cell/h left to right); and fraction of cells with non-mutated functional plasmid and mutated non-functional plasmid (bottom) for simulations modeling plasmid mutation with varying rates ($k_{PL}$ = 10$^{-8}$, 10$^{-6}$, 10$^{-4}$ h$^{-1}$ increasing left to right; 0 µg/mL antibiotic, and $V_{max}$ = 0) plotted for 20 consecutive batch growths. Differentiation rate ($k_{diff}$) for all simulations was 0.8 h$^{-1}$; $n_{div}$ = 4 for terminal differentiation. 3 of 8 total stochastic simulations plotted for each model. Endpoint production data from all 8 simulations shown in Figure 3.

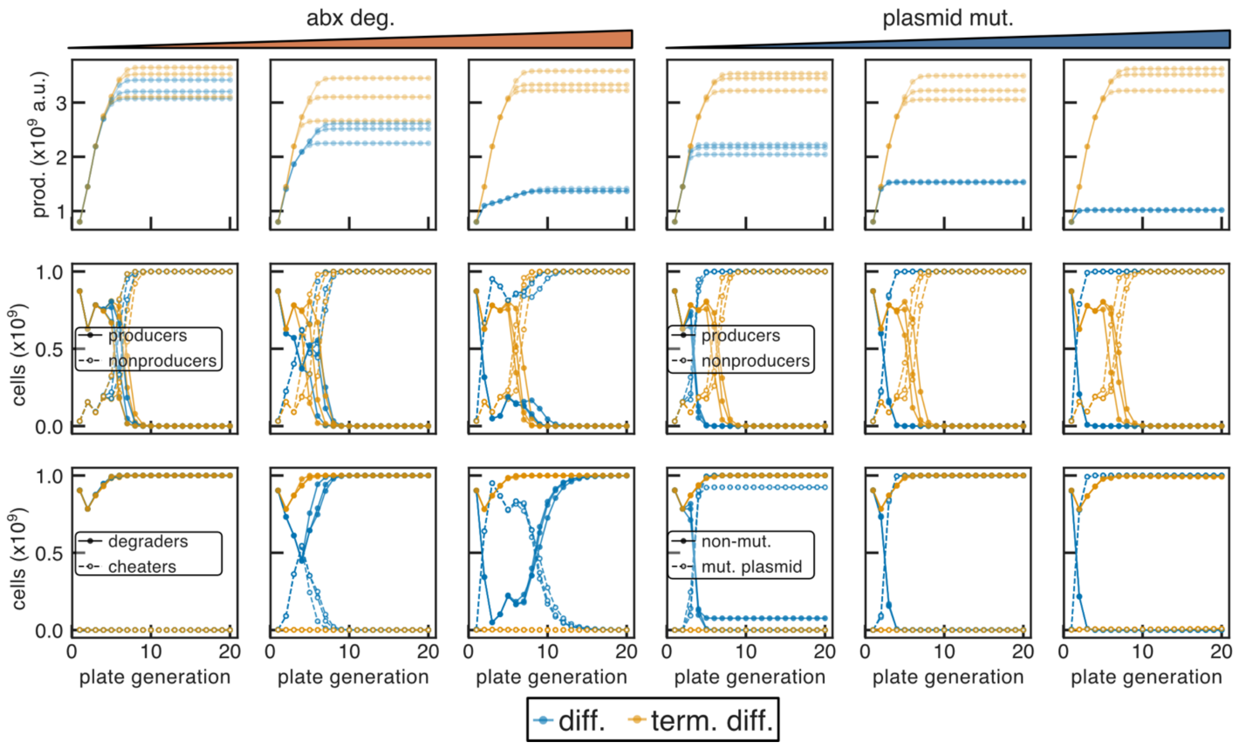


Figure S27: Simulations of 2x differentiation and terminal differentiation architectures with high (90%) burden. Endpoint cumulative production (top); producer/non-producer fractions (middle); fraction retaining (degraders) or having lost (cheaters) the antibiotic resistance/expression plasmid (bottom) for simulations modeling plasmid loss ($k_{PL}$ = $10^{-4}$ $h^{-1}$) with varying rates of antibiotic degradation ($V_{max}$ = 0, 2.52 x $10^{-6}$, and 1.26 x $10^{-5}$ µg/cell/h left to right); and fraction of cells with non-mutated functional plasmid and mutated non-functional plasmid (bottom) for simulations modeling plasmid mutation with varying rates ($k_{PL}$ = $10^{-8}$, $10^{-6}$, $10^{-4}$ $h^{-1}$ increasing left to right; 0 µg/mL antibiotic, and $V_{max}$ = 0) plotted for 20 consecutive batch growths. Differentiation rate ($k_{diff}$) for all simulations was 0.8 $h^{-1}$; $n_{div}$ = 4 for terminal differentiation. 3 of 8 total stochastic simulations plotted for each model. Endpoint production data from all 8 simulations shown in Figure 3.

| STRAIN | SITE | INTEGRATION |
|---|---|---|
| **eRWnaive1x** | P21 | POS1-$P_{Tac}$-ARL-T7(attL)RNAP-T12m-UNS3<br>$P_{LacI}$-LacI$^{AM}$-T2m-POSX |
| **eRWnaive2x** | P21 | POS1- $P_{Tac}$-ARL-T7(attL)RNAP-T12m-UNS3<br>$P_{LacI}$-LacI$^{AM}$-T2m-POSX |
| | HK022 | POS1-$P_{Tac}$-ARL-T7(attL)RNAP-T12m-UNS3<br>$P_{LacI}$-LacI$^{AM}$-T2m-POSX |
| **eRWdiff1x** | P21 | POS1-$P_{Tac}$-ARL-T7RNAP(L)-attB-T14m-UNS3<br>**$P_{LasAM}$-B0034-pir-T2m**-UNS4<br>T13m-attP-T7RNAP(R)-UNS5<br>$P_{4d}$-B0034-NahR$^{AM}$-BCD2-LasR$^{AM}$-T15m-UNS6<br>$P_{LacI}$-LacI$^{AM}$-T17m-POSX |
| | Φ186 Primary | POS1-$P_{SalTTC}$-B0034-Bxb1LAA-T2m-POSX |
| | Φ186 secondary | POS1-$P_{SalTTC}$-B0034-Bxb1LAA-T2m-POSX |
| **eRWdiff2x** | P21 | POS1-$P_{Tac}$-ARL-T7RNAP(L)-attB-T14m-UNS3<br>**$P_{LasAM}$-B0034-pirN:cfaN-T2m**-UNS4<br>T13m-attP-T7RNAP(R)-UNS5<br>$P_{4d}$-B0034-NahR$^{AM}$-BCD2-LasR$^{AM}$-T15m-UNS6<br>$P_{LacI}$-LacI$^{AM}$-T17m-POSX |
| | HK022 | POS1-$P_{Tac}$-ARL-T7RNAP(L)-attB-T14m-UNS3<br>**$P_{LasAM}$-B0034-cfaC:pirC-T2m**-UNS4<br>T13m-attP-T7RNAP(R)-UNS5f<br>$P_{4d}$-B0034-NahR$^{AM}$-BCD2-LasR$^{AM}$-T15m-UNS6<br>$P_{LacI}$-LacI$^{AM}$-T17m-POSX |
| | Φ186 Primary | POS1-$P_{SalTTC}$-B0034-Bxb1LAA-T2m-POSX |
| | Φ186 secondary | POS1-$P_{SalTTC}$-B0034-Bxb1LAA-T2m-POSX |

Table S2. Summary of strains used in this study. JS006 is parental strain for all strains. **sequences are reversed**. POS1, POSX,

#### eRWnaive1x

| Site | Sequence |
| --- | --- |
| P21<br>(T) | TAGAACTACCACCTGACCGGCCAGCCTACCCCAAAACTACCGAGGGGTTTTGATAACTTTTTTGATAAAATTTTGATAACCGTTAGCAGCCTAACAAAAAACAACGGGAGCGTTACGCTCCCGT<br>TAATAAATTTAACAAACTACGGCATTACATGTTTTCGATGATCGCGTCACCAAACTCTGAACATTTCAGCAGTTTAGCGCCATCCATCAGACGCTCGAAGTCATAGGTTACGGTTTTCGCGTTGAT<br>TGCGCCTTCCATACCTTTAACAATTAAGTCAGCCGCTTCGGTCCAACCCATGTGGCGCAGCAGTATATAAAAATTATGTATTCAACTCGCTGATATTTAATCATAAATATTATTTTCTTGTCTTTGT<br>ATTAGCCAAAATCAACTTTATTCAACCGTTTGATTTCATTGACGTTCGTTTTTATTTTGGGGAAAGATTTTCTCTACCATTTATATCAAAATTACAAGTCTACATTTTATTAACATCTCAATACATGT<br>GTGGCCTTGCAAGCCATTGCGAGGCCTTATGCTAGCTTCTTCGTCTGTTTCTACTGGTATTGGCACAAACCTGATTCCAATTTGAGCAAGGCTATGTGCCATCTCGATACTCGTTCTTAACTCAAC<br>AGAAGATGCTTTGTGCATACAGCCCCTCGTTTATTATTTATCTCCTCAGCCAGCCGCTGTGCTTTCAGTGGATTTCGGATAACAGAAAGGCCGGGAAATACCCAGCCTCGCTTTGTAACGGAGTA<br>GAGACGAAAGTGATTGCGCCTACCCGGATATTATCGTGAGGATGCGTCATCGCCATTAATTCACTGATCAGTGATAAGCTGTCAAACATGAGAATTGATCCGGCTGCCTCGCGCGTTTCGGTGA<br>TGACGGTGAAAACCTCTGACACATGCAGCTCCCGGAGACGGTCACAGCTTGTCTGTAAGCGGATGCCGGGAGCAGACAAGCCCGTCAGGGCGCGTCAGCGGGTGTTGGCGGGTGTCGGGGC<br>GCAGCCATGACCCAGTCACGTAGCGATAGCGGAGTGTATGCTGCACATGACATTAACCTATAAAAATAGGCGTATCACGAGGCCCTTTCGTCTTCAAGAATTAATTCCCAATTCCCCAGGCATCA<br>AATAAAACGAAAGGCTCAGTCGAAAGACTGGGCCTTTCGTTTTATCTGTTGTTTGTCGGTGAACGCTCTCCTGAGTAGGACAAATCCGCCGGGAGCGGATTTGAACGTTGCGAAGCAACGGCC<br>CGGAGGGTGGCGGGCAGGACGCCCGCCATAAACTGCCAGGAATTAATTCCCCAGGCATCAAATAAAACGAAAGGCTCAGTCGAAAGACTGGGCCTTTCGTTTTATCTGTTGTTTGTCGGTGAA<br>CGCTCTCCTGAGTAGGACAAATCCGCCGGGAGCGGATTTGAACGTTGCGAAGCAACGGCCCGGAGGGTGGCGGGCAGGACGCCCGCCATAAACTGCCAGGAATTAATTCCCCAGGCATCAAA<br>TAAAACGAAAGGCTCAGTCGAAAGACTGGGCCTTTCGTTTTATCTGTTGTTTGTCGGTGAACGCTCTCCTGAGTAGGACAAATCCGCCGGGAGCGGATTTGAACGTTGCGAAGCAACGGCCCG<br>GAGGGTGGCGGGCAGGACGCCCGCCATAAACT***GCCAGGAATTGGGGATCGGAATTCGAGCTCGGTACCCGGG***GGAGAACGATCGTTGGCTGTGTTGACAATTAATCATCGGCTCGTATAAT<br>GTGTGGAATTGTGAGCGCTCACAATTTACTGAAAGATGCGATTTACTAAATGAACACGATTAACATCGCTAAGAACGACTTCTCTGACATCGAACTGGCTGCTATCCCGTTCAACACTCTGGCTG<br>ACCATTACGGTGAGCGTTTAGCTCGCGAACAGTTGGCCCTTGAGCATGAGTCTTACGAGATGGGTGAAGCACGCTTCCGCAAGATGTTTGAGCGTCAACTTAAAGCTGGTGAGGTTGCGGATA<br>ACGCTGCCGCCAAGCCTCTCATCACTACCCTACTCCCTAAGATGATTGCACGCATCAACGACTGGTTTGAGGAAGTGAAAGCTAAGCGCGGCAAGCGCCCGACAGCCTTCCAGTTCCTGCAAGA<br>AATCAAGCCGGAAGCCGTAGCGTACATCACCATTAAGACCACTCTGGCTTGCCTAACCAGTGCTGACAATACAACCGTTCAGGCTGTAGCAAGCGCAATCGGTCGGGCCATTGAGGACGAGGC<br>TCGCTTCGGTCGTATCCGTGACCTTGAAGCTAAGCACTTCAAGAAAAACGTTGAGGAACAACTCAACAAGCGCGTAGGGCACGTCTACAAGGGCTTGTCGACGACGGCGTGCTCAGTGGTGTA<br>CGGTACAAACCAGGGTTCTAAAGCATTTATGCAAGTTGTCGAGGCTGACATGCTCTCTAAGGGTCTACTCGGTGGCGAGGCGTGGTCTTCGTGGCATAAGGAAGACTCTATTCATGTAGGAGT<br>ACGCTGCATCGAGATGCTCATTGAGTCAACCGGAATGGTTAGCTTACACCGCCAAAATGCTGGCGTAGTAGGTCAAGACTCTGAGACTATCGAACTCGCACCTGAATACGCTGAGGCTATCGCA<br>ACCCGTGCAGGTGCGCTGGCTGGCATCTCTCCGATGTTCCAACCTTGCGTAGTTCCTCCTAAGCCGTGGACTGGCATTACTGGTGGTGGCTATTGGGCTAACGGTCGTCGTCCTCTGGCGCTGG<br>TGCGTACTCACAGTAAGAAAGCACTGATGCGCTACGAAGACGTTTACATGCCTGAGGTGTACAAAGCGATTAACATTGCGCAAAACACCGCATGGAAAATCAACAAGAAAGTCCTAGCGGTCG<br>CCAACGTAATCACCAAGTGGAAGCATTGTCCGGTCGAGGACATCCCTGCGATTGAGCGTGAAGAACTCCCGATGAAACCGGAAGACATCGACATGAATCCTGAGGCTCTCACCGCGTGGAAAC<br>GTGCTGCCGCTGCTGTGTACCGCAAGGACAAGGCTCGCAAGTCTCGCCGTATCAGCCTTGAGTTCATGCTTGAGCAAGCCAATAAGTTTGCTAACCATAAGGCCATCTGGTTCCCTTACAACAT<br>GGACTGGCGCGGTCGTGTTTACGCTGTGTCAATGTTCAACCCGCAAGGTAACGATATGACCAAAGGACTGCTTACGCTGGCGAAAGGTAAACCAATCGGTAAGGAAGGTTACTACTGGCTGAA<br>AATCCACGGTGCAAACTGTGCGGGTGTCGATAAGGTTCCGTTCCCTGAGCGCATCAAGTTCATTGAGGAAAACCACGAGAACATCATGGCTTGCGCTAAGTCTCCACTGGAGAACACTTGGTG<br>GGCTGAGCAAGATTCTCCGTTCTGCTTCCTTGCGTTCTGCTTTGAGTACGCTGGGGTACAGCACCACGGCCTGAGCTATAACTGCTCCCTTCCGCTGGCGTTTGACGGGTCTTGCTCTGGCATCC<br>AGCACTTCTCCGCGATGCTCCGAGATGAGGTAGGTGGTCGCGCGGTTAACTTGCTTCCTAGTGAAACCGTTCAGGACATCTACGGGATTGTTGCTAAGAAAGTCAACGAGATTCTACAAGCAG<br>ACGCAATCAATGGGACCGATAACGAAGTAGTTACCGTGACCGATGAGAACACTGGTGAAATCTCTGAGAAAGTCAAGCTGGGCACTAAGGCACTGGCTGGTCAATGGCTGGCTTACGGTGTT<br>ACTCGCAGTGTGACTAAGCGTTCAGTCATGACGCTGGCTTACGGGTCCAAAGAGTTCGGCTTCCGTCAACAAGTGCTGGAAGATACCATTCAGCCAGCTATTGATTCCGGCAAGGGTCTGATGT<br>TCACTCAGCCGAATCAGGCTGCTGGATACATGGCTAAGCTGATTTGGGAATCTGTGAGCGTGACGGTGGTAGCTGCGGTTGAAGCAATGAACTGGCTTAAGTCTGCTGCTAAGCTGCTGGCTG<br>CTGAGGTCAAAGATAAGAAGACTGGAGAGATTCTTCGCAAGCGTTGCGCTGTGCATTGGGTAACTCCTGATGGTTTCCCTGTGTGGCAGGAATACAAGAAGCCTATTCAGACGCGCTTGAACC<br>TGATGTTCCTCGGTCAGTTCCGCTTACAGCCTACCATTAACACCAACAAAGATAGCGAGATTGATGCACACAAACAGGAGTCTGGTATCGCTCCTAACTTTGTACACAGCCAAGACGGTAGCCA<br>CCTTCGTAAGACTGTAGTGTGGGCACACGAGAAGTACGGAATCGAATCTTTTGCACTGATTCACGACTCCTTCGGTACCATTCCGGCTGACGCTGCGAACCTGTTCAAAGCAGTGCGCGAAACT<br>ATGGTTGACACATATGAGTCTTGTGATGTACTGGCTGATTTCTACGACCAGTTCGCTGACCAGTTGCACGAGTCTCAATTGGACAAAATGCCAGCACTTCCGGCTAAAGGTAACTTGAACCTCCG<br>TGACATCTTAGAGTCGGACTTCGCGTTCGCGTAATAAAGGTTTCAGCCAAAAAACTTAAGACCGCCGGTCTTGTCCACTACCTTGCAGTAATGCGGTGGACAGGATCGGCGGTTTTCTTTTCTCT<br>TCTCAAGCTT***GCACT_AAGGTCCTCAATCGCACTGGAAACATCAAGGTCG***GGAGGACACCATCGAATGGCGCAAAACCTTTCGCGGTATGGCATGATAGCGCCCGGAAGAGAGTCAATTCAG<br>GGTGGTGAATAATGAAACCAGTAACGTTATACGATGTCGCAGAGTATGCCGGTGTCTCTTATATGACCGTTTCCCGCGTGGTGAACCAGGCCAGCCACGTTTCTGCGAAAACGCGGGAAAAAG<br>TGGAAGCGGCGATGGTGGAGCTGAATTACATTCCCAACCGCGTGGCACAACAACTGGCGGGCAAACAGTCGTTGCTGATTGGCGTTGCCACCTCCAGTCTGGCCCTGCACGCGCCGTCGCAAA<br>TTGTCGCGGCGATTAAATCTCGCGCCGATCAACTGGGTGCCAGCGTGGTGGTGTCGATGGTAGAACGAAGCGGCGTCGAAGCCTGTAAAGCGGCGGTGCACAATCTTCTCGCGCAACGCGTC<br>AGTGGGCTGATCATTAACTATCCGCTGGATGACCAGGATGCCATTGCTGTGGAAGCTGCCTGCACTAATGTTCCGGCGTTATTTCTTGATGTCTCTGACCAGACACCCATCAACAGTATTATTTA<br>CTCCCATGAGGACGGTACGCGACTGGGCGTGGAGCATCTGGTCGCATTGGGTCACCAGCAAATCGCGCTGTTAGCGGGCCCATTAAGTTCTGTCTCGGCGCGTCTGCGTCTGGCTGGCTGGCA<br>TAAATATCTCACTCGCAATCAAATTCAGCCGATAGCGGAACGGGAAGGCGACTGGAGTGCCATGTCCGGTTTTCAACAAACCATGCAAATGCTGAATGAGGGCATCGTTCCCACTGCGATGCT<br>GGTTGCCAACGATCAGATGGCGCTGGGCGCAATGCGCGCCATTACCGAGTCCGGGCTGCGCGTTGGTGCGGATATCTCGGTAGTGGGATACGACGATACCGAAGATAGCTCATGTTATATCCC<br>GCCGTTAACCACCATCAAACAGGATTTTCGCCTGCTGGGGCAAACCAGCGTGGACCGCTTGCTGCAACTCTCTCAGGGCCAGGCGGTGAAGGGCAATCAGCTGTTGCCAGTCTCACTGGTGAA<br>AAGAAAAACCACCCTGGCGCCCAATACGCAAACCGCCTCTCCCCGCGCGTTGGCCGATTCATTAATGCAGCTGGCACGACAGGTTTCCCGACTGGAAAGCGGGCAGTGAGGTCTAACGCATGA<br>GAAAGCCCCCGGAAGATCACCTTCCGGGGGCTTTTTTATTGCGCCCTTGAGAGCTT***TAGCGGCCGCTGCAGGCATGCCTCGAGATGCATGGCGCCT***AACCTAAACTGACAGGCATCAAATTAA<br>GCAGAAGGCCATCCTGACGGATGGCCTTTTTGCGTTTCGAACAATTGAAAAAACCTCGCGCCTTACCTGTTGAGTAATAGTCAAAAGCCTCCGGTCGGAGGCTTTTGACTTTCTGCTTACTGAAT<br>TTCGGTGGTGCCGTTAATTAACCGGTGGGCCCTCATGATAATAATGGTTTCTTAGACGTCCGAAGTTCCTATTCTCTAGAAAGTATAGGAACTTCCATATGCCATGGAATGACCGACCGATACAA<br>TGATGATCGTGGTAAAGACTGGATTATCGTAGATATCAAAACAGCATAGAAAATAGCCAGTTTTGGGGAAGGGTTTTGGGGAAAGTTTTGGGGAAGATTTTACATCATCATAAAACAACGGGC<br>GTGTTATACGCCCGTTATAATATTTAACACATGTGGTAATTACATATTTTTGATGATCGCGTCACCAAACTCGCTGCATTTCAGCAGTTTAGCGCCTTCCATCAGACGTTCGAAGTCATAGGTTAC<br>GGTCTTGGCGTTGATTGCGCCTTCCATACCTTTAACAATCAGGTCTGCGGCTTCAGTCCAGCCCATATGGCGCAGCATCATCTCAGCGGAGAGAATAATAGAGCCAGGATTTACTTTGTCCTGAC<br>CGGCATATTTCGGCGCAGTACCGTGGGTGGCTTCAAACAGGGCGCATTCGTCACCGATGTTTGCACCAGGGGCGATACCGATACCGCCAACCTGCGCTGCCAGGGCGTCAGAAATGTAGTCAC<br>CGTTCAGGTTCATACAGGCGAT |

Table S3. Naive 1X strain with IPTG inducible T7 RNAP. Promoters, RBSs, coding sequences, inserted bases in T7 RNAP CDS, terminators, integrase attachment sites, ***Gibson homologies***, genomic pOSIP primer binding sites. Gap in expected UNS sequence for Gibson indicated by “_”.

#### eRWnaive2x

| Site | Sequence |
|---|---|
| P21<br>(T) | TAGAACTACCACCTGACCGGCCAGCCTACCCCAAAACTACCGAGGGGTTTTGATAACTTTTTTGATAAAATTTTGATAACCGTTAGCAGCCTAACAAAAAACAACGGGAGCGTTACGCTCCCGT<br>TAATAAATTTAACAAACTACGGCATTACATGTTTTCGATGATCGCGTCACCAAACTCTGAACATTTCAGCAGTTTAGCGCCATCCATCAGACGCTCGAAGTCATAGGTTACGGTTTTCGCGTTGAT<br>TGCGCCTTCCATACCTTTAACAATTAAGTCAGCCGCTTCGGTCCAACCCATGTGGCGCAGCAGTATATAAAAATTATGTATTCAACTCGCTGATATTTAATCATAAATATTATTTTCTTGTCTTTGT<br>ATTAGCCAAAATCAACTTTATTCAACCGTTTGATTTCATTGACGTTCGTTTTTATTTTGGGGAAAGATTTTCTCTACCATTTATATCAAAATTACAAGTCTACATTTTATTAACATCTCAATACATGT<br>GTGGCCTTGCAAGCCATTGCGAGGCCTTATGCTAGCTTCTTCGTCTGTTTCTACTGGTATTGGCACAAACCTGATTCCAATTTGAGCAAGGCTATGTGCCATCTCGATACTCGTTCTTAACTCAAC<br>AGAAGATGCTTTGTGCATACAGCCCCTCGTTTATTATTTATCTCCTCAGCCAGCCGCTGTGCTTTCAGTGGATTTCGGATAACAGAAAGGCCGGGAAATACCCAGCCTCGCTTTGTAACGGAGTA<br>GAGACGAAAGTGATTGCGCCTACCCGGATATTATCGTGAGGATGCGTCATCGCCATTAATTCACTGATCAGTGATAAGCTGTCAAACATGAGAATTGATCCGGCTGCCTCGCGCGTTTCGGTGA<br>TGACGGTGAAAACCTCTGACACATGCAGCTCCCGGAGACGGTCACAGCTTGTCTGTAAGCGGATGCCGGGAGCAGACAAGCCCGTCAGGGCGCGTCAGCGGGTGTTGGCGGGTGTCGGGGC<br>GCAGCCATGACCCAGTCACGTAGCGATAGCGGAGTGTATGCTGCACATGACATTAACCTATAAAAATAGGCGTATCACGAGGCCCTTTCGTCTTCAAGAATTAATTCCCAATTCCCCAGGCATCA<br>AATAAAACGAAAGGCTCAGTCGAAAGACTGGGCCTTTCGTTTTATCTGTTGTTTGTCGGTGAACGCTCTCCTGAGTAGGACAAATCCGCCGGGAGCGGATTTGAACGTTGCGAAGCAACGGCC<br>CGGAGGGTGGCGGGCAGGACGCCCGCCATAAACTGCCAGGAATTAATTCCCCAGGCATCAAATAAAACGAAAGGCTCAGTCGAAAGACTGGGCCTTTCGTTTTATCTGTTGTTTGTCGGTGAA<br>CGCTCTCCTGAGTAGGACAAATCCGCCGGGAGCGGATTTGAACGTTGCGAAGCAACGGCCCGGAGGGTGGCGGGCAGGACGCCCGCCATAAACTGCCAGGAATTAATTCCCCAGGCATCAAA<br>TAAAACGAAAGGCTCAGTCGAAAGACTGGGCCTTTCGTTTTATCTGTTGTTTGTCGGTGAACGCTCTCCTGAGTAGGACAAATCCGCCGGGAGCGGATTTGAACGTTGCGAAGCAACGGCCCG<br>GAGGGTGGCGGGCAGGACGCCCGCCATAAACT***GCCAGGAATTGGGGATCGGAATTCGAGCTCGGTACCCGGG***GGAGAACGATCGTTGGCTGTGTTGACAATTAATCATCGGCTCGTATAAT<br>GTGTGGAATTGTGAGCGCTCACAATTTACTGAAAGATGCGATTTACTAAATGAACACGATTAACATCGCTAAGAACGACTTCTCTGACATCGAACTGGCTGCTATCCCGTTCAACACTCTGGCTG<br>ACCATTACGGTGAGCGTTTAGCTCGCGAACAGTTGGCCCTTGAGCATGAGTCTTACGAGATGGGTGAAGCACGCTTCCGCAAGATGTTTGAGCGTCAACTTAAAGCTGGTGAGGTTGCGGATA<br>ACGCTGCCGCCAAGCCTCTCATCACTACCCTACTCCCTAAGATGATTGCACGCATCAACGACTGGTTTGAGGAAGTGAAAGCTAAGCGCGGCAAGCGCCCGACAGCCTTCCAGTTCCTGCAAGA<br>AATCAAGCCGGAAGCCGTAGCGTACATCACCATTAAGACCACTCTGGCTTGCCTAACCAGTGCTGACAATACAACCGTTCAGGCTGTAGCAAGCGCAATCGGTCGGGCCATTGAGGACGAGGC<br>TCGCTTCGGTCGTATCCGTGACCTTGAAGCTAAGCACTTCAAGAAAAACGTTGAGGAACAACTCAACAAGCGCGTAGGGCACGTCTACAAGGGCTTGTCGACGACGGCGTGCTCAGTGGTGTA<br>CGGTACAAACCAGGGTTCTAAAGCATTTATGCAAGTTGTCGAGGCTGACATGCTCTCTAAGGGTCTACTCGGTGGCGAGGCGTGGTCTTCGTGGCATAAGGAAGACTCTATTCATGTAGGAGT<br>ACGCTGCATCGAGATGCTCATTGAGTCAACCGGAATGGTTAGCTTACACCGCCAAAATGCTGGCGTAGTAGGTCAAGACTCTGAGACTATCGAACTCGCACCTGAATACGCTGAGGCTATCGCA<br>ACCCGTGCAGGTGCGCTGGCTGGCATCTCTCCGATGTTCCAACCTTGCGTAGTTCCTCCTAAGCCGTGGACTGGCATTACTGGTGGTGGCTATTGGGCTAACGGTCGTCGTCCTCTGGCGCTGG<br>TGCGTACTCACAGTAAGAAAGCACTGATGCGCTACGAAGACGTTTACATGCCTGAGGTGTACAAAGCGATTAACATTGCGCAAAACACCGCATGGAAAATCAACAAGAAAGTCCTAGCGGTCG<br>CCAACGTAATCACCAAGTGGAAGCATTGTCCGGTCGAGGACATCCCTGCGATTGAGCGTGAAGAACTCCCGATGAAACCGGAAGACATCGACATGAATCCTGAGGCTCTCACCGCGTGGAAAC<br>GTGCTGCCGCTGCTGTGTACCGCAAGGACAAGGCTCGCAAGTCTCGCCGTATCAGCCTTGAGTTCATGCTTGAGCAAGCCAATAAGTTTGCTAACCATAAGGCCATCTGGTTCCCTTACAACAT<br>GGACTGGCGCGGTCGTGTTTACGCTGTGTCAATGTTCAACCCGCAAGGTAACGATATGACCAAAGGACTGCTTACGCTGGCGAAAGGTAAACCAATCGGTAAGGAAGGTTACTACTGGCTGAA<br>AATCCACGGTGCAAACTGTGCGGGTGTCGATAAGGTTCCGTTCCCTGAGCGCATCAAGTTCATTGAGGAAAACCACGAGAACATCATGGCTTGCGCTAAGTCTCCACTGGAGAACACTTGGTG<br>GGCTGAGCAAGATTCTCCGTTCTGCTTCCTTGCGTTCTGCTTTGAGTACGCTGGGGTACAGCACCACGGCCTGAGCTATAACTGCTCCCTTCCGCTGGCGTTTGACGGGTCTTGCTCTGGCATCC<br>AGCACTTCTCCGCGATGCTCCGAGATGAGGTAGGTGGTCGCGCGGTTAACTTGCTTCCTAGTGAAACCGTTCAGGACATCTACGGGATTGTTGCTAAGAAAGTCAACGAGATTCTACAAGCAG<br>ACGCAATCAATGGGACCGATAACGAAGTAGTTACCGTGACCGATGAGAACACTGGTGAAATCTCTGAGAAAGTCAAGCTGGGCACTAAGGCACTGGCTGGTCAATGGCTGGCTTACGGTGTT<br>ACTCGCAGTGTGACTAAGCGTTCAGTCATGACGCTGGCTTACGGGTCCAAAGAGTTCGGCTTCCGTCAACAAGTGCTGGAAGATACCATTCAGCCAGCTATTGATTCCGGCAAGGGTCTGATGT<br>TCACTCAGCCGAATCAGGCTGCTGGATACATGGCTAAGCTGATTTGGGAATCTGTGAGCGTGACGGTGGTAGCTGCGGTTGAAGCAATGAACTGGCTTAAGTCTGCTGCTAAGCTGCTGGCTG<br>CTGAGGTCAAAGATAAGAAGACTGGAGAGATTCTTCGCAAGCGTTGCGCTGTGCATTGGGTAACTCCTGATGGTTTCCCTGTGTGGCAGGAATACAAGAAGCCTATTCAGACGCGCTTGAACC<br>TGATGTTCCTCGGTCAGTTCCGCTTACAGCCTACCATTAACACCAACAAAGATAGCGAGATTGATGCACACAAACAGGAGTCTGGTATCGCTCCTAACTTTGTACACAGCCAAGACGGTAGCCA<br>CCTTCGTAAGACTGTAGTGTGGGCACACGAGAAGTACGGAATCGAATCTTTTGCACTGATTCACGACTCCTTCGGTACCATTCCGGCTGACGCTGCGAACCTGTTCAAAGCAGTGCGCGAAACT<br>ATGGTTGACACATATGAGTCTTGTGATGTACTGGCTGATTTCTACGACCAGTTCGCTGACCAGTTGCACGAGTCTCAATTGGACAAAATGCCAGCACTTCCGGCTAAAGGTAACTTGAACCTCCG<br>TGACATCTTAGAGTCGGACTTCGCGTTCGCGTAATAAAGGTTTCAGCCAAAAAACTTAAGACCGCCGGTCTTGTCCACTACCTTGCAGTAATGCGGTGGACAGGATCGGCGGTTTTCTTTTCTCT<br>TCTCAAGCTT***GCACT_AAGGTCCTCAATCGCACTGGAAACATCAAGGTCG***GGAGGACACCATCGAATGGCGCAAAACCTTTCGCGGTATGGCATGATAGCGCCCGGAAGAGAGTCAATTCAG<br>GGTGGTGAATAATGAAACCAGTAACGTTATACGATGTCGCAGAGTATGCCGGTGTCTCTTATATGACCGTTTCCCGCGTGGTGAACCAGGCCAGCCACGTTTCTGCGAAAACGCGGGAAAAAG<br>TGGAAGCGGCGATGGTGGAGCTGAATTACATTCCCAACCGCGTGGCACAACAACTGGCGGGCAAACAGTCGTTGCTGATTGGCGTTGCCACCTCCAGTCTGGCCCTGCACGCGCCGTCGCAAA<br>TTGTCGCGGCGATTAAATCTCGCGCCGATCAACTGGGTGCCAGCGTGGTGGTGTCGATGGTAGAACGAAGCGGCGTCGAAGCCTGTAAAGCGGCGGTGCACAATCTTCTCGCGCAACGCGTC<br>AGTGGGCTGATCATTAACTATCCGCTGGATGACCAGGATGCCATTGCTGTGGAAGCTGCCTGCACTAATGTTCCGGCGTTATTTCTTGATGTCTCTGACCAGACACCCATCAACAGTATTATTTA<br>CTCCCATGAGGACGGTACGCGACTGGGCGTGGAGCATCTGGTCGCATTGGGTCACCAGCAAATCGCGCTGTTAGCGGGCCCATTAAGTTCTGTCTCGGCGCGTCTGCGTCTGGCTGGCTGGCA<br>TAAATATCTCACTCGCAATCAAATTCAGCCGATAGCGGAACGGGAAGGCGACTGGAGTGCCATGTCCGGTTTTCAACAAACCATGCAAATGCTGAATGAGGGCATCGTTCCCACTGCGATGCT<br>GGTTGCCAACGATCAGATGGCGCTGGGCGCAATGCGCGCCATTACCGAGTCCGGGCTGCGCGTTGGTGCGGATATCTCGGTAGTGGGATACGACGATACCGAAGATAGCTCATGTTATATCCC<br>GCCGTTAACCACCATCAAACAGGATTTTCGCCTGCTGGGGCAAACCAGCGTGGACCGCTTGCTGCAACTCTCTCAGGGCCAGGCGGTGAAGGGCAATCAGCTGTTGCCAGTCTCACTGGTGAA<br>AAGAAAAACCACCCTGGCGCCCAATACGCAAACCGCCTCTCCCCGCGCGTTGGCCGATTCATTAATGCAGCTGGCACGACAGGTTTCCCGACTGGAAAGCGGGCAGTGAGGTCTAACGCATGA<br>GAAAGCCCCCGGAAGATCACCTTCCGGGGGCTTTTTTATTGCGCCCTTGAGAGCTT***TAGCGGCCGCTGCAGGCATGCCTCGAGATGCATGGCGCCT***AACCTAAACTGACAGGCATCAAATTAA<br>GCAGAAGGCCATCCTGACGGATGGCCTTTTTGCGTTTCGAACAATTGAAAAAACCTCGCGCCTTACCTGTTGAGTAATAGTCAAAAGCCTCCGGTCGGAGGCTTTTGACTTTCTGCTTACTGAAT<br>TTCGGTGGTGCCGTTAATTAACCGGTGGGCCCTCATGATAATAATGGTTTCTTAGACGTCCGAAGTTCCTATTCTCTAGAAAGTATAGGAACTTCCATATGCCATGGAATGACCGACCGATACAA<br>TGATGATCGTGGTAAAGACTGGATTATCGTAGATATCAAAACAGCATAGAAAATAGCCAGTTTTGGGGAAGGGTTTTGGGGAAAGTTTTGGGGAAGATTTTACATCATCATAAAACAACGGGC<br>GTGTTATACGCCCGTTATAATATTTAACACATGTGGTAATTACATATTTTTGATGATCGCGTCACCAAACTCGCTGCATTTCAGCAGTTTAGCGCCTTCCATCAGACGTTCGAAGTCATAGGTTAC<br>GGTCTTGGCGTTGATTGCGCCTTCCATACCTTTAACAATCAGGTCTGCGGCTTCAGTCCAGCCCATATGGCGCAGCATCATCTCAGCGGAGAGAATAATAGAGCCAGGATTTACTTTGTCCTGAC<br>CGGCATATTTCGGCGCAGTACCGTGGGTGGCTTCAAACAGGGCGCATTCGTCACCGATGTTTGCACCAGGGGCGATACCGATACCGCCAACCTGCGCTGCCAGGGCGTCAGAAATGTAGTCAC<br>CGTTCAGGTTCATACAGGCGAT |

| | |
|---|---|
| HK022 | GGCATCAACAGCACATTCAGTGGTTTACCGTGCGCCCATTGCACCAGATATCGCCCCGGTTGATAGCCCATCTGAAACCAGGGCACACCAACGCGGCTTTTCACCTGGGGAGCATCAATAGCAT<br>TTACCAGTTCGATCACCGGCAGACTTGCTACCTGCTTTTGCAGGTCGGGAAATGAGGTCGTGCTACTACCGAGCAAAATGGCCTCTGCGCCCCACTGTTTACACTGGTCGATTTGTGCTTGCTGG<br>GTAGCCAACTGGCTGTAGCCGCCTGCCTCCAGCACTTTTAAATCCACACCGTAGCGGCGAGCAGCCTCCTGCATACCATAGTTCAACGATAACCAATATGAATCTTTCAGGCTGGGATAAAGCG<br>CGCACAGTTTCCATGCGCGTTTGGCTTTAAGCGGCGTAGAGGCTTGCACCGTGAAATGCTGCGCATCATGCCAGCGCAACAGGTTATCAGCCGAAAATGCCGGCAACATGAAAAGGGAAAGA<br>AGTAAAAATAGCAGTACGCGCATGATAGCCTCATCAATAATAAGGCTTTATGCTAGATGCATTCCGCTTTGCGACTCAACCTTTTTCACCTAAAGGATGACAAAATAACATTAATCACTTAAAAA<br>TCATCGCATTACACTAATCTGTGGTTAAATGATAGACTACATAATGCGACAAAACGCAACATATCCAGTCACTATGAATCAACTACTTAGATAGTATTAGTGACCTGAGACAGAGCATTAGCTAG<br>CTTCTTCGTCTGTTTCTACTGGTATTGGCACAAACCTGATTCCAATTTGAGCAAGGCTATGTGCCATCTCGATACTCGTTCTTAACTCAACAGAAGATGCTTTGTGCATACAGCCCCTCGTTTATTA<br>TTTATCTCCTCAGCCAGCCGCTGTGCTTTCAGTGGATTTCGGATAACATAAAGGCCGGGAAATACCCAGCCTCGCTTTGTAACGGAGTAGAGACGAAAGTGATTGCGCCTACCCGGATATTATC<br>GTGAGGATGCGTCATCGCCATTAATTCACTGATCAGTGATAAGCTGTCAAACATGAGAATTGATCCGGCTGCCTCGCGCGTTTCGGTGATGACGGTGAAAACCTCTGACACATGCAGCTCCCGG<br>AGACGGTCACAGCTTGTCTGTAAGCGGATGCCGGGAGCAGACAAGCCCGTCAGGGCGCGTCAGCGGGTGTTGGCGGGTGTCGGGGCGCAGCCATGACCCAGTCACGTAGCGATAGCGGAGT<br>GTATGCTGCACATGACATTAACCTATAAAAATAGGCGTATCACGAGGCCCTTTCGTCTTCAAGAATTAATTCCCAATTCCCCAGGCATCAAATAAAACGAAAGGCTCAGTCGAAAGACTGGGCC<br>TTTCGTTTTATCTGTTGTTTGTCGGTGAACGCTCTCCTGAGTAGGACAAATCCGCCGGGAGCGGATTTGAACGTTGCGAAGCAACGGCCCGGAGGGTGGCGGGCAGGACGCCCGCCATAAACT<br>GCCAGGAATTAATTCCCCAGGCATCAAATAAAACGAAAGGCTCAGTCGAAAGACTGGGCCTTTCGTTTTATCTGTTGTTTGTCGGTGAACGCTCTCCTGAGTAGGACAAATCCGCCGGGAGCGG<br>ATTTGAACGTTGCGAAGCAACGGCCCGGAGGGTGGCGGGCAGGACGCCCGCCATAAACTGCCAGGAATTAATTCCCCAGGCATCAAATAAAACGAAAGGCTCAGTCGAAAGACTGGGCCTTT<br>CGTTTTATCTGTTGTTTGTCGGTGAACGCTCTCCTGAGTAGGACAAATCCGCCGGGAGCGGATTTGAACGTTGCGAAGCAACGGCCCGGAGGGTGGCGGGCAGGACGCCCGCCATAAACT***GC***<br>***CAGGAATTGGGGATCGGAATTCGAGCTCGGTACCCGGG***GGAGAACGATCGTTGGCTGTGTTGACAATTAATCATCGGCTCGTATAATGTGTGGAATTGTGAGCGCTCACAATTTACTGAAAG<br>ATGCGATTTACTAAATGAACACGATTAACATCGCTAAGAACGACTTCTCTGACATCGAACTGGCTGCTATCCCGTTCAACACTCTGGCTGACCATTACGGTGAGCGTTTAGCTCGCGAACAGTTG<br>GCCCTTGAGCATGAGTCTTACGAGATGGGTGAAGCACGCTTCCGCAAGATGTTTGAGCGTCAACTTAAAGCTGGTGAGGTTGCGGATAACGCTGCCGCCAAGCCTCTCATCACTACCCTACTCC<br>CTAAGATGATTGCACGCATCAACGACTGGTTTGAGGAAGTGAAAGCTAAGCGCGGCAAGCGCCCGACAGCCTTCCAGTTCCTGCAAGAAATCAAGCCGGAAGCCGTAGCGTACATCACCATTA<br>AGACCACTCTGGCTTGCCTAACCAGTGCTGACAATACAACCGTTCAGGCTGTAGCAAGCGCAATCGGTCGGGCCATTGAGGACGAGGCTCGCTTCGGTCGTATCCGTGACCTTGAAGCTAAGC<br>ACTTCAAGAAAAACGTTGAGGAACAACTCAACAAGCGCGTAGGGCACGTCTACAAGGGCTTGTCGACGACGGCGTGCTCAGTGGTGTACGGTACAAACCAGGGTTCTAAAGCATTTATGCAA<br>GTTGTCGAGGCTGACATGCTCTCTAAGGGTCTACTCGGTGGCGAGGCGTGGTCTTCGTGGCATAAGGAAGACTCTATTCATGTAGGAGTACGCTGCATCGAGATGCTCATTGAGTCAACCGGA<br>ATGGTTAGCTTACACCGCCAAAATGCTGGCGTAGTAGGTCAAGACTCTGAGACTATCGAACTCGCACCTGAATACGCTGAGGCTATCGCAACCCGTGCAGGTGCGCTGGCTGGCATCTCTCCGA<br>TGTTCCAACCTTGCGTAGTTCCTCCTAAGCCGTGGACTGGCATTACTGGTGGTGGCTATTGGGCTAACGGTCGTCGTCCTCTGGCGCTGGTGCGTACTCACAGTAAGAAAGCACTGATGCGCTA<br>CGAAGACGTTTACATGCCTGAGGTGTACAAAGCGATTAACATTGCGCAAAACACCGCATGGAAAATCAACAAGAAAGTCCTAGCGGTCGCCAACGTAATCACCAAGTGGAAGCATTGTCCGGT<br>CGAGGACATCCCTGCGATTGAGCGTGAAGAACTCCCGATGAAACCGGAAGACATCGACATGAATCCTGAGGCTCTCACCGCGTGGAAACGTGCTGCCGCTGCTGTGTACCGCAAGGACAAGG<br>CTCGCAAGTCTCGCCGTATCAGCCTTGAGTTCATGCTTGAGCAAGCCAATAAGTTTGCTAACCATAAGGCCATCTGGTTCCCTTACAACATGGACTGGCGCGGTCGTGTTTACGCTGTGTCAATG<br>TTCAACCCGCAAGGTAACGATATGACCAAAGGACTGCTTACGCTGGCGAAAGGTAAACCAATCGGTAAGGAAGGTTACTACTGGCTGAAAATCCACGGTGCAAACTGTGCGGGTGTCGATAA<br>GGTTCCGTTCCCTGAGCGCATCAAGTTCATTGAGGAAAACCACGAGAACATCATGGCTTGCGCTAAGTCTCCACTGGAGAACACTTGGTGGGCTGAGCAAGATTCTCCGTTCTGCTTCCTTGCG<br>TTCTGCTTTGAGTACGCTGGGGTACAGCACCACGGCCTGAGCTATAACTGCTCCCTTCCGCTGGCGTTTGACGGGTCTTGCTCTGGCATCCAGCACTTCTCCGCGATGCTCCGAGATGAGGTAG<br>GTGGTCGCGCGGTTAACTTGCTTCCTAGTGAAACCGTTCAGGACATCTACGGGATTGTTGCTAAGAAAGTCAACGAGATTCTACAAGCAGACGCAATCAATGGGACCGATAACGAAGTAGTTA<br>CCGTGACCGATGAGAACACTGGTGAAATCTCTGAGAAAGTCAAGCTGGGCACTAAGGCACTGGCTGGTCAATGGCTGGCTTACGGTGTTACTCGCAGTGTGACTAAGCGTTCAGTCATGACGC<br>TGGCTTACGGGTCCAAAGAGTTCGGCTTCCGTCAACAAGTGCTGGAAGATACCATTCAGCCAGCTATTGATTCCGGCAAGGGTCTGATGTTCACTCAGCCGAATCAGGCTGCTGGATACATGGC<br>TAAGCTGATTTGGGAATCTGTGAGCGTGACGGTGGTAGCTGCGGTTGAAGCAATGAACTGGCTTAAGTCTGCTGCTAAGCTGCTGGCTGCTGAGGTCAAAGATAAGAAGACTGGAGAGATTC<br>TTCGCAAGCGTTGCGCTGTGCATTGGGTAACTCCTGATGGTTTCCCTGTGTGGCAGGAATACAAGAAGCCTATTCAGACGCGCTTGAACCTGATGTTCCTCGGTCAGTTCCGCTTACAGCCTACC<br>ATTAACACCAACAAAGATAGCGAGATTGATGCACACAAACAGGAGTCTGGTATCGCTCCTAACTTTGTACACAGCCAAGACGGTAGCCACCTTCGTAAGACTGTAGTGTGGGCACACGAGAAG<br>TACGGAATCGAATCTTTTGCACTGATTCACGACTCCTTCGGTACCATTCCGGCTGACGCTGCGAACCTGTTCAAAGCAGTGCGCGAAACTATGGTTGACACATATGAGTCTTGTGATGTACTGGC<br>TGATTTCTACGACCAGTTCGCTGACCAGTTGCACGAGTCTCAATTGGACAAAATGCCAGCACTTCCGGCTAAAGGTAACTTGAACCTCCGTGACATCTTAGAGTCGGACTTCGCGTTCGCGTAAT<br>AAAGGTTTCAGCCAAAAAACTTAAGACCGCCGGTCTTGTCCACTACCTTGCAGTAATGCGGTGGACAGGATCGGCGGTTTTCTTTTCTCTTCTCAAGCTT***GCACTGAAGGTCCTCAATCGCACTG***<br>***GAAACATCAAGGTCG***GGAGGACACCATCGAATGGCGCAAAACCTTTCGCGGTATGGCATGATAGCGCCCGGAAGAGAGTCAATTCAGGGTGGTGAATAATGAAACCAGTAACGTTATACGAT<br>GTCGCAGAGTATGCCGGTGTCTCTTATATGACCGTTTCCCGCGTGGTGAACCAGGCCAGCCACGTTTCTGCGAAAACGCGGGAAAAAGTGGAAGCGGCGATGGTGGAGCTGAATTACATTCCC<br>AACCGCGTGGCACAACAACTGGCGGGCAAACAGTCGTTGCTGATTGGCGTTGCCACCTCCAGTCTGGCCCTGCACGCGCCGTCGCAAATTGTCGCGGCGATTAAATCTCGCGCCGATCAACTG<br>GGTGCCAGCGTGGTGGTGTCGATGGTAGAACGAAGCGGCGTCGAAGCCTGTAAAGCGGCGGTGCACAATCTTCTCGCGCAACGCGTCAGTGGGCTGATCATTAACTATCCGCTGGATGACCA<br>GGATGCCATTGCTGTGGAAGCTGCCTGCACTAATGTTCCGGCGTTATTTCTTGATGTCTCTGACCAGACACCCATCAACAGTATTATTTACTCCCATGAGGACGGTACGCGACTGGGCGTGGAG<br>CATCTGGTCGCATTGGGTCACCAGCAAATCGCGCTGTTAGCGGGCCCATTAAGTTCTGTCTCGGCGCGTCTGCGTCTGGCTGGCTGGCATAAATATCTCACTCGCAATCAAATTCAGCCGATAGC<br>GGAACGGGAAGGCGACTGGAGTGCCATGTCCGGTTTTCAACAAACCATGCAAATGCTGAATGAGGGCATCGTTCCCACTGCGATGCTGGTTGCCAACGATCAGATGGCGCTGGGCGCAATGC<br>GCGCCATTACCGAGTCCGGGCTGCGCGTTGGTGCGGATATCTCGGTAGTGGGATACGACGATACCGAAGATAGCTCATGTTATATCCCGCCGTTAACCACCATCAAACAGGATTTTCGCCTGCT<br>GGGGCAAACCAGCGTGGACCGCTTGCTGCAACTCTCTCAGGGCCAGGCGGTGAAGGGCAATCAGCTGTTGCCAGTCTCACTGGTGAAAAGAAAAACCACCCTGGCGCCCAATACGCAAACCG<br>CCTCTCCCCGCGCGTTGGCCGATTCATTAATGCAGCTGGCACGACAGGTTTCCCGACTGGAAAGCGGGCAGTGAGGTCTAACGCATGAGAAAGCCCCCGGAAGATCACCTTCCGGGGGCTTTT<br>TTATTGCGCCCTTGAGAGCTT***TAGCGGCCGCTGCAGGCATGCCTCGAGATGCATGGCGCCT***AACCTAAACTGACAGGCATCAAATTAAGCAGAAGGCCATCCTGACGGATGGCCTTTTTGCGTT<br>TCGAACAATTGAAAAAACCTCGCGCCTTACCTGTTGAGTAATAGTCAAAAGCCTCCGGTCGGAGGCTTTTGACTTTCTGCTTACTGAATTTCGGTGGTGCCGTTAATTAACCGGTGGGCCCTCAT<br>GATAATAATGGTTTCTTAGACGTCCGAAGTTCCTATTCTCTAGAAAGTATAGGAACTTCCATATGCCATGGAATGACCGACCGATACAATGATGATCGTGGTAAAGACTGGATTATCGTAGATAT<br>CAAAACAGCATAGAAAATAGCCAGTTTTGGGGAAGGGTTTTGGGGAAAGTTTTGGGGAAGATTTTACATCATCATAAAACAACGGGCGTGTTATACGCCCGTTATAATATTTAACACATGTGGT<br>AATTACATATTTTTGATGATCGCGTCACCAAACTCGCTGCATTTCAGCAGTTTAGCGCCTTCCATCAGACGTTCGAAGTCATAGGTTACGGTCTTGGCGTTGATTGCGCCTTCCATACCTTTAACA<br>ATCAGGTCTGCGGCTTCAGTCCAGCCCATATGGCGCAGCATCATCTCAGCGGAGAGAATAATAGAGCCAGGATTTACTTTGTCCTGACCGGCATATTTCGGCGCAGTACCGTGGGTGGCTTCAA<br>ACAGGGCGCATTCGTCACCGATGTTTGCACCAGGGGCGATACCGATACCGCCAACCTGCGCTGCCAGGGCGTCAGAAATGTAGTCACCGTTCAGGTTCATACAGGCGAT |

Table S4. Naive sX strain with IPTG inducible T7 RNAP. Promoters, RBSs, coding sequences, inserted bases in T7 RNAP CDS, terminators, integrase attachment sites, ***Gibson homologies***, genomic pOSIP primer binding sites. Gap in expected UNS sequence for Gibson indicated by “_”.

### eRWdiff1x

| Cluster | Sequence |
|---|---|
| P21<br>(T) | TAGAACTACCACCTGACCGGCCAGCCTACCCCAAAACTACCGAGGGGTTTTGATAACTTTTTTTGATAAAATTTTGATAACCGTTAGCAGCCTAACAAAAAACAACGGGAGCGTTACGCTCCCGT<br>TAATAAATTTAACAAACTACGGCATTACATGTTTTCGATGATCGCGTCACCAAACTCTGAACATTTCAGCAGTTTAGCGCCATCCATCAGACGCTCGAAGTCATAGGTTACGGTTTTCGCGTTGAT<br>TGCGCCTTCCATACCTTTAACAATTAAGTCAGCCGCTTCGGTCCAACCCATGTGGCGCAGCAGTATATAAAAATTATGTATTCAACTCGCTGATATTTAATCATAAATATTATTTTCTTGTCTTTGT<br>ATTAGCCAAAATCAACTTTATTCAACCGTTTGATTTCATTGACGTTCGTTTTTATTTTGGGGAAAGATTTTCTCTACCATTTATATCAAAATTACAAGTCTACATTTTATTAACATCTCAATACATGT<br>GTGGCCTTGCAAGCCATTGCGAGGCCTTATGCTAGCTTCTTCGTCTGTTTCTACTGGTATTGGCACAAACCTGATTCCAATTTGAGCAAGGCTATGTGCCATCTCGATACTCGTTCTTAACTCAAC<br>AGAAGATGCTTTGTGCATACAGCCCCTCGTTTATTATTTATCTCCTCAGCCAGCCGCTGTGCTTTCAGTGGATTTCGGATAACAGAAAGGCCGGGAAATACCCAGCCTCGCTTTGTAACGGAGTA<br>GAGACGAAAGTGATTGCGCCTACCCGGATATTATCGTGAGGATGCGTCATCGCCATTAATTCACTGATCAGTGATAAGCTGTCAAACATGAGAATTGATCCGGCTGCCTCGCGCGTTTCGGTGA<br>TGACGGTGAAAACCTCTGACACATGCAGCTCCCGGAGACGGTCACAGCTTGTCTGTAAGCGGATGCCGGGAGCAGACAAGCCCGTCAGGGCGCGTCAGCGGGTGTTGGCGGGTGTCGGGGC<br>GCAGCCATGACCCAGTCACGTAGCGATAGCGGAGTGTATGCTGCACATGACATTAACCTATAAAAATAGGCGTATCACGAGGCCCTTTCGTCTTCAAGAATTAATTCCCAATTCCCCAGGCATCA<br>AATAAAACGAAAGGCTCAGTCGAAAGACTGGGCCTTTCGTTTTATCTGTTGTTTGTCGGTGAACGCTCTCCTGAGTAGGACAAATCCGCCGGGAGCGGATTTGAACGTTGCGAAGCAACGGCC<br>CGGAGGGTGGCGGGCAGGACGCCCGCCATAAACTGCCAGGAATTAATTCCCCAGGCATCAAATAAAACGAAAGGCTCAGTCGAAAGACTGGGCCTTTCGTTTTATCTGTTGTTTGTCGGTGAA<br>CGCTCTCCTGAGTAGGACAAATCCGCCGGGAGCGGATTTGAACGTTGCGAAGCAACGGCCCGGAGGGTGGCGGGCAGGACGCCCGCCATAAACTGCCAGGAATTAATTCCCCAGGCATCAAA<br>TAAAACGAAAGGCTCAGTCGAAAGACTGGGCCTTTCGTTTTATCTGTTGTTTGTCGGTGAACGCTCTCCTGAGTAGGACAAATCCGCCGGGAGCGGATTTGAACGTTGCGAAGCAACGGCCCG<br>GAGGGTGGCGGGCAGGACGCCCGCCATAAACT***GCCAGGAATTGGGGATCGGAATTCGAGCTCGGTACCCGGG***CTCGCATCCATTCTCAGGCTGTCTCGTCTCGTCTCGGAGAACGATCGTTG<br>GCTGTGTTGACAATTAATCATCGGCTCGTATAATGTGTGGAATTGTGAGCGCTCACAATTTACTGAAAGATGCGATTTACTAAATGAACACGATTAACATCGCTAAGAACGACTTCTCTGACATC<br>GAACTGGCTGCTATCCCGTTCAACACTCTGGCTGACCATTACGGTGAGCGTTTAGCTCGCGAACAGTTGGCCCTTGAGCATGAGTCTTACGAGATGGGTGAAGCACGCTTCCGCAAGATGTTTG<br>AGCGTCAACTTAAAGCTGGTGAGGTTGCGGATAACGCTGCCGCCAAGCCTCTCATCACTACCCTACTCCCTAAGATGATTGCACGCATCAACGACTGGTTTGAGGAAGTGAAAGCTAAGCGCG<br>GCAAGCGCCCGACAGCCTTCCAGTTCCTGCAAGAAATCAAGCCGGAAGCCGTAGCGTACATCACCATTAAGACCACTCTGGCTTGCCTAACCAGTGCTGACAATACAACCGTTCAGGCTGTAGC<br>AAGCGCAATCGGTCGGGCCATTGAGGACGAGGCTCGCTTCGGTCGTATCCGTGACCTTGAAGCTAAGCACTTCAAGAAAAACGTTGAGGAACAACTCAACAAGCGCGTAGGGCACGTCTACA<br>AGGGCTTGTCGACGACGGCGTGCTCCGTCGTCAGGATCATAGGTGACGAACAATAAGGCCTCCCTAACGGGGGGCCTTTTTTATTGATAACAAAAGCTT***GCACTGAAGGTCCTCAATCGCACT***<br>***GGAAACATCAAGGTCG***AAGCTCTCAAGGGCGCAATAAAAAAGCCCCCGGAAGGTGATCTTCCGGGGGCTTTCTCATGCGTTAGACCTTCACCCCTTAGCTTTTTGGGAGGTACGGTTTCATCA<br>AATACCTTGAGAAAAGCTGCATCAGCTTCAGATAAATTCATAAAAAAAGCTTCATCATCTTTATCGCCAGAAAATTCATCTTCATCAACGACAAATTCGAACTTCAGCTTACTAATTTTTCTTCCTT<br>CTTTTTCATGAACAGTGAAGCCAACAAACGATATTTCTGTTTTCTTTTTAATTTCAGCAATGGCTTTATTTAACACATCCCTTTTAAAAATAGGAAAGTCAGGGTATTTGTACTCAATATTTCCATCT<br>TTATCAAAAGTATAAGCTATTAACTCTTCCTTTAACTCATCAACGGAAATAATAAAATAATTTTTCTTCTTAAAATTAGAGTAATGCTTCCTGATAAGTTGATAAAGAGAAGATGAATACTGGCTA<br>CTTAAGCGTAAGCTTGCCGTTAACAATTGCGTTGTGAATTTATTTTTTTTCCCAATAAGGCTAGAGATATATGGTTCTATGGTTCTGGTGAATTTTAAAGACAAGTATCCTTCATCATTTGAATAAG<br>CTATCCACTCAATAATGTTAAGATCTAACTCTTCAGGGGAGTTTTTAGCAGTAAAGGGCAGGTTAAGCTCTTTAGCTAAAGCAATGATATCATCCCCTCTTAGCGAAATTTTGCTGGCACCAAGT<br>AATTTACCACCCTCTTTTAATTGTCGATAAGCAAGCGATGGGGTGATTTTGGCGAGCGCTGCAAGATCTTCAGCCCTAATTTTGAAAACTCGCCCTCGTTCAAGAGGTTCTTTGCTATCAATGGG<br>AGCAAGCGCCATATACATCACTCGCTTTGCGGGCAAAGGAAGTTGAGCCAGGGTGTGATTTAGCTCGTTTCGGTGGCGAATTTTCGTTTTTTTGTTCACGTCCATCATGACCTTGAGTCTCATTT<br>AGTATTTCTCCTCTTTCTCTAGTATTCTTCGAGCCTAGCAAGGGTCCGGGTTCACCGAAATCTATCTCATTTGCTAGTTCAGCCAACGATCGTTCTCC***CTGACCTCCTGCCAGCAATAGTAAGACA***<br>***ACACGCAAAGTC***GGAGGGAAACACAGAAAAAAGCCCGCACCTGACAGTGCGGGCTTTTTTTTTCGACCAAAGGTACTGGTTTGTCTGGTCAACCACCGCGTGCTCAGTGGTGTACGGTACAAA<br>CCAGGGTTCTAAAGCATTTATGCAAGTTGTCGAGGCTGACATGCTCTCTAAGGGTCTACTCGGTGGCGAGGCGTGGTCTTCGTGGCATAAGGAAGACTCTATTCATGTAGGAGTACGCTGCATC<br>GAGATGCTCATTGAGTCAACCGGAATGGTTAGCTTACACCGCCAAAATGCTGGCGTAGTAGGTCAAGACTCTGAGACTATCGAACTCGCACCTGAATACGCTGAGGCTATCGCAACCCGTGCA<br>GGTGCGCTGGCTGGCATCTCTCCGATGTTCCAACCTTGCGTAGTTCCTCCTAAGCCGTGGACTGGCATTACTGGTGGTGGCTATTGGGCTAACGGTCGTCGTCCTCTGGCGCTGGTGCGTACTC<br>ACAGTAAGAAAGCACTGATGCGCTACGAAGACGTTTACATGCCTGAGGTGTACAAAGCGATTAACATTGCGCAAAACACCGCATGGAAAATCAACAAGAAAGTCCTAGCGGTCGCCAACGTAA<br>TCACCAAGTGGAAGCATTGTCCGGTCGAGGACATCCCTGCGATTGAGCGTGAAGAACTCCCGATGAAACCGGAAGACATCGACATGAATCCTGAGGCTCTCACCGCGTGGAAACGTGCTGCCG<br>CTGCTGTGTACCGCAAGGACAAGGCTCGCAAGTCTCGCCGTATCAGCCTTGAGTTCATGCTTGAGCAAGCCAATAAGTTTGCTAACCATAAGGCCATCTGGTTCCCTTACAACATGGACTGGCG<br>CGGTCGTGTTTACGCTGTGTCAATGTTCAACCCGCAAGGTAACGATATGACCAAAGGACTGCTTACGCTGGCGAAAGGTAAACCAATCGGTAAGGAAGGTTACTACTGGCTGAAAATCCACGG<br>TGCAAACTGTGCGGGTGTCGATAAGGTTCCGTTCCCTGAGCGCATCAAGTTCATTGAGGAAAACCACGAGAACATCATGGCTTGCGCTAAGTCTCCACTGGAGAACACTTGGTGGGCTGAGCA<br>AGATTCTCCGTTCTGCTTCCTTGCGTTCTGCTTTGAGTACGCTGGGGTACAGCACCACGGCCTGAGCTATAACTGCTCCCTTCCGCTGGCGTTTGACGGGTCTTGCTCTGGCATCCAGCACTTCTC<br>CGCGATGCTCCGAGATGAGGTAGGTGGTCGCGCGGTTAACTTGCTTCCTAGTGAAACCGTTCAGGACATCTACGGGATTGTTGCTAAGAAAGTCAACGAGATTCTACAAGCAGACGCAATCAA<br>TGGGACCGATAACGAAGTAGTTACCGTGACCGATGAGAACACTGGTGAAATCTCTGAGAAAGTCAAGCTGGGCACTAAGGCACTGGCTGGTCAATGGCTGGCTTACGGTGTTACTCGCAGTG<br>TGACTAAGCGTTCAGTCATGACGCTGGCTTACGGGTCCAAAGAGTTCGGCTTCCGTCAACAAGTGCTGGAAGATACCATTCAGCCAGCTATTGATTCCGGCAAGGGTCTGATGTTCACTCAGCC<br>GAATCAGGCTGCTGGATACATGGCTAAGCTGATTTGGGAATCTGTGAGCGTGACGGTGGTAGCTGCGGTTGAAGCAATGAACTGGCTTAAGTCTGCTGCTAAGCTGCTGGCTGCTGAGGTCAA<br>AGATAAGAAGACTGGAGAGATTCTTCGCAAGCGTTGCGCTGTGCATTGGGTAACTCCTGATGGTTTCCCTGTGTGGCAGGAATACAAGAAGCCTATTCAGACGCGCTTGAACCTGATGTTCCTC<br>GGTCAGTTCCGCTTACAGCCTACCATTAACACCAACAAAGATAGCGAGATTGATGCACACAAACAGGAGTCTGGTATCGCTCCTAACTTTGTACACAGCCAAGACGGTAGCCACCTTCGTAAGA<br>CTGTAGTGTGGGCACACGAGAAGTACGGAATCGAATCTTTTGCACTGATTCACGACTCCTTCGGTACCATTCCGGCTGACGCTGCGAACCTGTTCAAAGCAGTGCGCGAAACTATGGTTGACAC<br>ATATGAGTCTTGTGATGTACTGGCTGATTTCTACGACCAGTTCGCTGACCAGTTGCACGAGTCTCAATTGGACAAAATGCCAGCACTTCCGGCTAAAGGTAACTTGAACCTCCGTGACATCTTAG<br>AGTCGGACTTCGCGTTCGCGTAATAAAGGTTTCAGCCAAAAAACTTAAGACCGCCGGTCTTGTCCACTACCTTGCAGTAATGCGGTGGACAGGATCGGCGGTTTTCTTTTCTCTTCTCAAGCTT***G***<br>***AGCCAACTCCCTTTACAACCTCACTCAAGTCCGTTAGAG***GGAGTTTACGGCTAGCTCAGTCCTAGGTATAGTGCTAGCTACTAGAGAAAGAGGAGAAATACTAAATGGAACTGCGTGACCTGG<br>ATTTAAACCTGCTGGTGGTGTTCAACCAGTTGCTGGTCGACAGACGCGTCTCTGTCACTGCGGAGAACCTGGGCCTGACCCAGCCTGCCGTGAGCAATGCGCTGAAACGCCTGCGCACCTCGCT<br>ACAGGACCCACTCTTCGTGCGCACACATCAGGGAATGGAACCCACACCCTATGCCGCGCATCTGGCCGAGCACGTCACTTCGGCCATGCACGCACTGCGCAACGCCCTACAGCACCATGAAAG<br>CTTCGATCCGCTGACCAGCGAGCGTACCTTCACCCTGGCCATGACCGACATTGGCGAGATCTACTTCATGCCGCGGCTGATGGATGCGCTGGCTCACCAGGCCCCCAATTGCGTGATCAGTACG<br>GTGCGCGACAGTTCGATGAGCCTGATGCAGGCCTTGCAGAACGGAACCGTGGACTTGGCCGTGGGCCTGCTTCCCAATCTGCAAACTGGCTTCTTTCAGCGCCGGCTGCTCCAGAATCACTACG<br>TGTGCCTATGTCGCAAGGACCATCCAGTCACCCGCGAACCCCTGACTCTGGAGCGCTTCTGTTCCTACGGCCACGTGCGTGTCATCGCCGCTGGCACCGGCCACGGCGAGGTGGACACGTACAT<br>GACACGGGTCGGCATCCGGCGCGACATCCGTCTGGAAGTGCCGCACTTCGCCGCCGTTGGCCACATCCTCCAGCGCACCGATCTGCTCGCCACTGTGCCGATATGTTTAGCCGACTGCTGCGTA<br>GAGCCCTTCGGCCTAAGCGCCTTGCCGCACCCAGTCGTCTTGCCTGAAATAGCCATCAACATGTTCTGGCATGCGAAGTACCACAAGGACCTAGCCAATATTTGGTTGCGGCAACTGATGTTTG<br>ACCTGTTTACGGATTGATACTGGGCCCAAGTTCACTTAAAAAGGAGATCAACAATGAAAGCAATTTTCGTACTGAAACATCTTAATCATGCTAAGGAGGTTTTCTAATGGCCTTGGTTGACGGTT<br>TTCTTGAGCTGGAACGCTCAAGTGGAAAATTGGAGTTGAGCGCCATCCTGCAGAAGATGGCGAGCGACCTCGGATTCTCGAAGATCCTGTTCGGCCTGTTGCCTAAGGACAGCCAGGACTACG<br>AGAACGCCTTCATCGTCAGCAACTACCCGGCCGCCTGGCGCGAGCATTACGACCGAACTGGCTACGCGCGGGTCGACCCGACGGTCAGTCACTGTACCCAGAGCGTACTGCCGATTTTCTGGG<br>AACCGCCCATCTACCAGACGCGAAAGCAGCACGAGTTCTTCGAGGAAGCCTTGGCCGCTGGCCTGGTGTATGGACTGACCATGCCGCTGCATGGTGCTCGCGGCGAACTCGGCGCGCTGAGC<br>CTCAGCGTGGAAGCGGAAAACCGGGCCGAGGCCAACCGTTTCATGGAGTCGGTCCTGCCGACCCTGTGGATGCTCAAGGACTACGCACTGCAGAGCGGTGTCGGAATGGCCTTCGAACATCC<br>GATCAGCAAACCGGTGGTTCTGACCAGCCGAGAGAAGGAAGTGTTACAGTGGTGCGCCATCGGCAAGACCAGCTGGGAGATATCGGTTATCTGCAACTGCTCGGAAGCCAATGTGAACTTCC<br>ATATGGGAAATATTCGGCGGAAGTTCGGTGTGACCTCCCGCCGCGTAGCGGCCATTATGGCCGTTAATTTGGGTCTTATTACTCTCTGAGGTCCAATTATTGAACACCCTTCGGGGTGTTTTTTT<br>GTTTCTGGTCTGCCGCTT***CTCGTTCGCTGCCACCTAAGAATACTCTACGGTCACATAC***GGAGGACACCATCGAATGGCGCAAAACCTTTCGCGGTATGGCATGATAGCGCCCGGAAGAGAGTC<br>AATTCAGGGTGGTGAATAATGAAACCAGTAACGTTATACGATGTCGCAGAGTATGCCGGTGTCTCTTATATGACCGTTTCCCGCGTGGTGAACCAGGCCAGCCACGTTTCTGCGAAAACGCGG<br>GAAAAAGTGGAAGCGGCGATGGTGGAGCTGAATTACATTCCCAACCGCGTGGCACAACAACTGGCGGGCAAACAGTCGTTGCTGATTGGCGTTGCCACCTCCAGTCTGGCCCTGCACGCGCC<br>GTCGCAAATTGTCGCGGCGATTAAATCTCGCGCCGATCAACTGGGTGCCAGCGTGGTGGTGTCGATGGTAGAACGAAGCGGCGTCGAAGCCTGTAAAGCGGCGGTGCACAATCTTCTCGCGC<br>AACGCGTCAGTGGGCTGATCATTAACTATCCGCTGGATGACCAGGATGCCATTGCTGTGGAAGCTGCCTGCACTAATGTTCCGGCGTTATTTCTTGATGTCTCTGACCAGACACCCATCAACAGT<br>ATTATTTACTCCCATGAGGACGGTACGCGACTGGGCGTGGAGCATCTGGTCGCATTGGGTCACCAGCAAATCGCGCTGTTAGCGGGCCCATTAAGTTCTGTCTCGGCGCGTCTGCGTCTGGCTG<br>GCTGGCATAAATATCTCACTCGCAATCAAATTCAGCCGATAGCGGAACGGGAAGGCGACTGGAGTGCCATGTCCGGTTTTCAACAAACCATGCAAATGCTGAATGAGGGCATCGTTCCCACTG<br>CGATGCTGGTTGCCAACGATCAGATGGCGCTGGGCGCAATGCGCGCCATTACCGAGTCCGGGCTGCGCGTTGGTGCGGATATCTCGGTAGTGGGATACGACGATACCGAAGATAGCTCATGT<br>TATATCCCGCCGTTAACCACCATCAAACAGGATTTTCGCCTGCTGGGGCAAACCAGCGTGGACCGCTTGCTGCAACTCTCTCAGGGCCAGGCGGTGAAGGGCAATCAGCTGTTGCCAGTCTCAC<br>TGGTGAAAAGAAAAACCACCCTGGCGCCCAATACGCAAACCGCCTCTCCCCGCGCGTTGGCCGATTCATTAATGCAGCTGGCACGACAGGTTTCCCGACTGGAAAGCGGGCAGTGAGGTTCTC<br>GGTACCAAATTCCAGAAAAGAGACGCTTTCGAGCGTCTTTTTTCGTTTTGGTCCGCTT***TAGCGGCCGCTGCAGGCATGCCTCGAGATGCATGGCGCCT***AACCTAAACTGACAGGCATCAAATTA<br>AGCAGAAGGCCATCCTGACGGATGGCCTTTTTGCGTTTCGAACAATTGAAAAAACCTCGCGCCTTACCTGTTGAGTAATAGTCAAAAGCCTCCGGTCGGAGGCTTTTGACTTTCTGCTTACTGAA<br>TTTCGGTGGTGCCGTTAATTAACCGGTGGGCCCTCATGATAATAATGGTTTCTTAGACGTCCGAAGTTCCTATTCTCTAGAAAGTATAGGAACTTCCATATGCCATGGAATGACCGACCGATACA<br>ATGATGATCGTGGTAAAGACTGGATTATCGTAGATATCAAAACAGCATAGAAAATAGCCAGTTTTGGGGAAGGGTTTTGGGGAAAGTTTTGGGGAAGATTTTACATCATCATAAAACAACGGG<br>CGTGTTATACGCCCGTTATAATATTTAACACATGTGGTAATTACATATTTTTGATGATCGCGTCACCAAACTCGCTGCATTTCAGCAGTTTAGCGCCTTCCATCAGACGTTCGAAGTCATAGGTTA<br>CGGTCTTGGCGTTGATTGCGCCTTCCATACCTTTAACAATCAGGTCTGCGGCTTCAGTCCAGCCCATATGGCGCAGCATCATCTCAGCGGAGAGAATAATAGAGCCAGGATTTACTTTGTCCTGA<br>CCGGCATATTTCGGCGCAGTACCGTGGGTGGCTTCAAACAGGGCGCATTCGTCACCGATGTTTGCACCAGGGGCGATACCGATACCGCCAACCTGCGCTGCCAGGGCGTCAGAAATGTAGTCA<br>CCGTTCAGGTTCATACAGGCGAT |

| | |
|---|---|
| Φ186 #1 | GATCATCATGTTTATTGCGTGGTTTATAAGTATTGTTATTGCACACTCAAGGCCCTTTAGCTCAGTGGTTAGAGCAGGCGACTCATAATCGCTTGGTCGCTGGTTCAAGTCCAGCAGGGGCCACC<br>AAATTTAAGTAGTAAATACATACAGTTAGGCCACTCTCTCGAGTGGCCTTTTTGTTGCCTCGTTTTCGAGTGGCGATAGAATGGCGGTGGATTTTTTACCGCCACTTTCCTGAGGAGTGCATAGC<br>TAGCTTCTTCGTCTGTTTCTACTGGTATTGGCACAAACCTGATTCCAATTTGAGCAAGGCTATGTGCCATCTCGATACTCGTTCTTAACTCAACAGAAGATGCTTTGTGCATACAGCCCCTCGTTTA<br>TTATTTATCTCCTCAGCCAGCCGCTGTGCTTTCAGTGGATTTCGGATAACAGAAAGGCCGGGAAATACCCAGCCTCGCTTTGTAACGGAGTAGAGACGAAAGTGATTGCGCCTACCCGGATATT<br>ATCGTGAGGATGCGTCATCGCCATTAATTCACTGATCAGTGATAAGCTGTCAAACATGAGAATTGATCCGGCTGCCTCGCGCGTTTCGGTGATGACGGTGAAAACCTCTGACACATGCAGCTCC<br>CGGAGACGGTCACAGCTTGTCTGTAAGCGGATGCCGGGAGCAGACAAGCCCGTCAGGGCGCGTCAGCGGGTGTTGGCGGGTGTCGGGGCGCAGCCATGACCCAGTCACGTAGCGATAGCG<br>GAGTGTATGCTGCACATGACATTAACCTATAAAAATAGGCGTATCACGAGGCCCTTTCGTCTTCAAGAATTAATTCCCAATTCCCCAGGCATCAAATAAAACGAAAGGCTCAGTCGAAAGACTG<br>GGCCTTTCGTTTTATCTGTTGTTTGTCGGTGAACGCTCTCCTGAGTAGGACAAATCCGCCGGGAGCGGATTTGAACGTTGCGAAGCAACGGCCCGGAGGGTGGCGGGCAGGACGCCCGCCATA<br>AACTGCCAGGAATTAATTCCCCAGGCATCAAATAAAACGAAAGGCTCAGTCGAAAGACTGGGCCTTTCGTTTTATCTGTTGTTTGTCGGTGAACGCTCTCCTGAGTAGGACAAATCCGCCGGGA<br>GCGGATTTGAACGTTGCGAAGCAACGGCCCGGAGGGTGGCGGGCAGGACGCCCGCCATAAACTGCCAGGAATTAATTCCCCAGGCATCAAATAAAACGAAAGGCTCAGTCGAAAGACTGGG<br>CCTTTCGTTTTATCTGTTGTTTGTCGGTGAACGCTCTCCTGAGTAGGACAAATCCGCCGGGAGCGGATTTGAACGTTGCGAAGCAACGGCCCGGAGGGTGGCGGGCAGGACGCCCGCCATAAA<br>CT***GCCAGGAATTGGGGATCGGAATTCGAGCTCGGTACCCGGG***CTCGCATCCATTCTCAGGCTGTCTCGTCTCGTCTCGGAGAACGATCGTTGGCTGGGGGCCTCGCTTGGGTTATTGCTGGTG<br>CCCGGCCGGGCGCAATATTCATGTTGATGATTTATTATATATCGAGTGGTGTATTTATTTATATTGTTTGCTCCGTTACCGTTATTAACTACTAGAGAAAGAGGAGAAATACTAAATGGTGAGAG<br>CCCTGGTAGTCATCCGCCTGTCCCGCGTCACCGATGCTACGACTTCACCGGAGCGTCAGCTGGAGTCTTGCCAGCAGCTCTGCGCCCAGCGCGGCTGGGACGTCGTCGGGGTAGCGGAGGATC<br>TGGACGTCTCCGGGGCGGTCGATCCGTTCGACCGGAAGCGCAGACCGAACCTGGCCCGGTGGCTAGCGTTCGAGGAGCAACCGTTTGACGTGATCGTGGCGTACCGGGTAGATCGGTTGACC<br>CGATCGATCCGGCATCTTCAGCAGCTGGTCCACTGGGCCGAGGACCACAAGAAGCTGGTCGTCTCCGCGACCGAAGCGCACTTCGATACGACGACGCCGTTTGCGGCGGTCGTCATCGCGCTT<br>ATGGGAACGGTGGCGCAGATGGAATTAGAAGCGATCAAAGAGCGGAACCGTTCGGCTGCGCATTTCAATATCCGCGCCGGGAAATACCGAGGATCCCTGCCGCCGTGGGGATACCTGCCTAC<br>GCGCGTGGACGGGGAGTGGCGGCTGGTGCCGGACCCTGTGCAGCGAGAGCGCATCCTCGAGGTGTATCACCGCGTCGTCGACAACCACGAGCCGCTGCATCTGGTGGCCCACGACCTGAACC<br>GGCGTGGTGTCCTGTCGCCGAAGGACTACTTCGCGCAGCTGCAAGGCCGCGAGCCGCAGGGCCGGGAGTGGTCGGCTACCGCGCTGAAGCGATCGATGATCTCCGGGGCGATGCTCGGGTA<br>CGCGACTCTGAACGGTAAGACCGTCCGAGACGACGACGGAGCCCCGCTGGTGCGGGCTGAGCCGATCCTGACCCGTGAGCAGCTGGAGGCGCTGCGCGCCGAGCTCGTCAAGACCTCCCGG<br>GCGAAGCCCGCGGTGTCTACCCCGTCGCTGCTGCTGCGGGTGTTGTTCTGCGCGGTGTGCGGGGAGCCCGCGTACAAGTTCGCCGGGGGAGGACGTAAGCACCCGCGCTACCGCTGCCGCTC<br>GATGGGGTTCCCGAAGCACTGCGGGAACGGCACGGTGGCGATGGCCGAGTGGGACGCGTTCTGCGAGGAGCAGGTACTGGATCTGCTCGGGGACGCGGAGCGTCTGGAGAAAGTCTGGGT<br>AGCGGGCTCGGACTCCGCGGTCGAACTCGCGGAGGTGAACGCGGAGCTGGTGGACCTGACGTCGCTGATCGGCTCCCCGGCCTACCGGGCGGGCTCTCCGCAGCGAGAAGCACTGGATGCC<br>CGTATTGCGGCGCTGGCCGCGCGGCAAGAGGAGCTGGAGGGCCTGGAGGCTCGCCCGTCTGGCTGGGAGTGGCGCGAAACCGGGCAGCGGTTCGGGGACTGGTGGCGGGAGCAGGACAC<br>CGCGGCAAAGAACACCTGGCTTCGGTCGATGAACGTTCGGCTGACGTTCGACGTCCGCGGCGGGCTGACTCGCACGATCGACTTCGGGGATCTTCAGGAGTACGAGCAGCATCTCAGGCTCG<br>GCAGCGTGGTCGAACGGCTACACACCGGGATGTCGGCGGCTAACGACGAGAATTATGCATTGGCCGCCTAATAAAGGTCTAACGCATGAGAAAGCCCCCGGAAGATCACCTTCCGGGGGCTT<br>TTTTATTGCGCCCTTGAGAGCTTCCAGGATACATAGATTACCACAACTCCGAGCCC***TAGCGGCCGCTGCAGGCATGCCTCGAGATGCATGGCGCCT***AACCTAAACTGACAGGCATCAAATTAAG<br>CAGAAGGCCATCCTGACGGATGGCCTTTTTGCGTTTCGAACAATTGAAAAAACCTCGCGCCTTACCTGTTGAGTAATAGTCAAAAGCCTCCGGTCGGAGGCTTTTGACTTTCTGCTTACTGAATT<br>TCGGTGGTGCCGTTAATTAACCGGTGGGCCCTCATGATAATAATGGTTTCTTAGACGTCCGAAGTTCCTATTCTCTAGAAAGTATAGGAACTTCCATATGCCATGGGCGATGGTTCTGAGTAACA<br>GATAATAGAATGGCGATAAAGTGGCGGTAGAAATGGCGAATAATAGGTAATTATTGGCAAACAATGGCAATCTATGTCAATGATAAATAACGCATACTATTGATTTTCGGTTGTTCTTGTAGGA<br>ACTCATAATCGCTTGGTCGCTGGTTCAAGTCCAGCAAGGGCCACCAAATCATTATTTACGTATCAGGAAGAGTTAAGGGTACAAGGGGTACTCGATGGCGGTCTATATTTCATGACTAACGCCA<br>TTTCAAGGAAGAAGTGCGGTGGGTGGTTTCGAATGAG |
| Φ186 #2 | CCCTGGAGCCAAAATATCCTCAACCATATTTACTCCTTTCTAACATCACCAGGAAAAGTATAACGGATTGAAAATACATTGTTTATAAAAACAGCAGGCGCGCGGTAATGGCTGGATTGCGACA<br>CGGAGTTACTTTATAATCCGCTACCATGGCCCCTTAGCTCAGTGGTTAGAGCAGGCGACTCATAATCGCTTGGTCGCTGGTTCAAGTCCAGCAGGGGCCACCAAATTTAAGTAGTAAATACATA<br>CAGTTAGGCCACTCTCTCGAGTGGCCTTTTTGTTGCCTCGTTTTCGAGTGGCGATAGAATGGCGGTGGATTTTTTACCGCCACTTTCCTGAGGAGTGCATAGCTAGCTTCTTCGTCTGTTTCTACT<br>GGTATTGGCACAAACCTGATTCCAATTTGAGCAAGGCTATGTGCCATCTCGATACTCGTTCTTAACTCAACAGAAGATGCTTTGTGCATACAGCCCCTCGTTTATTATTTATCTCCTCAGCCAGCC<br>GCTGTGCTTTCAGTGGATTTCGGATAACAGAAAGGCCGGGAAATACCCAGCCTCGCTTTGTAACGGAGTAGAGACGAAAGTGATTGCGCCTACCCGGATATTATCGTGAGGATGCGTCATCGC<br>CATTAATTCACTGATCAGTGATAAGCTGTCAAACATGAGAATTGATCCGGCTGCCTCGCGCGTTTCGGTGATGACGGTGAAAACCTCTGACACATGCAGCTCCCGGAGACGGTCACAGCTTGTC<br>TGTAAGCGGATGCCGGGAGCAGACAAGCCCGTCAGGGCGCGTCAGCGGGTGTTGGCGGGTGTCGGGGCGCAGCCATGACCCAGTCACGTAGCGATAGCGGAGTGTATGCTGCACATGACAT<br>TAACCTATAAAAATAGGCGTATCACGAGGCCCTTTCGTCTTCAAGAATTAATTCCCAATTCCCCAGGCATCAAATAAAACGAAAGGCTCAGTCGAAAGACTGGGCCTTTCGTTTTATCTGTTGTTT<br>GTCGGTGAACGCTCTCCTGAGTAGGACAAATCCGCCGGGAGCGGATTTGAACGTTGCGAAGCAACGGCCCGGAGGGTGGCGGGCAGGACGCCCGCCATAAACTGCCAGGAATTAATTCCCCA<br>GGCATCAAATAAAACGAAAGGCTCAGTCGAAAGACTGGGCCTTTCGTTTTATCTGTTGTTTGTCGGTGAACGCTCTCCTGAGTAGGACAAATCCGCCGGGAGCGGATTTGAACGTTGCGAAGC<br>AACGGCCCGGAGGGTGGCGGGCAGGACGCCCGCCATAAACTGCCAGGAATTAATTCCCCAGGCATCAAATAAAACGAAAGGCTCAGTCGAAAGACTGGGCCTTTCGTTTTATCTGTTGTTTGT<br>CGGTGAACGCTCTCCTGAGTAGGACAAATCCGCCGGGAGCGGATTTGAACGTTGCGAAGCAACGGCCCGGAGGGTGGCGGGCAGGACGCCCGCCATAAACT***GCCAGGAATTGGGGATCGG***<br>***AATTCGAGCTCGGTACCCGGG***CTCGCATCCATTCTCAGGCTGTCTCGTCTCGTCTCGGAGAACGATCGTTGGCTGGGGGCCTCGCTTGGGTTATTGCTGGTGCCCGGCCGGGCGCAATATTCAT<br>GTTGATGATTTATTATATATCGAGTGGTGTATTTATTTATATTGTTTGCTCCGTTACCGTTATTAACTACTAGAGAAAGAGGAGAAATACTAAATGGTGAGAGCCCTGGTAGTCATCCGCCTGTCC<br>CGCGTCACCGATGCTACGACTTCACCGGAGCGTCAGCTGGAGTCTTGCCAGCAGCTCTGCGCCCAGCGCGGCTGGGACGTCGTCGGGGTAGCGGAGGATCTGGACGTCTCCGGGGCGGTCGA<br>TCCGTTCGACCGGAAGCGCAGACCGAACCTGGCCCGGTGGCTAGCGTTCGAGGAGCAACCGTTTGACGTGATCGTGGCGTACCGGGTAGATCGGTTGACCCGATCGATCCGGCATCTTCAGCA<br>GCTGGTCCACTGGGCCGAGGACCACAAGAAGCTGGTCGTCTCCGCGACCGAAGCGCACTTCGATACGACGACGCCGTTTGCGGCGGTCGTCATCGCGCTTATGGGAACGGTGGCGCAGATGG<br>AATTAGAAGCGATCAAAGAGCGGAACCGTTCGGCTGCGCATTTCAATATCCGCGCCGGGAAATACCGAGGATCCCTGCCGCCGTGGGGATACCTGCCTACGCGCGTGGACGGGGAGTGGCG<br>GCTGGTGCCGGACCCTGTGCAGCGAGAGCGCATCCTCGAGGTGTATCACCGCGTCGTCGACAACCACGAGCCGCTGCATCTGGTGGCCCACGACCTGAACCGGCGTGGTGTCCTGTCGCCGAA<br>GGACTACTTCGCGCAGCTGCAAGGCCGCGAGCCGCAGGGCCGGGAGTGGTCGGCTACCGCGCTGAAGCGATCGATGATCTCCGGGGCGATGCTCGGGTACGCGACTCTGAACGGTAAGACC<br>GTCCGAGACGACGACGGAGCCCCGCTGGTGCGGGCTGAGCCGATCCTGACCCGTGAGCAGCTGGAGGCGCTGCGCGCCGAGCTCGTCAAGACCTCCCGGGCGAAGCCCGCGGTGTCTACCC<br>CGTCGCTGCTGCTGCGGGTGTTGTTCTGCGCGGTGTGCGGGGAGCCCGCGTACAAGTTCGCCGGGGGAGGACGTAAGCACCCGCGCTACCGCTGCCGCTCGATGGGGTTCCCGAAGCACTGC<br>GGGAACGGCACGGTGGCGATGGCCGAGTGGGACGCGTTCTGCGAGGAGCAGGTACTGGATCTGCTCGGGGACGCGGAGCGTCTGGAGAAAGTCTGGGTAGCGGGCTCGGACTCCGCGGTC<br>GAACTCGCGGAGGTGAACGCGGAGCTGGTGGACCTGACGTCGCTGATCGGCTCCCCGGCCTACCGGGCGGGCTCTCCGCAGCGAGAAGCACTGGATGCCCGTATTGCGGCGCTGGCCGCGC<br>GGCAAGAGGAGCTGGAGGGCCTGGAGGCTCGCCCGTCTGGCTGGGAGTGGCGCGAAACCGGGCAGCGGTTCGGGGACTGGTGGCGGGAGCAGGACACCGCGGCAAAGAACACCTGGCTT<br>CGGTCGATGAACGTTCGGCTGACGTTCGACGTCCGCGGCGGGCTGACTCGCACGATCGACTTCGGGGATCTTCAGGAGTACGAGCAGCATCTCAGGCTCGGCAGCGTGGTCGAACGGCTACA<br>CACCGGGATGTCGGCGGCTAACGACGAGAATTATGCATTGGCCGCCTAATAAAGGTCTAACGCATGAGAAAGCCCCCGGAAGATCACCTTCCGGGGGCTTTTTTATTGCGCCCTTGAGAGCTT<br>CCAGGATACATAGATTACCACAACTCCGAGCCC***TAGCGGCCGCTGCAGGCATGCCTCGAGATGCATGGCGCCT***AACCTAAACTGACAGGCATCAAATTAAGCAGAAGGCCATCCTGACGGATG<br>GCCTTTTTGCGTTTCGAACAATTGAAAAAACCTCGCGCCTTACCTGTTGAGTAATAGTCAAAAGCCTCCGGTCGGAGGCTTTTGACTTTCTGCTTACTGAATTTCGGTGGTGCCGTTAATTAACCG<br>GTGGGCCCTCATGATAATAATGGTTTCTTAGACGTCCGAAGTTCCTATTCTCTAGAAAGTATAGGAACTTCCATATGCCATGGGCGATGGTTCTGAGTAACAGATAATAGAATGGCGATAAAGT<br>GGCGGTAGAAATGGCGAATAATAGGTAATTATTGGCAAACAATGGCAATCTATGTCAATGATAAATAACGCATACTATTGATTTTCGGTTGTTCTTGTAGGAACTCATAATCGCTTGGTCGCTG<br>GTTCAAGTCCAGCAGGGGCCACCAGATATAGCAAAGGCTGACGAGAAATCGTCAGCCTTTTTCTTTTTATATATCAGTTACTTTGCGTGCCAGTAAGCCGCTGCACGTACCCGCTGTGGGTCATA<br>CTGTTCCGCTTCAAAGCGGCGGCTTAAATTCTTAACGACTTTACCTTCGCCGGTTATCCAGATGAAGTAATCATCGGCAGGGATTTGCATCTGCGCCAGACGCGCATCTACCGCCTGCTCATCAT<br>GTGCCAGCCATTCGATATTAAAACCATCAAGGTGCGCGAGATAATCCTGACAGGCGTTATCCCGCACGCTAACCAGCGCACTAACTTGCGGTTTAACGGCAAGTTTGCTCAACGTTTCCAGGCG<br>GCGGCGCAATGCAGGCATTCCGGA |

Table S5. 1x differentiation strain with 2 copies salicylate inducible Bxb1-LAA. Promoters, RBSs, coding sequences, inserted bases in T7 RNAP CDS, terminators, integrase attachment sites, ***Gibson homologies***, genomic pOSIP primer binding sites

### eRWdiff2x

| Cluster | Sequence |
|---|---|
| P21 (T) | TAGAACTACCACCTGACCGGCCAGCCTACCCCAAAACTACCGAGGGGTTTTGATAACTTTTTTGATAAAATTTTGATAACCGTTAGCAGCCTAACAAAAAACAACGGGAGCGTTACGCTCCCGTT<br>AATAAATTTAACAAACTACGGCATTACATGTTTTCGATGATCGCGTCACCAAACTCTGAACATTTCAGCAGTTTAGCGCCATCCATCAGACGCTCGAAGTCATAGGTTACGGTTTTCGCGTTGATT<br>GCGCCTTCCATACCTTTAACAATTAAGTCAGCCGCTTCGGTCCAACCCATGTGGCGCAGCAGTATATAAAAATTATGTATTCAACTCGCTGATATTTAATCATAAATATTATTTTCTTGTCTTTGTA<br>TTAGCCAAAATCAACTTTATTCAACCGTTTGATTTCATTGACGTTCGTTTTTATTTTGGGGAAAGATTTTCTCTACCATTTATATCAAAATTACAAGTCTACATTTTATTAACATCTCAATACATGTGT<br>GGCCTTGCAAGCCATTGCGAGGCCTTATGCTAGCTTCTTCGTCTGTTTCTACTGGTATTGGCACAAACCTGATTCCAATTTGAGCAAGGCTATGTGCCATCTCGATACTCGTTCTTAACTCAACAG<br>AAGATGCTTTGTGCATACAGCCCCTCGTTTATTATTTATCTCCTCAGCCAGCCGCTGTGCTTTCAGTGGATTTCGGATAACAGAAAGGCCGGGAAATACCCAGCCTCGCTTTGTAACGGAGTAGA<br>GACGAAAGTGATTGCGCCTACCCGGATATTATCGTGAGGATGCGTCATCGCCATTAATTCACTGATCAGTGATAAGCTGTCAAACATGAGAATTGATCCGGCTGCCTCGCGCGTTTCGGTGATG<br>ACGGTGAAAACCTCTGACACATGCAGCTCCCGGAGACGGTCACAGCTTGTCTGTAAGCGGATGCCGGGAGCAGACAAGCCCGTCAGGGCGCGTCAGCGGGTGTTGGCGGGTGTCGGGGCGC<br>AGCCATGACCCAGTCACGTAGCGATAGCGGAGTGTATGCTGCACATGACATTAACCTATAAAAATAGGCGTATCACGAGGCCCTTTCGTCTTCAAGAATTAATTCCCAATTCCCCAGGCATCAAA<br>TAAAACGAAAGGCTCAGTCGAAAGACTGGGCCTTTCGTTTTATCTGTTGTTTGTCGGTGAACGCTCTCCTGAGTAGGACAAATCCGCCGGGAGCGGATTTGAACGTTGCGAAGCAACGGCCCG<br>GAGGGTGGCGGGCAGGACGCCCGCCATAAACTGCCAGGAATTAATTCCCCAGGCATCAAATAAAACGAAAGGCTCAGTCGAAAGACTGGGCCTTTCGTTTTATCTGTTGTTTGTCGGTGAACG<br>CTCTCCTGAGTAGGACAAATCCGCCGGGAGCGGATTTGAACGTTGCGAAGCAACGGCCCGGAGGGTGGCGGGCAGGACGCCCGCCATAAACTGCCAGGAATTAATTCCCCAGGCATCAAATA<br>AAACGAAAGGCTCAGTCGAAAGACTGGGCCTTTCGTTTTATCTGTTGTTTGTCGGTGAACGCTCTCCTGAGTAGGACAAATCCGCCGGGAGCGGATTTGAACGTTGCGAAGCAACGGCCCGGA<br>GGGTGGCGGGCAGGACGCCCGCCATAAACT***GCCAGGAATTGGGGATCGGAATTCGAGCTCGGTACCCGGG***CTCGCATCCATTCTCAGGCTGTCTCGTCTCGTCTCGGAGAACGATCGTTGGC<br>TGTGTTGACAATTAATCATCGGCTCGTATAATGTGTGGAATTGTGAGCGCTCACAATTTACTGAAAGATGCGATTTACTAAATGAACACGATTAACATCGCTAAGAACGACTTCTCTGACATCGA<br>ACTGGCTGCTATCCCGTTCAACACTCTGGCTGACCATTACGGTGAGCGTTTAGCTCGCGAACAGTTGGCCCTTGAGCATGAGTCTTACGAGATGGGTGAAGCACGCTTCCGCAAGATGTTTGAG<br>CGTCAACTTAAAGCTGGTGAGGTTGCGGATAACGCTGCCGCCAAGCCTCTCATCACTACCCTACTCCCTAAGATGATTGCACGCATCAACGACTGGTTTGAGGAAGTGAAAGCTAAGCGCGGCA<br>AGCGCCCGACAGCCTTCCAGTTCCTGCAAGAAATCAAGCCGGAAGCCGTAGCGTACATCACCATTAAGACCACTCTGGCTTGCCTAACCAGTGCTGACAATACAACCGTTCAGGCTGTAGCAAG<br>CGCAATCGGTCGGGCCATTGAGGACGAGGCTCGCTTCGGTCGTATCCGTGACCTTGAAGCTAAGCACTTCAAGAAAAACGTTGAGGAACAACTCAACAAGCGCGTAGGGCACGTCTACAAGG<br>GCTTGTCGACGACGGCGTGCTCCGTCGTCAGGATCATAGGTGACGAACAATAAGGCCTCCCTAACGGGGGGCCTTTTTTATTGATAACAAAAGCTT***GCACTGAAGGTCCTCAATCGCACTGGA***<br>***AACATCAAGGTCG***AAGCTCTCAAGGGCGCAATAAAAAAGCCCCCGGAAGGTGATCTTCCGGGGGCTTTCTCATGCGTTAGACCTTTACGGCAAACCATCAACTTGTTTCAGGTCTAACCCCCGT<br>TCGAAAATCTCATCAATCGGTAACATCTGACCGTCGGTCGTCATAAATTTGTGATCTTTCGTTGCACGGATAATCGAACCGTCTTCTAAGCAGTATTCGAAAACTTCCTGTTCTCCCCGGTTATGC<br>CACTGAGCAATAGGTTGTGTATACACAAAGCCATTTTTATCTACCGTATAGACTGTGCATTCAATCCGTTCCTCCACGATCTTACCGATAGGAAGGAATCCATATTCAACGGTCAGAATCTCTGTG<br>TCGTAAGACAGGCAGTTTTTAGCAGTAAAGGGCAGGTTAAGCTCTTTAGCTAAAGCAATGATATCATCCCCTCTTAGCGAAATTTTGCTGGCACCAAGTAATTTACCACCCTCTTTTAATTGTCGA<br>TAAGCAAGCGATGGGGTGATTTTGGCGAGCGCTGCAAGATCTTCAGCCCTAATTTTGAAAACTCGCCCTCGTTCAAGAGGTTCTTTGCTATCAATGGGAGCAAGCGCCATATACATCACTCGCTT<br>TGCGGGCAAAGGAAGTTGAGCCAGGGTGTGATTTAGCTCGTTTCGGTGGCGAATTTTCGTTTTTTTGTTCACGTCCATCATGACCTTGAGTCTCATTTAGTATTTCTCCTCTTTCTCTAGTATTCTT<br>CGAGCCTAGCAAGGGTCCGGGTTCACCGAAATCTATCTCATTTGCTAGTTCAGCCAACGATCGTTCTCC***CTGACCTCCTGCCAGCAATAGTAAGACAACACGCAAAGTC***GGAGGGAAACACAG<br>AAAAAAGCCCGCACCTGACAGTGCGGGCTTTTTTTTCGACCAAAGGTACTGGTTTGTCTGGTCAACCACCGCGTGCTCAGTGGTGTACGGTACAAACCAGGGTTCTAAAGCATTTATGCAAGT<br>TGTCGAGGCTGACATGCTCTCTAAGGGTCTACTCGGTGGCGAGGCGTGGTCTTCGTGGCATAAGGAAGACTCTATTCATGTAGGAGTACGCTGCATCGAGATGCTCATTGAGTCAACCGGAAT<br>GGTTAGCTTACACCGCCAAAATGCTGGCGTAGTAGGTCAAGACTCTGAGACTATCGAACTCGCACCTGAATACGCTGAGGCTATCGCAACCCGTGCAGGTGCGCTGGCTGGCATCTCTCCGAT<br>GTTCCAACCTTGCGTAGTTCCTCCTAAGCCGTGGACTGGCATTACTGGTGGTGGCTATTGGGCTAACGGTCGTCGTCCTCTGGCGCTGGTGCGTACTCACAGTAAGAAAGCACTGATGCGCTAC<br>GAAGACGTTTACATGCCTGAGGTGTACAAAGCGATTAACATTGCGCAAAACACCGCATGGAAAATCAACAAGAAAGTCCTAGCGGTCGCCAACGTAATCACCAAGTGGAAGCATTGTCCGGTC<br>GAGGACATCCCTGCGATTGAGCGTGAAGAACTCCCGATGAAACCGGAAGACATCGACATGAATCCTGAGGCTCTCACCGCGTGGAAACGTGCTGCCGCTGCTGTGTACCGCAAGGACAAGGC<br>TCGCAAGTCTCGCCGTATCAGCCTTGAGTTCATGCTTGAGCAAGCCAATAAGTTTGCTAACCATAAGGCCATCTGGTTCCCTTACAACATGGACTGGCGCGGTCGTGTTTACGCTGTGTCAATGT<br>TCAACCCGCAAGGTAACGATATGACCAAAGGACTGCTTACGCTGGCGAAAGGTAAACCAATCGGTAAGGAAGGTTACTACTGGCTGAAAATCCACGGTGCAAACTGTGCGGGTGTCGATAAG<br>GTTCCGTTCCCTGAGCGCATCAAGTTCATTGAGGAAAACCACGAGAACATCATGGCTTGCGCTAAGTCTCCACTGGAGAACACTTGGTGGGCTGAGCAAGATTCTCCGTTCTGCTTCCTTGCGTT<br>CTGCTTTGAGTACGCTGGGGTACAGCACCACGGCCTGAGCTATAACTGCTCCCTTCCGCTGGCGTTTGACGGGTCTTGCTCTGGCATCCAGCACTTCTCCGCGATGCTCCGAGATGAGGTAGGT<br>GGTCGCGCGGTTAACTTGCTTCCTAGTGAAACCGTTCAGGACATCTACGGGATTGTTGCTAAGAAAGTCAACGAGATTCTACAAGCAGACGCAATCAATGGGACCGATAACGAAGTAGTTACC<br>GTGACCGATGAGAACACTGGTGAAATCTCTGAGAAAGTCAAGCTGGGCACTAAGGCACTGGCTGGTCAATGGCTGGCTTACGGTGTTACTCGCAGTGTGACTAAGCGTTCAGTCATGACGCTG<br>GCTTACGGGTCCAAAGAGTTCGGCTTCCGTCAACAAGTGCTGGAAGATACCATTCAGCCAGCTATTGATTCCGGCAAGGGTCTGATGTTCACTCAGCCGAATCAGGCTGCTGGATACATGGCTA<br>AGCTGATTTGGGAATCTGTGAGCGTGACGGTGGTAGCTGCGGTTGAAGCAATGAACTGGCTTAAGTCTGCTGCTAAGCTGCTGGCTGCTGAGGTCAAAGATAAGAAGACTGGAGAGATTCTT<br>CGCAAGCGTTGCGCTGTGCATTGGGTAACTCCTGATGGTTTCCCTGTGTGGCAGGAATACAAGAAGCCTATTCAGACGCGCTTGAACCTGATGTTCCTCGGTCAGTTCCGCTTACAGCCTACCAT<br>TAACACCAACAAAGATAGCGAGATTGATGCACACAAACAGGAGTCTGGTATCGCTCCTAACTTTGTACACAGCCAAGACGGTAGCCACCTTCGTAAGACTGTAGTGTGGGCACACGAGAAGTA<br>CGGAATCGAATCTTTTGCACTGATTCACGACTCCTTCGGTACCATTCCGGCTGACGCTGCGAACCTGTTCAAAGCAGTGCGCGAAACTATGGTTGACACATATGAGTCTTGTGATGTACTGGCTG<br>ATTTCTACGACCAGTTCGCTGACCAGTTGCACGAGTCTCAATTGGACAAAATGCCAGCACTTCCGGCTAAAGGTAACTTGAACCTCCGTGACATCTTAGAGTCGGACTTCGCGTTCGCGTAATAA<br>AGGTTTCAGCCAAAAAACTTAAGACCGCCGGTCTTGTCCACTACCTTGCAGTAATGCGGTGGACAGGATCGGCGGTTTTCTTTTCTCTTCTCAAGCTT***GAGCCAACTCCCTTTACAACCTCACTCA***<br>***AGTCCGTTAGAG***GGAGTTTACGGCTAGCTCAGTCCTAGGTATAGTGCTAGCTACTAGAGAAAGAGGAGAAATACTAAATGGAACTGCGTGACCTGGATTTAAACCTGCTGGTGGTGTTCAACC<br>AGTTGCTGGTCGACAGACGCGTCTCTGTCACTGCGGAGAACCTGGGCCTGACCCAGCCTGCCGTGAGCAATGCGCTGAAACGCCTGCGCACCTCGCTACAGGACCCACTCTTCGTGCGCACAC<br>ATCAGGGAATGGAACCCACACCCTATGCCGCGCATCTGGCCGAGCACGTCACTTCGGCCATGCACGCACTGCGCAACGCCCTACAGCACCATGAAAGCTTCGATCCGCTGACCAGCGAGCGTA<br>CCTTCACCCTGGCCATGACCGACATTGGCGAGATCTACTTCATGCCGCGGCTGATGGATGCGCTGGCTCACCAGGCCCCCAATTGCGTGATCAGTACGGTGCGCGACAGTTCGATGAGCCTGAT<br>GCAGGCCTTGCAGAACGGAACCGTGGACTTGGCCGTGGGCCTGCTTCCCAATCTGCAAACTGGCTTCTTTCAGCGCCGGCTGCTCCAGAATCACTACGTGTGCCTATGTCGCAAGGACCATCCA<br>GTCACCCGCGAACCCCTGACTCTGGAGCGCTTCTGTTCCTACGGCCACGTGCGTGTCATCGCCGCTGGCACCGGCCACGGCGAGGTGGACACGTACATGACACGGGTCGGCATCCGGCGCGAC<br>ATCCGTCTGGAAGTGCCGCACTTCGCCGCCGTTGGCCACATCCTCCAGCGCACCGATCTGCTCGCCACTGTGCCGATATGTTTAGCCGACTGCTGCGTAGAGCCCTTCGGCCTAAGCGCCTTGCC<br>GCACCCAGTCGTCTTGCCTGAAATAGCCATCAACATGTTCTGGCATGCGAAGTACCACAAGGACCTAGCCAATATTTGGTTGCGGCAACTGATGTTTGACCTGTTTACGGATTGATACTGGGCCC<br>AAGTTCACTTAAAAAGGAGATCAACAATGAAAGCAATTTTCGTACTGAAACATCTTAATCATGCTAAGGAGGTTTTCTAATGGCCTTGGTTGACGGTTTTCTTGAGCTGGAACGCTCAAGTGGA<br>AAATTGGAGTTGAGCGCCATCCTGCAGAAGATGGCGAGCGACCTCGGATTCTCGAAGATCCTGTTCGGCCTGTTGCCTAAGGACAGCCAGGACTACGAGAACGCCTTCATCGTCAGCAACTAC<br>CCGGCCGCCTGGCGCGAGCATTACGACCGAACTGGCTACGCGCGGGTCGACCCGACGGTCAGTCACTGTACCCAGAGCGTACTGCCGATTTTCTGGGAACCGCCCATCTACCAGACGCGAAAG<br>CAGCACGAGTTCTTCGAGGAAGCCTTGGCCGCTGGCCTGGTGTATGGACTGACCATGCCGCTGCATGGTGCTCGCGGCGAACTCGGCGCGCTGAGCCTCAGCGTGGAAGCGGAAAACCGGGC<br>CGAGGCCAACCGTTTCATGGAGTCGGTCCTGCCGACCCTGTGGATGCTCAAGGACTACGCACTGCAGAGCGGTGTCGGAATGGCCTTCGAACATCCGATCAGCAAACCGGTGGTTCTGACCAG<br>CCGAGAGAAGGAAGTGTTACAGTGGTGCGCCATCGGCAAGACCAGCTGGGAGATATCGGTTATCTGCAACTGCTCGGAAGCCAATGTGAACTTCCATATGGGAAATATTCGGCGGAAGTTCG<br>GTGTGACCTCCCGCCGCGTAGCGGCCATTATGGCCGTTAATTTGGGTCTTATTACTCTCTGAGGTCCAATTATTGAACACCCTTCGGGGTGTTTTTTTGTTTCTGGTCTGCCGCTT***CTCGTTCGCTG***<br>***CCACCTAAGAATACTCTACGGTCACATAC***GGAGGACACCATCGAATGGCGCAAAACCTTTCGCGGTATGGCATGATAGCGCCCGGAAGAGAGTCAATTCAGGGTGGTGAATAATGAAACCAG<br>TAACGTTATACGATGTCGCAGAGTATGCCGGTGTCTCTTATATGACCGTTTCCCGCGTGGTGAACCAGGCCAGCCACGTTTCTGCGAAAACGCGGGAAAAAGTGGAAGCGGCGATGGTGGAG<br>CTGAATTACATTCCCAACCGCGTGGCACAACAACTGGCGGGCAAACAGTCGTTGCTGATTGGCGTTGCCACCTCCAGTCTGGCCCTGCACGCGCCGTCGCAAATTGTCGCGGCGATTAAATCTC<br>GCGCCGATCAACTGGGTGCCAGCGTGGTGGTGTCGATGGTAGAACGAAGCGGCGTCGAAGCCTGTAAAGCGGCGGTGCACAATCTTCTCGCGCAACGCGTCAGTGGGCTGATCATTAACTAT<br>CCGCTGGATGACCAGGATGCCATTGCTGTGGAAGCTGCCTGCACTAATGTTCCGGCGTTATTTCTTGATGTCTCTGACCAGACACCCATCAACAGTATTATTTTCTCCCATGAGGACGGTACGCG<br>ACTGGGCGTGGAGCATCTGGTCGCATTGGGTCACCAGCAAATCGCGCTGTTAGCGGGCCCATTAAGTTCTGTCTCGGCGCGTCTGCGTCTGGCTGGCTGGCATAAATATCTCACTCGCAATCAA<br>ATTCAGCCGATAGCGGAACGGGAAGGCGACTGGAGTGCCATGTCCGGTTTTCAACAAACCATGCAAATGCTGAATGAGGGCATCGTTCCCACTGCGATGCTGGTTGCCAACGATCAGATGGC<br>GCTGGGCGCAATGCGCGCCATTACCGAGTCCGGGCTGCGCGTTGGTGCGGATATCTCGGTAGTGGGATACGACGATACCGAAGATAGCTCATGTTATATCCCGCCGTTAACCACCATCAAACA<br>GGATTTTCGCCTGCTGGGGCAAACCAGCGTGGACCGCTTGCTGCAACTCTCTCAGGGCCAGGCGGTGAAGGGCAATCAGCTGTTGCCAGTCTCACTGGTGAAAAGAAAAACCACCCTGGCGCC<br>CAATACGCAAACCGCCTCTCCCCGCGCGTTGGCCGATTCATTAATGCAGCTGGCACGACAGGTTTCCCGACTGGAAAGCGGGCAGTGAGGTTCTCGGTACCAAATTCCAGAAAAGAGACGCTT<br>TCGAGCGTCTTTTTTCGTTTTGGTCCGCTT***TAGCGGCCGCTGCAGGCATGCCTCGAGATGCATGGCGCCT***AACCTAAACTGACAGGCATCAAATTAAGCAGAAGGCCATCCTGACGGATGGCCT<br>TTTTGCGTTTCGAACAATTGAAAAAACCTCGCGCCTTACCTGTTGAGTAATAGTCAAAAGCCTCCGGTCGGAGGCTTTTGACTTTCTGCTTACTGAATTTCGGTGGTGCCGTTAATTAACCGGTG<br>GGCCCTCATGATAATAATGGTTTCTTAGACGTCCGAAGTTCCTATTCTCTAGAAAGTATAGGAACTTCCATATGCCATGGAATGACCGACCGATACAATGATGATCGTGGTAAAGACTGGATTAT<br>CGTAGATATCAAAACAGCATAGAAAATAGCCAGTTTTGGGGAAGGGTTTTGGGGAAAGTTTTGGGGAAGATTTTACATCATCATAAAACAACGGGCGTGTTATACGCCCGTTATAATATTTAAC<br>ACATGTGGTAATTACATATTTTTGATGATCGCGTCACCAAACTCGCTGCATTTCAGCAGTTTAGCGCCTTCCATCAGACGTTCGAAGTCATAGGTTACGGTCTTGGCGTTGATTGCGCCTTCCATA<br>CCTTTAACAATCAGGTCTGCGGCTTCAGTCCAGCCCATATGGCGCAGCATCATCTCAGCGGAGAGAATAATAGAGCCAGGATTTACTTTGTCCTGACCGGCATATTTCGGCGCAGTACCGTGGG<br>TGGCTTCAAACAGGGCGCATTCGTCACCGATGTTTGCACCAGGGGCGATACCGATACCGCCAACCTGCGCTGCCAGGGCGTCAGAAATGTAGTCACCGTTCAGGTTCATACAGGCGAT |

HK022

GGCATCAACAGCACATTCAGTGGTTTACCGTGCGCCCATTGCACCAGATATCGCCCCGGTTGATAGCCCATCTGAAACCAGGGCACACCAACGCGGCTTTTCACCTGGGGAGCATCAATAGCAT
TTACCAGTTCGATCACCGGCAGACTTGCTACCTGCTTTTGCAGGTCGGGAAATGAGGTCGTGCTACTACCGAGCAAAATGGCCTCTGCGCCCCACTGTTTACACTGGTCGATTTGTGCTTGCTGG
GTAGCCAACTGGCTGTAGCCGCCTGCCTCCAGCACTTTTAAATCCACACCGTAGCGGCGAGCAGCCTCCTGCATACCATAGTTCAACGATAACCAATATGAATCTTTCAGGCTGGGATAAAGCG
CGCACAGTTTCCATGCGCGTTTGGCTTTAAGCGGCGTAGAGGCTTGCACCGTGAAATGCTGCGCATCATGCCAGCGCAACAGGTTATCAGCCGAAAATGCCGGCAACATGAAAAGGGAAAGA
AGTAAAAATAGCAGTACGCGCATGATAGCCTCATCAATAATAAGGCTTTATGCTAGATGCATTCCGCTTTGCGACTCAACCTTTTTCACCTAAAGGATGACAAAATAACATTAATCACTTAAAAA
TCATCGCATTACACTAATCTGTGGTTAAATGATAGACTACATAATGCGACAAAACGCAACATATCCAGTCACTATGAATCAACTACTTAGATAGTATTAGTGACCTGAGACAGAGCATTAGCTAG
CTTCTTCGTCTGTTTCTACTGGTATTGGCACAAACCTGATTCCAATTTGAGCAAGGCTATGTGCCATCTCGATACTCGTTCTTAACTCAACAGAAGATGCTTTGTGCATACAGCCCCTCGTTTATTA
TTTATCTCCTCAGCCAGCCGCTGTGCTTTCAGTGGATTTCGGATAACAGAAAGGCCGGGAAATACCCAGCCTCGCTTTGTAACGGAGTAGAGACGAAAGTGATTGCGCCTACCCGGATATTATC
GTGAGGATGCGTCATCGCCATTAATTCACTGATCAGTGATAAGCTGTCAAACATGAGAATTGATCCGGCTGCCTCGCGCGTTTCGGTGATGACGGTGAAAACCTCTGACACATGCAGCTCCCGG
AGACGGTCACAGCTTGTCTGTAAGCGGATGCCGGGAGCAGACAAGCCCGTCAGGGCGCGTCAGCGGGTGTTGGCGGGTGTCGGGGCGCAGCCATGACCCAGTCACGTAGCGATAGCGGAGT
GTATGCTGCACATGACATTAACCTATAAAAATAGGCGTATCACGAGGCCCTTTCGTCTTCAAGAATTAATTCCCAATTCCCCAGGCATCAAATAAAACGAAAGGCTCAGTCGAAAGACTGGGCC
TTTCGTTTTATCTGTTGTTTGTCGGTGAACGCTCTCCTGAGTAGGACAAATCCGCCGGGAGCGGATTTGAACGTTGCGAAGCAACGGCCCGGAGGGTGGCGGGCAGGACGCCCGCCATAAACT
GCCAGGAATTAATTCCCCAGGCATCAAATAAAACGAAAGGCTCAGTCGAAAGACTGGGCCTTTCGTTTTATCTGTTGTTTGTCGGTGAACGCTCTCCTGAGTAGGACAAATCCGCCGGGAGCGG
ATTTGAACGTTGCGAAGCAACGGCCCGGAGGGTGGCGGGCAGGACGCCCGCCATAAACT***GCCAGGAATTGGGGATCGGAATTCGAGCTCGGTACCCGGG***CTCGCATCCATTCTCAGGCTGTC
TCGTCTCGTCTCGGAGAACGATCGTTGGCTGTGTTGACAATTAATCATCGGCTCGTATAATGTGTGGAATTGTGAGCGCTCACAATTTACTGAAAGATGCGATTTACTAAATGAACACGATTAAC
ATCGCTAAGAACGACTTCTCTGACATCGAACTGGCTGCTATCCCGTTCAACACTCTGGCTGACCATTACGGTGAGCGTTTAGCTCGCGAACAGTTGGCCCTTGAGCATGAGTCTTACGAGATGG
GTGAAGCACGCTTCCGCAAGATGTTTGAGCGTCAACTTAAAGCTGGTGAGGTTGCGGATAACGCTGCCGCCAAGCCTCTCATCACTACCCTACTCCCTAAGATGATTGCACGCATCAACGACTG
GTTTGAGGAAGTGAAAGCTAAGCGCGGCAAGCGCCCGACAGCCTTCCAGTTCCTGCAAGAAATCAAGCCGGAAGCCGTAGCGTACATCACCATTAAGACCACTCTGGCTTGCCTAACCAGTGC
TGACAATACAACCGTTCAGGCTGTAGCAAGCGCAATCGGTCGGGCCATTGAGGACGAGGCTCGCTTCGGTCGTATCCGTGACCTTGAAGCTAAGCACTTCAAGAAAAACGTTGAGGAACAACT
CAACAAGCGCGTAGGGCACGTCTACAAGGGCTTGTCGACGACGGCGTGCTCCGTCGTCAGGATCATAGGTGACGAACAATAAGGCCTCCCTAACGGGGGGCCTTTTTTATTGATAACAAAAGC
TT***GCACTGAAGGTCCTCAATCGCACTGGAAACATCAAGGTCG***AAGCTCTCAAGGGCGCAATAAAAAAGCCCCCGGAAGGTGATCTTCCGGGGGCTTTCTCATGCGTTAGACCTTTACCCCTTA
GCTTTTTGGGAGGTACGGTTTCATCAAATACCTTGAGAAAAGCTGCATCAGCTTCAGATAAATTCATAAAAAAAGCTTCATCATCTTTATCGCCAGAAAATTCATCTTCATCAACGACAAATTCG
AACTTCAGCTTACTAATTTTTCTTCCTTCTTTTTCATGAACAGTGAAGCCAACAAACGATATTTCTGTTTTCTTTTTAATTTCAGCAATGGCTTTATTTAACACATCCCTTTTAAAAATAGGAAAGTCA
GGGTATTTGTACTCAATATTTCCATCTTTATCAAAAGTATAAGCTATTAACTCTTCCTTTAACTCATCAACGGAAATAATAAAATAATTTTTCTTCTTAAAATTAGAGTAATGCTTCCTGATAAGTTG
ATAAAGAGAAGATGAATACTGGCTACTTAAGCGTAAGCTTGCCGTTAACAATTGCGTTGTGAATTTATTTTTTTTCCCAATAAGGCTAGAGATATATGGTTCTATGGTTCTGGTGAATTTTAAAG
ACAAGTATCCTTCATCATTTGAATAAGCTATCCACTCAATAATGTTAAGATCTAACTCTTCAGGGGAGTTAGAGGCCACCAGCCCATTCTTCAGCAAAAAATTATGATCTTTTTCTACTCCAATATC
GTAGACGTTTTGAGTGCCCAGACTCTTACGACTAATGATCTTGACCATTTAGTATTTCTCCTCTTTCTCTAGTATTCTTCGAGCCTAGCAAGGGTCCGGGTTCACCGAAATCTATCTCATTTGCTAG
TTCAGCCAACGATCGTTCTCC***CTGACCTCCTGCCAGCAATAGTAAGACAACACGCAAAGTC***GGAGGGAAACACAGAAAAAAGCCCGCACCTGACAGTGCGGGCTTTTTTTTTCGACCAAAGGT
ACTGGTTTGTCTGGTCAACCACCGCGTGCTCAGTGGTGTACGGTACAAACCAGGGTTCTAAAGCATTTATGCAAGTTGTCGAGGCTGACATGCTCTCTAAGGGTCTACTCGGTGGCGAGGCGTG
GTCTTCGTGGCATAAGGAAGACTCTATTCATGTAGGAGTACGCTGCATCGAGATGCTCATTGAGTCAACCGGAATGGTTAGCTTACACCGCCAAAATGCTGGCGTAGTAGGTCAAGACTCTGA
GACTATCGAACTCGCACCTGAATACGCTGAGGCTATCGCAACCCGTGCAGGTGCGCTGGCTGGCATCTCTCCGATGTTCCAACCTTGCGTAGTTCCTCCTAAGCCGTGGACTGGCATTACTGGT
GGTGGCTATTGGGCTAACGGTCGTCGTCCTCTGGCGCTGGTGCGTACTCACAGTAAGAAAGCACTGATGCGCTACGAAGACGTTTACATGCCTGAGGTGTACAAAGCGATTAACATTGCGCAA
AACACCGCATGGAAAATCAACAAGAAAGTCCTAGCGGTCGCCAACGTAATCACCAAGTGGAAGCATTGTCCGGTCGAGGACATCCCTGCGATTGAGCGTGAAGAACTCCCGATGAAACCGGA
AGACATCGACATGAATCCTGAGGCTCTCACCGCGTGGAAACGTGCTGCCGCTGCTGTGTACCGCAAGGACAAGGCTCGCAAGTCTCGCCGTATCAGCCTTGAGTTCATGCTTGAGCAAGCCAA
TAAGTTTGCTAACCATAAGGCCATCTGGTTCCCTTACAACATGGACTGGCGCGGTCGTGTTTACGCTGTGTCAATGTTCAACCCGCAAGGTAACGATATGACCAAAGGACTGCTTACGCTGGCG
AAAGGTAAACCAATCGGTAAGGAAGGTTACTACTGGCTGAAAATCCACGGTGCAAACTGTGCGGGTGTCGATAAGGTTCCGTTCCCTGAGCGCATCAAGTTCATTGAGGAAAACCACGAGAAC
ATCATGGCTTGCGCTAAGTCTCCACTGGAGAACACTTGGTGGGCTGAGCAAGATTCTCCGTTCTGCTTCCTTGCGTTCTGCTTTGAGTACGCTGGGGTACAGCACCACGGCCTGAGCTATAACTG
CTCCCTTCCGCTGGCGTTTGACGGGTCTTGCTCTGGCATCCAGCACTTCTCCGCGATGCTCCGAGATGAGGTAGGTGGTCGCGCGGTTAACTTGCTTCCTAGTGAAACCGTTCAGGACATCTACG
GGATTGTTGCTAAGAAAGTCAACGAGATTCTACAAGCAGACGCAATCAATGGGACCGATAACGAAGTAGTTACCGTGACCGATGAGAACACTGGTGAAATCTCTGAGAAAGTCAAGCTGGGC
ACTAAGGCACTGGCTGGTCAATGGCTGGCTTACGGTGTTACTCGCAGTGTGACTAAGCGTTCAGTCATGACGCTGGCTTACGGGTCCAAAGAGTTCGGCTTCCGTCAACAAGTGCTGGAAGAT
ACCATTCAGCCAGCTATTGATTCCGGCAAGGGTCTGATGTTCACTCAGCCGAATCAGGCTGCTGGATACATGGCTAAGCTGATTTGGGAATCTGTGAGCGTGACGGTGGTAGCTGCGGTTGAA
GCAATGAACTGGCTTAAGTCTGCTGCTAAGCTGCTGGCTGCTGAGGTCAAAGATAAGAAGACTGGAGAGATTCTTCGCAAGCGTTGCGCTGTGCATTGGGTAACTCCTGATGGTTTCCCTGTGT
GGCAGGAATACAAGAAGCCTATTCAGACGCGCTTGAACCTGATGTTCCTCGGTCAGTTCCGCTTACAGCCTACCATTAACACCAACAAAGATAGCGAGATTGATGCACACAAACAGGAGTCTG
GTATCGCTCCTAACTTTGTACACAGCCAAGACGGTAGCCACCTTCGTAAGACTGTAGTGTGGGCACACGAGAAGTACGGAATCGAATCTTTTGCACTGATTCACGACTCCTTCGGTACCATTCCG
GCTGACGCTGCGAACCTGTTCAAAGCAGTGCGCGAAACTATGGTTGACACATATGAGTCTTGTGATGTACTGGCTGATTTCTACGACCAGTTCGCTGACCAGTTGCACGAGTCTCAATTGGACA
AAATGCCAGCACTTCCGGCTAAAGGTAACTTGAACCTCCGTGACATCTTAGAGTCGGACTTCGCGTTCGCGTAATAAAGGTTTCAGCCAAAAAACTTAAGACCGCCGGTCTTGTCCACTACCTTG
CAGTAATGCGGTGGACAGGATCGGCGGTTTTCTTTTCTCTTCTCAAGCTT***GAGCCAACTCCCTTTACAACCTCACTCAAGTCCGTTAGAG***GGAGTTTACGGCTAGCTCAGTCCTAGGTATAGTGC
TAGCTACTAGAGAAAGAGGAGAAATACTAAATGGAACTGCGTGACCTGGATTTAAACCTGCTGGTGGTGTTCAACCAGTTGCTGGTCGACAGACGCGTCTCTGTCACTGCGGAGAACCTGGGC
CTGACCCAGCCTGCCGTGAGCAATGCGCTGAAACGCCTGCGCACCTCGCTACAGGACCCACTCTTCGTGCGCACACATCAGGGAATGGAACCCACACCCTATGCCGCGCATCTGGCCGAGCAC
GTCACTTCGGCCATGCACGCACTGCGCAACGCCCTACAGCACCATGAAAGCTTCGATCCGCTGACCAGCGAGCGTACCTTCACCCTGGCCATGACCGACATTGGCGAGATCTACTTCATGCCGC
GGCTGATGGATGCGCTGGCTCACCAGGCCCCCAATTGCGTGATCAGTACGGTGCGCGACAGTTCGATGAGCCTGATGCAGGCCTTGCAGAACGGAACCGTGGACTTGGCCGTGGGCCTGCTT
CCCAATCTGCAAACTGGCTTCTTTCAGCGCCGGCTGCTCCAGAATCACTACGTGTGCCTATGTCGCAAGGACCATCCAGTCACCCGCGAACCCCTGACTCTGGAGCGCTTCTGTTCCTACGGCCA
CGTGCGTGTCATCGCCGCTGGCACCGGCCACGGCGAGGTGGACACGTACATGACACGGGTCGGCATCCGGCGCGACATCCGTCTGGAAGTGCCGCACTTCGCCGCCGTTGGCCACATCCTCCA
GCGCACCGATCTGCTCGCCACTGTGCCGATATGTTTAGCCGACTGCTGCGTAGAGCCCTTCGGCCTAAGCGCCTTGCCGCACCCAGTCGTCTTGCCTGAAATAGCCATCAACATGTTCTGGCATG
CGAAGTACCACAAGGACCTAGCCAATATTTGGTTGCGGCAACTGATGTTTGACCTGTTTACGGATTGATACTGGGCCCAAGTTCACTTAAAAAGGAGATCAACAATGAAAGCAATTTTCGTACT
GAAACATCTTAATCATGCTAAGGAGGTTTTCTAATGGCCTTGGTTGACGGTTTTCTTGAGCTGGAACGCTCAAGTGGAAAATTGGAGTTGAGCGCCATCCTGCAGAAGATGGCGAGCGACCTC
GGATTCTCGAAGATCCTGTTCGGCCTGTTGCCTAAGGACAGCCAGGACTACGAGAACGCCTTCATCGTCAGCAACTACCCGGCCGCCTGGCGCGAGCATTACGACCGAACTGGCTACGCGCGG
GTCGACCCGACGGTCAGTCACTGTACCCAGAGCGTACTGCCGATTTTCTGGGAACCGCCCATCTACCAGACGCGAAAGCAGCACGAGTTCTTCGAGGAAGCCTTGGCCGCTGGCCTGGTGTAT
GGACTGACCATGCCGCTGCATGGTGCTCGCGGCGAACTCGGCGCGCTGAGCCTCAGCGTGGAAGCGGAAAACCGGGCCGAGGCCAACCGTTTCATGGAGTCGGTCCTGCCGACCCTGTGGAT
GCTCAAGGACTACGCACTGCAGAGCGGTGTCGGAATGGCCTTCGAACATCCGATCAGCAAACCGGTGGTTCTGACCAGCCGAGAGAAGGAAGTGTTACAGTGGTGCGCCATCGGCAAGACCA
GCTGGGAGATATCGGTTATCTGCAACTGCTCGGAAGCCAATGTGAACTTCCATATGGGAAATATTCGGCGGAAGTTCGGTGTGACCTCCCGCCGCGTAGCGGCCATTATGGCCGTTAATTTGG
GTCTTATTACTCTCTGAGGTCCAATTATTGAACACCCTTCGGGGTGTTTTTTTGTTTCTGGTCTGCCGCTT***CTCGTTCGCTGCCACCTAAGAATACTCTACGGTCACATAC***GGAGGACACCATCGA
ATGGCGCAAAACCTTTCGCGGTATGGCATGATAGCGCCCGGAAGAGAGTCAATTCAGGGTGGTGAATAATGAAACCAGTAACGTTATACGATGTCGCAGAGTATGCCGGTGTCTCTTATATGA
CCGTTTCCCGCGTGGTGAACCAGGCCAGCCACGTTTCTGCGAAAACGCGGGAAAAAGTGGAAGCGGCGATGGTGGAGCTGAATTACATTCCCAACCGCGTGGCACAACAACTGGCGGGCAAA
CAGTCGTTGCTGATTGGCGTTGCCACCTCCAGTCTGGCCCTGCACGCGCCGTCGCAAATTGTCGCGGCGATTAAATCTCGCGCCGATCAACTGGGTGCCAGCGTGGTGGTGTCGATGGTAGAA
CGAAGCGGCGTCGAAGCCTGTAAAGCGGCGGTGCACAATCTTCTCGCGCAACGCGTCAGTGGGCTGATCATTAACTATCCGCTGGATGACCAGGATGCCATTGCTGTGGAAGCTGCCTGCACT
AATGTTCCGGCGTTATTTCTTGATGTCTCTGACCAGACACCCATCAACAGTATTATTTACTCCCATGAGGACGGTACGCGACTGGGCGTGGAGCATCTGGTCGCATTGGGTCACCAGCAAATCGC
GCTGTTAGCGGGCCCATTAAGTTCTGTCTCGGCGCGTCTGCGTCTGGCTGGCTGGCATAAATATCTCACTCGCAATCAAATTCAGCCGATAGCGGAACGGGAAGGCGACTGGAGTGCCATGTC
CGGTTTTCAACAAACCATGCAAATGCTGAATGAGGGCATCGTTCCCACTGCGATGCTGGTTGCCAACGATCAGATGGCGCTGGGCGCAATGCGCGCCATTACCGAGTCCGGGCTGCGCGTTGG
TGCGGATATCTCGGTAGTGGGATACGACGATACCGAAGATAGCTCATGTTATATCCCGCCGTTAACCACCATCAAACAGGATTTTCGCCTGCTGGGGCAAACCAGCGTGGACCGCTTGCTGCAA
CTCTCTCAGGGCCAGGCGGTGAAGGGCAATCAGCTGTTGCCAGTCTCACTGGTGAAAAGAAAAACCACCCTGGCGCCCAATACGCAAACCGCCTCTCCCCGCGCGTTGGCCGATTCATTAATGC
AGCTGGCACGACAGGTTTCCCGACTGGAAAGCGGGCAGTGAGGTTCTCGGTACCAAATTCCAGAAAAGAGACGCTTTCGAGCGTCTTTTTTCGTTTTGGTCCGCTT***TAGCGGCCGCTGCAGGC***
***ATGCCTCGAGATGCATGGCGCCT***AACCTAAACTGACAGGCATCAAATTAAGCAGAAGGCCATCCTGACGGATGGCCTTTTTGCGTTTCGAACAATTGAAAAAACCTCGCGCCTTACCTGTTGAG
TAATAGTCAAAAGCCTCCGGTCGGAGGCTTTTGACTTTCTGCTTACTGAATTTCGGTGGTGCCGTTAATTAACCGGTGGGCCCTCATGATAATAATGGTTTCTTAGACGTCCGAAGTTCCTATTCT
CTAGAAAGTATAGGAACTTCCATATGCCATGGGACAAAATTGAAATCGACAAATGATTTTATTTTGACTAATAATGACCTACTTACATTAATTTACTGATAATTAAAGAGATTTTAAATATACAAC
TTATTCACCTAAAGTGCACCGACCGTGAATTTAACCCTGACCCGAAGACTCTGGATGGGCTTTGCCCTGATGGCGCTGTTAACCCTGACCAGTACCCTGGTGGGATGGTACAACCTGCGCTTTAT
CAGCCAGGTGGAAAAAGACAACACTCAGGCATTGATTCC

| | |
|---|---|
| Φ186 #1 | GATCATCATGTTTATTGCGTGGTTTATAAGTATTGTTATTGCACACTCAAGGCCCTTTAGCTCAGTGGTTAGAGCAGGCGACTCATAATCGCTTGGTCGCTGGTTCAAGTCCAGCAGGGGCCACC<br>AAATTTAAGTAGTAAATACATACAGTTAGGCCACTCTCTCGAGTGGCCTTTTTGTTGCCTCGTTTTCGAGTGGCGATAGAATGGCGGTGGATTTTTTACCGCCACTTTCCTGAGGAGTGCATAGC<br>TAGCTTCTTCGTCTGTTTCTACTGGTATTGGCACAAACCTGATTCCAATTTGAGCAAGGCTATGTGCCATCTCGATACTCGTTCTTAACTCAACAGAAGATGCTTTGTGCATACAGCCCCTCGTTTA<br>TTATTTATCTCCTCAGCCAGCCGCTGTGCTTTCAGTGGATTTCGGATAACAGAAAGGCCGGGAAATACCCAGCCTCGCTTTGTAACGGAGTAGAGACGAAAGTGATTGCGCCTACCCGGATATT<br>ATCGTGAGGATGCGTCATCGCCATTAATTCACTGATCAGTGATAAGCTGTCAAACATGAGAATTGATCCGGCTGCCTCGCGCGTTTCGGTGATGACGGTGAAAACCTCTGACACATGCAGCTCC<br>CGGAGACGGTCACAGCTTGTCTGTAAGCGGATGCCGGGAGCAGACAAGCCCGTCAGGGCGCGTCAGCGGGTGTTGGCGGGTGTCGGGGCGCAGCCATGACCCAGTCACGTAGCGATAGCG<br>GAGTGTATGCTGCACATGACATTAACCTATAAAAATAGGCGTATCACGAGGCCCTTTCGTCTTCAAGAATTAATTCCCAATTCCCCAGGCATCAAATAAAACGAAAGGCTCAGTCGAAAGACTG<br>GGCCTTTCGTTTTATCTGTTGTTTGTCGGTGAACGCTCTCCTGAGTAGGACAAATCCGCCGGGAGCGGATTTGAACGTTGCGAAGCAACGGCCCGGAGGGTGGCGGGCAGGACGCCCGCCATA<br>AACTGCCAGGAATTAATTCCCCAGGCATCAAATAAAACGAAAGGCTCAGTCGAAAGACTGGGCCTTTCGTTTTATCTGTTGTTTGTCGGTGAACGCTCTCCTGAGTAGGACAAATCCGCCGGGA<br>GCGGATTTGAACGTTGCGAAGCAACGGCCCGGAGGGTGGCGGGCAGGACGCCCGCCATAAACTGCCAGGAATTAATTCCCCAGGCATCAAATAAAACGAAAGGCTCAGTCGAAAGACTGGG<br>CCTTTCGTTTTATCTGTTGTTTGTCGGTGAACGCTCTCCTGAGTAGGACAAATCCGCCGGGAGCGGATTTGAACGTTGCGAAGCAACGGCCCGGAGGGTGGCGGGCAGGACGCCCGCCATAAA<br>CT***GCCAGGAATTGGGGATCGGAATTCGAGCTCGGTACCCGGG***CTCGCATCCATTCTCAGGCTGTCTCGTCTCGTCTCGGAGAACGATCGTTGGCTGGGGGCCTCGCTTGGGTTATTGCTGGTG<br>CCCGGCCGGGCGCAATATTCATGTTGATGATTTATTATATATCGAGTGGTGTATTTATTTATATTGTTTGCTCCGTTACCGTTATTAACTACTAGAGAAAGAGGAGAAATACTAAATGGTGAGAG<br>CCCTGGTAGTCATCCGCCTGTCCCGCGTCACCGATGCTACGACTTCACCGGAGCGTCAGCTGGAGTCTTGCCAGCAGCTCTGCGCCCAGCGCGGCTGGGACGTCGTCGGGGTAGCGGAGGATC<br>TGGACGTCTCCGGGGCGGTCGATCCGTTCGACCGGAAGCGCAGACCGAACCTGGCCCGGTGGCTAGCGTTCGAGGAGCAACCGTTTGACGTGATCGTGGCGTACCGGGTAGATCGGTTGACC<br>CGATCGATCCGGCATCTTCAGCAGCTGGTCCACTGGGCCGAGGACCACAAGAAGCTGGTCGTCTCCGCGACCGAAGCGCACTTCGATACGACGACGCCGTTTGCGGCGGTCGTCATCGCGCTT<br>ATGGGAACGGTGGCGCAGATGGAATTAGAAGCGATCAAAGAGCGGAACCGTTCGGCTGCGCATTTCAATATCCGCGCCGGGAAATACCGAGGATCCCTGCCGCCGTGGGGATACCTGCCTAC<br>GCGCGTGGACGGGGAGTGGCGGCTGGTGCCGGACCCTGTGCAGCGAGAGCGCATCCTCGAGGTGTATCACCGCGTCGTCGACAACCACGAGCCGCTGCATCTGGTGGCCCACGACCTGAACC<br>GGCGTGGTGTCCTGTCGCCGAAGGACTACTTCGCGCAGCTGCAAGGCCGCGAGCCGCAGGGCCGGGAGTGGTCGGCTACCGCGCTGAAGCGATCGATGATCTCCGGGGCGATGCTCGGGTA<br>CGCGACTCTGAACGGTAAGACCGTCCGAGACGACGACGGAGCCCCGCTGGTGCGGGCTGAGCCGATCCTGACCCGTGAGCAGCTGGAGGCGCTGCGCGCCGAGCTCGTCAAGACCTCCCGG<br>GCGAAGCCCGCGGTGTCTACCCCGTCGCTGCTGCTGCGGGTGTTGTTCTGCGCGGTGTGCGGGGAGCCCGCGTACAAGTTCGCCGGGGGAGGACGTAAGCACCCGCGCTACCGCTGCCGCTC<br>GATGGGGTTCCCGAAGCACTGCGGGAACGGCACGGTGGCGATGGCCGAGTGGGACGCGTTCTGCGAGGAGCAGGTACTGGATCTGCTCGGGGACGCGGAGCGTCTGGAGAAAGTCTGGGT<br>AGCGGGCTCGGACTCCGCGGTCGAACTCGCGGAGGTGAACGCGGAGCTGGTGGACCTGACGTCGCTGATCGGCTCCCCGGCCTACCGGGCGGGCTCTCCGCAGCGAGAAGCACTGGATGCC<br>CGTATTGCGGCGCTGGCCGCGCGGCAAGAGGAGCTGGAGGGCCTGGAGGCTCGCCCGTCTGGCTGGGAGTGGCGCGAAACCGGGCAGCGGTTCGGGGACTGGTGGCGGGAGCAGGACAC<br>CGCGGCAAAGAACACCTGGCTTCGGTCGATGAACGTTCGGCTGACGTTCGACGTCCGCGGCGGGCTGACTCGCACGATCGACTTCGGGGATCTTCAGGAGTACGAGCAGCATCTCAGGCTCG<br>GCAGCGTGGTCGAACGGCTACACACCGGGATGTCGGCGGCTAACGACGAGAATTATGCATTGGCCGCCTAATAAAGGTCTAACGCATGAGAAAGCCCCCGGAAGATCACCTTCCGGGGGCTT<br>TTTTATTGCGCCCTTGAGAGCTTCCAGGATACATAGATTACCACAACTCCGAGCCC***TAGCGGCCGCTGCAGGCATGCCTCGAGATGCATGGCGCCT***AACCTAAACTGACAGGCATCAAATTAAG<br>CAGAAGGCCATCCTGACGGATGGCCTTTTTGCGTTTCGAACAATTGAAAAAACCTCGCGCCTTACCTGTTGAGTAATAGTCAAAAGCCTCCGGTCGGAGGCTTTTGACTTTCTGCTTACTGAATT<br>TCGGTGGTGCCGTTAATTAACCGGTGGGCCCTCATGATAATAATGGTTTCTTAGACGTCCGAAGTTCCTATTCTCTAGAAAGTATAGGAACTTCCATATGCCATGGGCGATGGTTCTGAGTAACA<br>GATAATAGAATGGCGATAAAGTGGCGGTAGAAATGGCGAATAATAGGTAATTATTGGCAAACAATGGCAATCTATGTCAATGATAAATAACGCATACTATTGATTTTCGGTTGTTCTTGTAGGA<br>ACTCATAATCGCTTGGTCGCTGGTTCAAGTCCAGCAAGGGCCACCAAATCATTATTTACGTATCAGGAAGAGTTAAGGGTACAAGGGGTACTCGATGGCGGTCTATATTTCATGACTAACGCCA<br>TTTCAAGGAAGAAGTGCGGTGGGTGGTTTCGAATGAG |
| Φ186 #2 | CCCTGGAGCCAAAATATCCTCAACCATATTTACTCCTTTCTAACATCACCAGGAAAAGTATAACGGATTGAAAATACATTGTTTATAAAAACAGCAGGCGCGCGGTAATGGCTGGATTGCGACA<br>CGGAGTTACTTTATAATCCGCTACCATGGCCCCTTAGCTCAGTGGTTAGAGCAGGCGACTCATAATCGCTTGGTCGCTGGTTCAAGTCCAGCAGGGGCCACCAAATTTAAGTAGTAAATACATA<br>CAGTTAGGCCACTCTCTCGAGTGGCCTTTTTGTTGCCTCGTTTTCGAGTGGCGATAGAATGGCGGTGGATTTTTTACCGCCACTTTCCTGAGGAGTGCATAGCTAGCTTCTTCGTCTGTTTCTACT<br>GGTATTGGCACAAACCTGATTCCAATTTGAGCAAGGCTATGTGCCATCTCGATACTCGTTCTTAACTCAACAGAAGATGCTTTGTGCATACAGCCCCTCGTTTATTATTTATCTCCTCAGCCAGCC<br>GCTGTGCTTTCAGTGGATTTCGGATAACAGAAAGGCCGGGAAATACCCAGCCTCGCTTTGTAACGGAGTAGAGACGAAAGTGATTGCGCCTACCCGGATATTATCGTGAGGATGCGTCATCGC<br>CATTAATTCACTGATCAGTGATAAGCTGTCAAACATGAGAATTGATCCGGCTGCCTCGCGCGTTTCGGTGATGACGGTGAAAACCTCTGACACATGCAGCTCCCGGAGACGGTCACAGCTTGTC<br>TGTAAGCGGATGCCGGGAGCAGACAAGCCCGTCAGGGCGCGTCAGCGGGTGTTGGCGGGTGTCGGGGCGCAGCCATGACCCAGTCACGTAGCGATAGCGGAGTGTATGCTGCACATGACAT<br>TAACCTATAAAAATAGGCGTATCACGAGGCCCTTTCGTCTTCAAGAATTAATTCCCAATTCCCCAGGCATCAAATAAAACGAAAGGCTCAGTCGAAAGACTGGGCCTTTCGTTTTATCTGTTGTTT<br>GTCGGTGAACGCTCTCCTGAGTAGGACAAATCCGCCGGGAGCGGATTTGAACGTTGCGAAGCAACGGCCCGGAGGGTGGCGGGCAGGACGCCCGCCATAAACTGCCAGGAATTAATTCCCCA<br>GGCATCAAATAAAACGAAAGGCTCAGTCGAAAGACTGGGCCTTTCGTTTTATCTGTTGTTTGTCGGTGAACGCTCTCCTGAGTAGGACAAATCCGCCGGGAGCGGATTTGAACGTTGCGAAGC<br>AACGGCCCGGAGGGTGGCGGGCAGGACGCCCGCCATAAACTGCCAGGAATTAATTCCCCAGGCATCAAATAAAACGAAAGGCTCAGTCGAAAGACTGGGCCTTTCGTTTTATCTGTTGTTTGT<br>CGGTGAACGCTCTCCTGAGTAGGACAAATCCGCCGGGAGCGGATTTGAACGTTGCGAAGCAACGGCCCGGAGGGTGGCGGGCAGGACGCCCGCCATAAACT***GCCAGGAATTGGGGATCGG***<br>***AATTCGAGCTCGGTACCCGGG***CTCGCATCCATTCTCAGGCTGTCTCGTCTCGTCTCGGAGAACGATCGTTGGCTGGGGGCCTCGCTTGGGTTATTGCTGGTGCCCGGCCGGGCGCAATATTCAT<br>GTTGATGATTTATTATATATCGAGTGGTGTATTTATTTATATTGTTTGCTCCGTTACCGTTATTAACTACTAGAGAAAGAGGAGAAATACTAAATGGTGAGAGCCCTGGTAGTCATCCGCCTGTCC<br>CGCGTCACCGATGCTACGACTTCACCGGAGCGTCAGCTGGAGTCTTGCCAGCAGCTCTGCGCCCAGCGCGGCTGGGACGTCGTCGGGGTAGCGGAGGATCTGGACGTCTCCGGGGCGGTCGA<br>TCCGTTCGACCGGAAGCGCAGACCGAACCTGGCCCGGTGGCTAGCGTTCGAGGAGCAACCGTTTGACGTGATCGTGGCGTACCGGGTAGATCGGTTGACCCGATCGATCCGGCATCTTCAGCA<br>GCTGGTCCACTGGGCCGAGGACCACAAGAAGCTGGTCGTCTCCGCGACCGAAGCGCACTTCGATACGACGACGCCGTTTGCGGCGGTCGTCATCGCGCTTATGGGAACGGTGGCGCAGATGG<br>AATTAGAAGCGATCAAAGAGCGGAACCGTTCGGCTGCGCATTTCAATATCCGCGCCGGGAAATACCGAGGATCCCTGCCGCCGTGGGGATACCTGCCTACGCGCGTGGACGGGGAGTGGCG<br>GCTGGTGCCGGACCCTGTGCAGCGAGAGCGCATCCTCGAGGTGTATCACCGCGTCGTCGACAACCACGAGCCGCTGCATCTGGTGGCCCACGACCTGAACCGGCGTGGTGTCCTGTCGCCGAA<br>GGACTACTTCGCGCAGCTGCAAGGCCGCGAGCCGCAGGGCCGGGAGTGGTCGGCTACCGCGCTGAAGCGATCGATGATCTCCGGGGCGATGCTCGGGTACGCGACTCTGAACGGTAAGACC<br>GTCCGAGACGACGACGGAGCCCCGCTGGTGCGGGCTGAGCCGATCCTGACCCGTGAGCAGCTGGAGGCGCTGCGCGCCGAGCTCGTCAAGACCTCCCGGGCGAAGCCCGCGGTGTCTACCC<br>CGTCGCTGCTGCTGCGGGTGTTGTTCTGCGCGGTGTGCGGGGAGCCCGCGTACAAGTTCGCCGGGGGAGGACGTAAGCACCCGCGCTACCGCTGCCGCTCGATGGGGTTCCCGAAGCACTGC<br>GGGAACGGCACGGTGGCGATGGCCGAGTGGGACGCGTTCTGCGAGGAGCAGGTACTGGATCTGCTCGGGGACGCGGAGCGTCTGGAGAAAGTCTGGGTAGCGGGCTCGGACTCCGCGGTC<br>GAACTCGCGGAGGTGAACGCGGAGCTGGTGGACCTGACGTCGCTGATCGGCTCCCCGGCCTACCGGGCGGGCTCTCCGCAGCGAGAAGCACTGGATGCCCGTATTGCGGCGCTGGCCGCGC<br>GGCAAGAGGAGCTGGAGGGCCTGGAGGCTCGCCCGTCTGGCTGGGAGTGGCGCGAAACCGGGCAGCGGTTCGGGGACTGGTGGCGGGAGCAGGACACCGCGGCAAAGAACACCTGGCTT<br>CGGTCGATGAACGTTCGGCTGACGTTCGACGTCCGCGGCGGGCTGACTCGCACGATCGACTTCGGGGATCTTCAGGAGTACGAGCAGCATCTCAGGCTCGGCAGCGTGGTCGAACGGCTACA<br>CACCGGGATGTCGGCGGCTAACGACGAGAATTATGCATTGGCCGCCTAATAAAGGTCTAACGCATGAGAAAGCCCCCGGAAGATCACCTTCCGGGGGCTTTTTTATTGCGCCCTTGAGAGCTT<br>CCAGGATACATAGATTACCACAACTCCGAGCCC***TAGCGGCCGCTGCAGGCATGCCTCGAGATGCATGGCGCCT***AACCTAAACTGACAGGCATCAAATTAAGCAGAAGGCCATCCTGACGGATG<br>GCCTTTTTGCGTTTCGAACAATTGAAAAAACCTCGCGCCTTACCTGTTGAGTAATAGTCAAAAGCCTCCGGTCGGAGGCTTTTGACTTTCTGCTTACTGAATTTCGGTGGTGCCGTTAATTAACCG<br>GTGGGCCCTCATGATAATAATGGTTTCTTAGACGTCCGAAGTTCCTATTCTCTAGAAAGTATAGGAACTTCCATATGCCATGGGCGATGGTTCTGAGTAACAGATAATAGAATGGCGATAAAGT<br>GGCGGTAGAAATGGCGAATAATAGGTAATTATTGGCAAACAATGGCAATCTATGTCAATGATAAATAACGCATACTATTGATTTTCGGTTGTTCTTGTAGGAACTCATAATCGCTTGGTCGCTG<br>GTTCAAGTCCAGCAGGGGCCACCAGATATAGCAAAGGCTGACGAGAAATCGTCAGCCTTTTTCTTTTTATATATCAGTTACTTTGCGTGCCAGTAAGCCGCTGCACGTACCCGCTGTGGGTCATA<br>CTGTTCCGCTTCAAAGCGGCGGCTTAAATTCTTAACGACTTTACCTTCGCCGGTTATCCAGATGAAGTAATCATCGGCAGGGATTTGCATCTGCGCCAGACGCGCATCTACCGCCTGCTCATCAT<br>GTGCCAGCCATTCGATATTAAAACCATCAAGGTGCGCGAGATAATCCTGACAGGCGTTATCCCGCACGCTAACCAGCGCACTAACTTGCGGTTTAACGGCAAGTTTGCTCAACGTTTCCAGGCG<br>GCGGCGCAATGCAGGCATTCCGGA |

Table S6. 2x differentiation strain with split-π protein design with 2 copies salicylate inducible Bxb1-LAA. Promoters, RBSs, coding sequences, inserted bases in T7 RNAP CDS, terminators, integrase attachment sites, ***Gibson homologies***, genomic pOSIP primer binding sites

| PLASMID | ORIGIN | RESITANCE | INSERT |
|---|---|---|---|
| **pRW-01** | Cole1 | AmpR | UNS1-PT7-B0034-sfGFP-T7T-UNSX |
| **pRW-02** | Cole1 | KanR | UNS1-PT7-B0034-sfGFP-T7T- UNSX |
| **pRW-03** | Cole1 | AmpR | UNS1-PT7-B0034-sfGFP-T7T-UNS3-pSalAM-BCD22-Bxb1LAA-T2m-UNSX |
| **pRW-04** | Cole1 | KanR | UNS1-T13m-PT7-B0034-sfGFP-T7T-UNSX |
| **pRW-05** | Cole1 | KanR | UNS1-T13m-T12m-PT7-B0034-sfGFP-T7T-UNSX |
| **pRW-06** | Cole1 | KanR | UNS1-T13m-T12m-PT7-U1d-dnaseI-T7T-UNSX |
| **pRW-07** | R6K | CmR | UNS1-p4d-BCD2-sfGFP-T2m-UNSX |
| **pRW-08** | R6K | CmR | UNS1-p4d-BCD8-mScarletI-T2m-UNSX |
| **pRW-09** | R6K | CmR | empty |
| **pRW-10** | p15a | AmpR | UNS1-pSalAM-BCD8-Bxb1LAA-T2m-UNSX |
| **pRW-11** | p15a | KanR | UNS1-pLasAM-B0034-pirN:cfaN-UNS3-B0034-cfaC:pirC-T2m-UXSX |
| **pRW-12** | p15a | TcR | UNS1-pLasAM-B0034-pirN:cfaN-UNSX |
| **pRW-13** | p15a | KanR | UNS1-pLasAM-B0034-cfaC:pirC-UNSX |

Table S7. Summary of plasmids used in this study.

**Determining inducer concentrations for differentiation experiments**

In determining the appropriate inducer concentrations for these experiments, we noted in a pilot experiment with 3 plate generations that while the behavior of 1x differentiation was minimally affected by the induction level of π-protein expression with Las-AHL (10-300 nM; Figure S7, S9), 2x differentiation using the split-π protein was quite sensitive (Figure S10, S12). Specifically we observed R6K plasmid copy number as inferred through a constitutively expressed mScarletI to be more sensitive to Las-AHL induction in the range of concentrations tested, with higher copy number than 1x differentiation observed across all concentrations. As well, increased π-protein induction appears to result in a higher effective differentiation rate for a given level of integrase induction. When differentiation is induced at a high level in media containing chloramphenicol which selects for the R6K plasmid, a decrease in endpoint cell density is observed as cells are repeatedly diluted for batch growths. This effect was enhanced for 2x differentiation at lower induction levels of differentiation, particularly for higher burden expression. While we did not follow up on the potential mechanisms underpinning these effects, these experiments informed the choice of 10 nM Las-AHL for all long-term differentiation experiments.

**Supplementary methods**

*mScarlet/sfGFP differentiation experiments*

Cells were grown from glycerol stock in 3 mL culture of M9CA glucose (Teknova M8010) with 34 µg/mL chloramphenicol, 100 µg/mL carbenicillin, and 1 µM Las-AHL. Overnight cultures were diluted 1:100 into the same media and grown ~2-3 hours to OD 0.2-0.4. To avoid cross-over of antibiotics and inducers, cells were pelleted (3500g for 10 min) before resuspending in M9CA with appropriate antibiotics (carb for differentiation, carb + chlor for differentiation with selection) to OD ~0.1. Cells at OD ~0.1 were diluted 1:10 into a total volume of 300 µL containing appropriate antibiotics (carbenicillin +/- chloramphenicol) and various inducer concentrations (salicylate, Las-AHL). Cells were grown in 96-well square-well plate (Brooks MGB096-1-2-LG-L) at 37°C with maximum-speed linear shaking in a BioTek Synergy H1m. OD700, sfGFP fluorescence (485/515 nm excitation/emission; gain 61 and 100), and mScarletI fluorescence (565/595 nm excitation/emission; gain 100) were measured at 10 minute intervals as appropriate. For long-term experiments, cells were diluted 1:50 after ~12 h growth into the same media conditions into a replicate plate.

*Flow-cytometry*

Immediately after the conclusion of a 12 h growth, cells were diluted 1:50 into 100 µL 1X PBS for analysis with flow cytometry. Samples were run on a Miltenyi Biotech MACSQuant VYB Flow Cytometer equipped with Violet 405 nm, Blue 488 nm, and

Yellow 561 nm lasers. sfGFP was measured with the 405 nm laser with 525/50 nm filter, and mScarletI with the 561 nm laser with 661/20 nm filter. 50,000 ungated events were recorded for each sample, and results were analyzed with custom Python code. Briefly, peak locations were determined from KDE fits of ungated flow data, gaussian mixture models used to assign cells to peaks, and cells within peaks were designated positive or negative for the respective fluorescent protein using a chosen threshold for peak mean. Peaks with mean log10(mScarletI) >3 were designated as *differentiated*.

#### Model implementation

*Cell growth*

For all simulations, we implemented carrying capacity limited growth of the form

$$\frac{dX_i}{dt} = \mu_i X_i \frac{K - X_{tot}}{K}, \tag{1}$$

where $\mu_i$ is the specific growth rate of cell type $X_i$, $K$ is the carrying capacity (number of cells), and $X_{tot}$ is the current total number of cells of all cell types. For simulations in Fig. 3.1, we modeled carrying capacity limited growth in a chemostat with constant dilution rate $D$ ($h^{-1}$):

$$\frac{dX_i}{dt} = \mu_i X_i \frac{K - X_{tot}}{K} - DX_i. \tag{2}$$

*Burden, differentiation, and integrase mutations*

In order to generate the models for these simulations, we first generated all possible genotypes that could be present in the simulation. For the naive case, each genomically integrated cassette of inducible T7 RNAP can be in two states:

1. Producer (P): Cassette will enable production and have associated burden.

2. Non-producer (N): Cassette will not produce T7 RNAP and has no associated burden.

For 1x naive, there are only two genotypes (P, N), while for 2x naive, there are three genotypes (PP, PN, NN). Though we could explicitly model both PN and NP, this is unnecessary. For the naive case, there is only a single type of mutation, the burden mutation, which occurs at rate $k_{MB}$ ($h^{-1}$). Because mutations require DNA replication/cell division, this rate is further dependent on the current growth rate:

$$X_P \xrightarrow{k_{M_B}\mu_P\frac{K-X_{tot}}{K}} X_N. \tag{3}$$

As the rate of the mutation is also proportional to the number of loci that could be mutated, for the general case of *n* cassettes we have that

$$X(P = i, N = n - i) \xrightarrow{ik_{MB}\mu_P\frac{K-X_{tot}}{K}} X(P = i - 1, N = n - i + 1), \tag{4}$$

where *i* is the number of producer cassettes, and *n* - *i* is the number of non-producer cassettes.

Differentiation brings two additional types of mutations, both of which act to disrupt the process of differentiation.

1. Differentiation mutation: Occurs at rate $k_{MD}$, and disrupts the capacity for an individual cassette to undergo recombination.
2. Integrase mutation: Occurs at rate $k_{MI}$, and disrupts integrase expression at one locus.

Before consideration of the integrase mutation, we can enumerate the possible genotypes for a single cassette:

1. PP : (P)rogenitor, (P)roducer. Cassette is in the un-recombined progenitor state, and would yield a producer cassette upon recombination.
2. PN : (P)rogenitor, (N)on-producer. Cassette is in the un-recombined progenitor state, and would yield a non-producer cassette upon recombination.
3. DP : (D)ifferentiated, (P)roducer. Cassette is in the recombined differentiated state, and is producing T7 RNAP.
4. DN : (D)ifferentiated, (N)on-producer. Cassette is in the recombined differentiated state, and is not producing T7 RNAP due to the burden mutation.
5. M- : (M)utated. Cassette is in the un-recombined progenitor state, but has incurred a differentiation mutation which prevents it from recombining. The Producer/Non-producer state is therefore not relevant and is neglected.

As each integrase cassette is either functional or not-functional in our model, we simply consider the number of functional integrase cassettes in our genotype. For example, the starting genotype for cells with two differentiation cassettes and two integrase cassettes would be (PPPP2). Mutation of one integrase cassette would then yield (PPPP1).

As with the burden mutation considered in the naive case, the rates of the burden and integrase mutations scale with the number of loci that could mutate, and depend on the current growth rate of the cells which could incur the mutation.

*Terminal differentiation*

In order to address the case of terminal differentiation, we explicitly modeled the number of cells divisions a differentiated cell could undergo. To illustrate this, we consider the case of a cell with two differentiation cassettes and two integrase cassettes (PPPP2). For differentiation without limited division, this would yield the genotype DPPP2. With terminal division, we include the subscript to indicate the number of divisions the cell has undergone: ($DPPP2_0$). With the general case where there is a limit of $n$ cell divisions, for $i < n$ we have

$$X_{DPPP2_i} \xrightarrow{\mu_P \frac{K - X_{tot}}{K}} 2X_{DPPP2_{i+1}}. \tag{5}$$

For $i = n$ we have

$$X_{DPPP2_i} \xrightarrow{\mu_P \frac{K - X_{tot}}{K}} \Phi. \tag{6}$$

*Production and burden*

In addition to tracking the population of producer cells, we also can track the production of an arbitrary product. To do so, we determine the production rate β ($cell^{-1}$ $h^{-1}$) specific to each strain in the simulation, and assume further that production is also dependent on the current growth rate. For simplicity, we set β = 1. In the naive case, any genotype that has $P > 0$, and in the differentiation case, any genotype that has $DP > 0$, is a producer. While in the single cassette case this is quite simple, for $n > 1$ cassettes we must address how to deal with the production rate for $P = \{0, 1, ..., n\}$ for the naive case, and $DP = \{0, 1, ..., n\}$ for the differentiation case. To do so, we will address production rate and burden simultaneously.

In the case of differentiation, we assign $\mu_N$ to be the specific growth rate of non-producer cells ($DP = 0$), $\mu_P$ to be the growth rate of cells with a single cassette producing T7 RNAP ($DP = 1$), and β to be the production rate for this genotype. Because growth rates must be non-negative, and cells have an inherent metabolic capacity, we assume each additional cassette will affect the growth rate proportionally according to

$$\mu_i = \mu_N \left(\frac{\mu_P}{\mu_N}\right)^{DP(i)} = \frac{\mu_P^{DP(i)}}{\mu_N^{DP(i)-1}}. \tag{7}$$

We similarly assume that the production rate does not linearly increase with the number of producer cassettes, but negatively correlates linearly with the growth rate according to

$$\beta_i = \beta \frac{\mu_N - \mu_i}{\mu_N - \mu_P}. \tag{8}$$

For $\mu_N = 1$ and $\mu_P = 0.5$ and two cassettes, production and growth rates would be

| $DP_i$ | $\mu_i$ | $\beta_i$ |
|---|---|---|
| 0 | 1 | 0 |
| 1 | 0.5 | 1 |
| 2 | 0.25 | 1.5 |

The naive case is treated identically with the number of *P* cassettes considered instead of *DP* with differentiation. However, because the initial genotype of cells with *n* cassettes has *n* producer cassettes, we must decide whether $\mu_P$ and β describe cells with $P = 1$ or $P = n$. We consider both cases in our stochastic simulations, where for the two cassette case "2x" indicates that $\mu_P$ and β describe the genotypes where $P = 2$ and "2x*" indicates these parameters describe the case of $P = 1$. For the later case, growth rates and production rates are determined as above. For the former case, the growth rate is calculated similarly by

$$\mu_i = \mu_N \left(\frac{\mu_P}{\mu_N}\right)^{DP(i)/n}, \tag{9}$$

where *n* is the total number of cassettes. The production rates are then determined as above.

*Differentiation rates*

We model the process of differentiation at a high level, and do not explicitly considering the underlying transfer functions describing the production of Bxb1 integrase or the rate of recombination given the number of intact and recombined cassettes. A mechanistic model would be difficult to implement in the context of these simulations, may prove prohibitively complex to implement with the relevant scales of population size, and is not necessary to capture the important features. Instead we set the tunable rate of differentiation $k_{diff}$ ($h^{-1}$) that is constant throughout a given simulation. This rate is the maximum total rate of differentiation when all differentiation cassettes are in the progenitor state, and all integrase cassettes are functional. Reducing the amount of integrase expression by mutation of an integrase cassette will reduce the rate of differentiation, and we assume differentiation rate varies linearly with the number of functional integrase cassettes. We also assume that the effective differentiation rate is not affected by growth rate. While this may not be completely accurate, decreased protein production rates when cells are growing more slowly will be counteracted by decreased dilution from cell growth and division. Therefore for the case of a single differentiation cassette we have

$$X_i \xrightarrow{k_{diff}\frac{I_i}{n_I}} X_j. \tag{10}$$

Where $I_i$ is the number of functional integrase cassettes for genotype $X_i$, $n_I$ is the total number of integrase cassettes, $PP_i = 1$, $DP_i = PN_i = DN_i = 0$, $I_i = 2$, and $DP_j = 1$, $PP_j = PN_j = DN_j = 0$, $I_j = 2$.

For numbers of cassettes ($n$) greater than one, the total differentiation rate across all cassettes is $k_{diff}$, and therefore the differentiation rate for any individual progenitor cassette is $k_{diff}/n$. Here we implicitly assume that the state of one cassette does not affect the rate of differentiation of any other cassette. Generally, the rate of transitioning from one genotype to another through differentiation is given by

$$X_i \xrightarrow{k_{\text{diff}}\frac{PP_i\,I_i}{n\;n_I}} X_j, \tag{11}$$

for the differentiation of a PP cassette, where $PP_j = PP_{i-1}$, $DP_j = DP_{i+1}$, $PN_j = PN_i$, $DN_j = DN_i$, $I_j = I_i$. Similarly, for the differentiation of a PN cassette, we have that

$$X_i \xrightarrow{\frac{PN_i\,I_i}{n\;n_I}} X_j, \tag{12}$$

where $PP_j = PP_i$, $DP_j = DP_i$, $PN_j = PN_{i-1}$, $DN_j = DN_{i+1}$, $I_j = I_i$.

*Plasmid loss, antibiotic degradation, and growth inhibition*

With the above components of the model, we also incorporated features to describe loss of the ColE1 plasmid, degradation of the antibiotic in the medium, and growth inhibition of sensitive cells. We gathered initial parameters from a study investigating the role bacterial cheating in driving population dynamics of plasmids with cooperative antibiotic resistance \cite{Yurtsev2013}. For a population of cells $X_R$ which are resistant and degrade the antibiotic, the concentration of antibiotic is described by

$$\frac{dA}{dt} = -X_R\frac{V_{max}}{V}\frac{A}{A+K_m} + D(A_{in} - A), \tag{13}$$

where $A$ (µg/mL) is the concentration of antibiotic in the medium; $V_{max}$ (µg cell$^{-1}$ h$^{-1}$) is the maximum rate of antibiotic degradation; $V$ (mL) is the volume of the culture; $K_m$ (µg/mL) is the Michaelis constant, the concentration at which the rate of antibiotic degradation is half-maximal; $A_{in}$ (µg/mL) is the concentration of antibiotic in the feed media, and $D$ (h$^{-1}$) is the dilution rate. For all simulations, we modeled batch dilutions ($D$ = 0). The rate of antibiotic degradation $V_{max}$ as experimentally determined by Yurtsev et al. was $10^6$ molecules cell$^{-1}$ s$^{-1}$ (~2.5x10$^{-6}$ µg cell$^{-1}$ h$^{-1}$) for antibiotic resistance encoded on a low copy plasmid when ampicillin was used. This rate was used as 1x antibiotic degradation in the simulations. As experimentally the antibiotic resistance was encoded on a high copy ColE1 plasmid, we also examined the case of 5x this rate.

To incorporate plasmid loss in the model, all genotypes and reactions associated with the genotypes were duplicated, and designated as either $R$ for resistant, or $S$ for sensitive. Growth rates and production rates for sensitive cells are determined as previously described for resistant cells, however for sensitive cells, growth rates are set to that of non-producers ($\mu_N$), and the production rate (β) to 0. While the growth rate of resistant cells is not affected by the concentration of antibiotic, the growth rate of sensitive cells was modeled using a Heaviside function:

$$\frac{dX_{i(S)}}{dt} = \mu_{i(S)} X_{i(S)} \frac{K - X_{tot}}{K} \mathcal{H}(\mathrm{MIC} - A), \tag{14}$$

where MIC is the minimum inhibitory concentration of the antibiotic, and for $A \geq MIC$, $\mathcal{H}(MIC - A) = 0$; and for $A < \mathrm{MIC}$, $\mathcal{H}(MIC - A) = 1$.

Finally, we modeled loss of the plasmid in the same manner as mutation, occuring at rate $k_{PL}$ (h$^{-1}$), and again dependent on the current growth rate:

$$X_{i(R)} \xrightarrow{k_{PL} \mu_{i(R)} \frac{K - X_{tot}}{K}} X_{i(S)}, \tag{15}$$

where $X_{i(R)}$ and $X_{i(S)}$ have identical genotypes apart from the presence or absence of plasmid. Though copy number of the plasmid certainly varies continuously and is not binary as we model it here, this captures the general feature and is tractable to implement.

*Plasmid mutation*

After incorporation of plasmid loss and antibiotic degradation, modeling plasmid mutation could be modeled without additional modification to the models. We modeled this by setting the antibiotic concentration and antibiotic degradation rates to 0. While the plasmid of interest is present at a high copy number, and the fraction of mutated plasmid can vary and with it the associated production and growth rates, modeling this explicitly would not be tractable in the context of our models. Because random partitioning affects can result in a high fraction of mutated plasmids from a single initial mutation in a time scale much faster than acquiring additonal mutations \cite{Halleran2019}, we make the simplifying assumption that a single mutation abolishes all expression with a single event.

*Deterministic and stochastic implementations*

In order to allow our models to be simulated either deterministically or stochastically, we implemented a manual ODE solver using Euler's method with time step 0.01 h that allowed selection of the simulation mode. For both deterministic and stochastic simulation, cell growth, production, and differentiation were modeled according to the previously described equations, and only mutation events were modeled stochastically. Because of the continuous nature of cell numbers produced using deterministic ODEs, cell numbers were rounded down for determining the number of cells which would mutate. To calculate the number of cells that would mutate in a given time step, we sampled randomly from a binomial distribution according to

$$n_{mut} = \mathcal{B}(N, kdt), \tag{16}$$

where $n_{mut}$ is the number of cells that mutate in a given time step, $N$ is the number of cells of the genotype considered for this reaction and the number of Bernoulli trials, $k$ is the rate that was determined for this specific event, $dt$ is the time step, and $kdt$ is the probability of the event occuring in the time step. The number of cells is then subtracted from the source genotype, and added to the destination genotype.

In the simulations shown, we model dilution of batch cultures in a manner similar to our experiments. For deterministic modelling, at the end of each batch growth, the cell population for each genotype is divided by the dilution factor (\$d\$) for restarting the next batch growth. The antibiotic concentration for the start of the next batch growth is calculated as

$$A_* = A/d + (1 - 1/d)A_0, \tag{17}$$

where $A_*$ is the concentration of antibiotic at the start of the next batch growth, and $A_0$ is the concentration of antibiotic in fresh media.

For dilution in stochastic simulations, the number of cells for each genotype in the next batch growth was determined by drawing from a binomial distribution according to

$$X_i^* = \mathcal{B}(X_i, 1/d).$$

( 18 )

Finally, because the metric of interest for production is the total amount of production and not the concentration of product, the amount of arbitrary product is not diluted but tracked continuously through subsequent batch growths. Production rate was assumed to vary with growth phase according to

$$\beta_* = \beta \frac{K - X_{tot}}{K},$$

( 19 )

where $\beta_*$ is the realized production rate given the current population size and carrying capacity.

*Deterministic modeling simulations*

Deterministic simulations in Figure 1 and Figure S1 were run with models generated as described above considering the case of a single cassette for all circuits, neglecting the integrase mutation, and not considering plasmid mutation or antibiotic degradation. Models used carrying capacity limited growtsh with constant dilution, and were terminated after 1000 hours of simulated time.
